# DNA barcoding reveals that injected transgenes are predominantly processed by homologous recombination in mouse zygote

**DOI:** 10.1101/603381

**Authors:** Alexander Smirnov, Anastasia Yunusova, Alexey Korablev, Irina Serova, Veniamin Fishman, Nariman Battulin

**Author notes:** Email addresses: AS; AY; AK; IS; V.F.; N.R. AS and NB are corresponding authors.

## Abstract

Mechanisms that ensure repair of double-stranded DNA breaks play a key role in the integration of foreign DNA into the genome of transgenic organisms. After pronuclear microinjection, exogenous DNA is usually found in the form of concatemer consisting of multiple co-integrated transgene copies. Here we investigated contribution of various DSB repair pathways to the concatemer formation. We injected a pool of linear DNA molecules carrying unique barcodes at both ends into mouse zygotes and obtained 10 transgenic embryos with transgene copy number ranging from 1 to 300 copies. Sequencing of the barcodes allowed us to assign relative positions to the copies in concatemers and to detect recombination events that happened during integration. Cumulative analysis of approximately 1000 integrated copies revealed that more than 80% of copies underwent recombination when their linear ends were processed by SDSA or DSBR. We also observed evidence of double Holliday junction (dHJ) formation and crossing-over during the formation of concatemers. Additionally, sequencing of indels between copies showed that at least 10% of the DNA molecules introduced into the zygote are ligated by non-homologous end joining (NHEJ). Our barcoding approach documents high activity of homologous recombination after exogenous DNA injection in mouse zygote.

## Introduction

Genetically modified organisms have become an important element of biomedical research (1), production of pharmaceutical proteins (2), and in agriculture (3). Despite the widespread use of transgenic organisms, many long-standing questions remain unanswered, especially concerning molecular mechanisms involved in DNA integration. It is known that the repair of double-stranded breaks (DSBs) plays an important role during genome editing and integration of foreign DNA into the genome. Homologous recombination (HR) and nonhomologous end-joining (NHEJ) are the two major pathways responsible for DSB repair in eukaryotic cells. The majority of random DSBs in somatic cells are repaired by NHEJ (4), while HR is necessary to resolve more specific problems such as rescuing a stalled replication fork or providing recombination in meiosis (5). Homology-based pathways involve invasion of single-stranded DNA filaments originating from DSB ends into homologous template region, resulting in formation of a D-loop and DNA synthesis. Different HR outcomes are possible, depending on the D-loop processing (6). In synthesis-dependent strand annealing (SDSA), restored DNA end is released after D-loop disruption, and anneals with the second DSB end. Break-induced replication (BIR) initiates long-range DNA synthesis in the absence of a second DSB end (replication fork collapse or telomere shortening). Double-strand break repair (DSBR) operates when the displaced strand from template anneals to the second broken DNA end. This way, both invading ends become physically linked in a double Holliday junction (dHJ) that could be resolved with or without crossing-over (6). With the development of CRISPR-based genome editing, the focus has shifted to other roles of DNA repair pathways: HR is exploited for precise genetic modifications, such as transgene knock-ins, and NHEJ is used to knock-out genes (7, 8). The obvious importance of these pathways for the field of genome editing serves as a driver for studying activity of these pathways in different cells (9, 10); understanding crosstalk between NHEJ and HR (11); and for developing new methods for the preferential activation of a particular pathway (12). Modification of the genome at the zygote stage is an advantageous method for obtaining genetically modified animals. However, in the literature there is no quantitative estimate of the activity of these DNA repair pathways in the early mammalian embryogenesis.

We decided to test the activity of various ways of processing double-stranded breaks in mouse zygotes after pronuclear microinjection of genetic constructs. In this method, exogenous DNA solution is injected inside the pronucleus that contains genetic material of the sperm or egg prior to fertilization (13). Usually, hundreds to thousands of DNA molecules are injected and processed subsequently by cellular DNA repair machinery, resulting in a stable DNA integration. Extensive work of many pioneer groups in the early years of transgenesis revealed several prominent features of pronuclear microinjection (14, 15). For instance, transgenes always integrate at one or few sites in the host genome and most of the transgene copies are prevalently arranged into head-to-tail tandemly oriented copies (concatemers) (data aggregated in Sup. Table 1). Here we investigated mechanisms of exogenous DNA molecule processing by combining the classical approach of pronuclear microinjection with transgene barcoding technique and next-generation sequencing (NGS). Barcoding strategy was initially applied to address individual cell fates among heterogeneous cell population (16, 17) and could be extended to trace transgene copies as well. Ten transgenic mouse embryos with varying amount of barcoded transgenes were analyzed with NGS to read terminal barcodes of each integrated transgene copy. Knowing initial barcode tags of the injected transgenes we could reconstitute connections of most of the transgene copies in concatemers and found various signatures of homology-based DNA repair pathways.

## Materials and methods

### Cloning of the barcoded vector library

Plasmid pcDNA3-Clover (Addgene #40259) was used as a base vector for cloning barcodes. Vector was digested with PciI, dephosphorylated and ligated with short adapter fragment, introducing SbfI and NheI recognition sites. These sites were used to insert 500 bp PCR product amplified from human genome with primers carrying barcode sequences. Human region was selected to avoid potential recombination of the transgene ends with mouse genome. Sequences of the barcoded primers were as follows: 5’ -CCTGCAGGNNCGANNGCANNTGCNNCTTGAATGACAACTAGTGCTCCAGG -3’ (Primer with Tail barcode), 5’- GCTAGCNNACTNNGATNNGGTNNCTATCCTGACCCTGCTTGGCT -3’ (Primer with Head barcode) (Sup. Fig. 1). Barcoded plasmid library was electroporated into Top10 cells, plated and extracted using GeneJET Plasmid Midiprep Kit (Thermo Fisher Scientific, USA). Number of colonies was estimated to be around 10000 individual clones.

### Generation of the transgenic embryos by pronuclear microinjection and PCR genotyping

Barcoded plasmid library was linearized with type IIS BsmBI restriction enzyme to provide incompatible 4bp overhangs (250bp distance from each barcode). Digested DNA was gel purified and eluted from the agarose gel using Diagene columns (Dia-M, Russia) according to the manufacturer’s recommendations. Eluate was additionally purified with AMPure XP magnetic beads (Beckman Coulter, USA) and finally dissolved in TE microinjection buffer (0.01 M Tris–HCl, 0.25 mM EDTA, pH 7.4). Fertilized oocytes were collected from superovulated F1 (CBA × C57BL/6) females crossed with C57BL/6 males. DNA was injected into the male pronuclei (1-2 pl ∼ 1000-2000 copies) as described elsewhere (18). Microinjected zygotes were transferred into oviducts of recipient pseudopregnant CD-1 females. Embryos were extracted at the 13.5 day of development and PCR genotyped with primers for Clover backbone or transgene-transgene junctions (same primers that we used for generating NGS PCR products) (Sup. Table 3). TAIL-PCR was performed as described earlier (19) with primers complementary to the 5’- or 3’-ends of the transgene (Sup. Table 3). Conventional PCR reactions with Taq, Q5 or LongAmp polymerases (NEB, USA) were set up to amplify various rearrangements in concatemer structure, study copy order with barcode-specific primers and validate transgene-genomic borders (Sup. Table 3).

All experiments were conducted at the Centre for Genetic Resources of Laboratory Animals at the Institute of Cytology and Genetics, SB RAS (RFMEFI61914X0005 and RFMEFI61914X0010). All experiments were performed in accordance with protocols and guidelines approved by the Animal Care and Use Committee Federal Research Centre of the Institute of Cytology and Genetics, SB RAS operating under standards set by regulations documents Federal Health Ministry (2010/708n/RF), NRC and FELASA recommendations. Experimental protocols were approved by the Bioethics Review Committee of the Institute of Cytology and Genetics.

### Copy number quantification by ddPCR

Droplet Digital PCR (ddPCR) was performed using ddPCR Supermix for Probes (No dUTP) and QX100 ddPCR Systems (Bio-Rad, USA) according to manufacturer’s recommendations. The 20 μl reaction contained 1× ddPCR Supermix, 900 nM primers, 250 nM probes and 3-60 ng digested genomic DNA. We adjusted input DNA quantity for each embryo to account for tissue mosaicism and transgene copy number variation which affected transgene:control relative dilutions in every embryo. We tested various restriction digestion conditions in order to reliable separate all transgene copies (Sup. Fig. 3) and decided to perform overnight digestions with HindIII-HF or DpnII (NEB, USA) in CutSmart buffer. Transgene copy number (Clover gene) was normalized to Emid1 control gene (Codner et al., 2016) or the unique transgene-genome border region (identified for four embryos). PCR was conducted according to the following program: 95 °C for 10 min, then 40 cycles of 95 °C for 30 s and 61 °C for 1 min, with a final step of 98 °C for 7 min and 20 °C for 30 min. All steps had a ramp rate of 2 °C per second. ddPCR was performed in two independent technical replicates. Sequences for primers and probes are available in Sup. Table 3. Data were analyzed using QuantaSoft (Bio-Rad, USA).

### DNA library preparation for NGS

NGS library 1 (original barcoded plasmids): barcoded plasmid library was digested with NheI and SbfI (Fig. 1A). 640 bp DNA fragment with a pair of barcodes was gel purified, eluted in TE buffer and sequenced.

**Figure 1.**
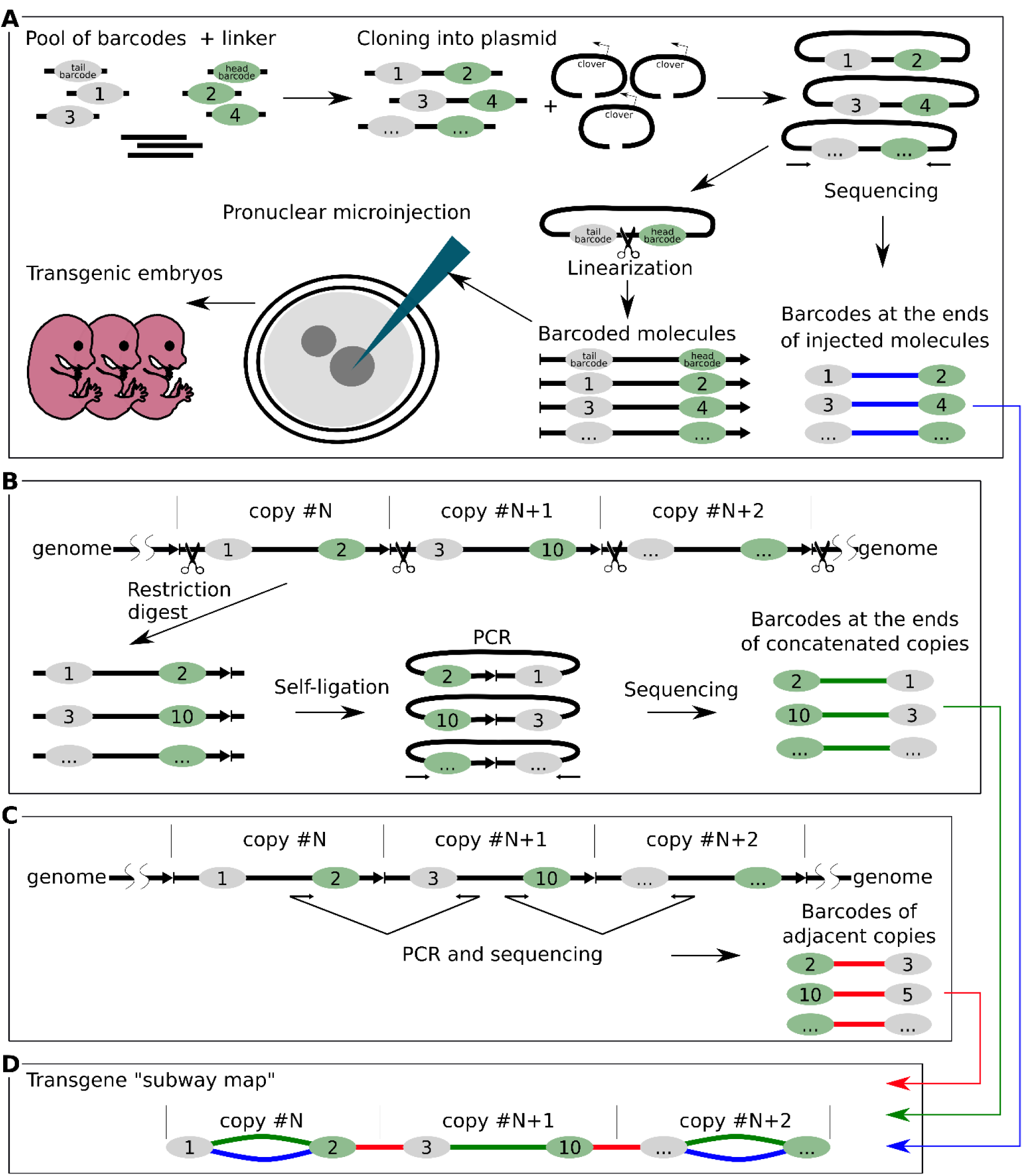
Schematic overview of the experimental design. **(A)** The main stages of generation of the barcoded plasmid library (NGS library 1) and pronuclear microinjection of barcoded molecules. **(B)** Determining barcodes at the ends of concatenated copies by inverse PCR. DNA was digested with PciI and ligated in conditions favoring self-ligation (NGS library 3). **(C)** Determining barcodes of adjacent copies (NGS library 2). **(D)** Element of a transgene «subway map» - our visualization approach that combined NGS data from three sequencing experiments. Colors indicate different barcode connections: green – actual transgene copy in concatemer from the inverse PCR data; red - transgene-transgene junctions; blue – copies that retained combinations of barcodes which were present in the injected plasmids.

NGS library 2 (PCR of the barcoded transgene-transgene junctions): Concatemer junctions containing barcodes were PCR amplified using Q5 polymerase (Fig. 1C). PCR conditions were tested to avoid accumulation of PCR artifacts in late cycles and to account for copy number variation between embryos (range of 0.2 copies to 300 copies). PCR program: 98°C for 30 s, then 25-30 cycles of 98°C for 15 s, 64°C for 30 s, 72°C for 1 min, with 3 min of final extension at 72°C. 25 µl of PCR reaction were purified using AMPure XP magnetic beads (Beckman Coulter, USA), eluted in TE buffer and sequenced. In order to obtain additional information about transgene-transgene junctions we also sought to sequence PCR products corresponding to the internal junctions (100 bp from BsmBI cut site) in three multicopy lines (embryos #2, 3, 7). To stay in the range of acceptable read length (∼150 bp) we used one primer close to the junction and another one flanking barcode (primer pairs 2+14 and 1+13 at Sup. Fig. 6). PCR products were generated for both orientations. This way, a unique signature of the trimmed junction could be assigned to specific barcodes (Fig. 6).

NGS library 3 (inverse PCR of the barcoded transgene ends): Genomic DNA from transgenic embryos was digested overnight with excess of PciI enzyme that cuts 87 bp away from one of the barcodes, inside the junction (Fig. 1B). Digested DNA was purified with AMPure XP magnetic beads. To avoid incomplete digestion, sticky ends generated with PciI were filled-in with Klenow enzyme to produce blunt ends. This signature was later used for filtering during sequence analysis. After heat inactivation of Klenow enzyme (20 min at 75°C), 300 ng of digested DNA were ligated overnight at 16°C in a large reaction volume (100 µl) to facilitate self-ligation of transgene monomers. Terminal barcodes of self-ligated DNA fragments were PCR amplified with Q5 polymerase with the same primers and conditions as were used for NGS library 2 generation. To remove PCR fragments originating from undigested DNA, Illumina prepared adapter-ligated PCR products were additionally treated with PciI, gel purified and used for sequencing. We have prepared 10 libraries corresponding to 10 embryos and one control library representing a 1:1 mix of self-ligated DNA from embryos #1 and #4 under the same conditions (150 ng + 150 ng of digested genomic DNA in 100 µl ligation). Control library served as a quantitative measurement of random transgene ligation as opposed to self-ligation.

DNA fragments from three libraries were prepared with NEBNext Ultra DNA Library Prep Kit for Illumina (NEB, USA), pooled together and sequenced on the Illumina HiSeq 2500 platform (Illumina, USA). Libraries were assessed using an Agilent 2100 bioanalyzer (Agilent, USA) and a Qubit dsDNA HS assay kit (Life Technologies, USA).

All of the above experiments were performed using Q5 polymerase (NEB) to avoid PCR artifacts (barcode mutations or switching), as it was shown that other polymerases have much higher rate of strand conversion (20).

### Computational data analysis

#### Overview of data processing pipeline

NGS data processing contained four steps. First, reads were trimmed using cutadapt to remove constant sequences flanking barcodes. Second, read pairs were searched for complete or partial match of barcode pattern (NN CGA NN GCA NN TGC NN for tail barcode, NN ACT NN GAT NN GGT NN for head barcode) and pairs sharing identical barcodes were merged. This results in initial set of barcode pairs, which was further filtered at the third step of the pipeline to produce final set of pairs. These resulting pairs were visualized using *Network* module of the *vis.js* framework (http://visjs.org/). All computations were performed using nodes of Novosibirsk State University high-throughput computational cluster.

### Reads trimming

We used cutadapt v. 1.15 (21) to perform reads trimming. For NGS Library 1 (initial plasmid library) we ran cutadapt independently on each of two mates from the read pair, and then combined obtained results. For NGS Library 2 (Junction PCR) we ran cutadapt in paired-end mode using –G option, which allowed us to distinguish PCR products containing barcodes and PCR products containing internal junction sequences. Each read pair may have one of two orientations. In the Forward-Reverse orientation, first mate corresponds to the 5’-end of the PCR product and the second mate corresponds to the 3’-end. In the Reverse-Forward orientation first mate corresponds to the 3’-end and second mate to the 5’-end. To account for both orientations, we ran cutadapt on the input data twice, each time searching for one of these two scenarios (see Sup. Table 4), and combined obtained results. For NGS Library 3 (Inverse PCR), we first ran cutadapt with adapter sequences, which include inverse PCR ligation junctions (cutadapt’s first pass in the Sup. Table 4). We considered reads where inverse PCR ligation junction was present as inverse PCR products, whereas reads with intact site of PciI enzyme were discarded, as they represented junction between two adjacent transgene molecules. Since PciI site is located far from barcode sequence, we ran cutadapt again on the reads, which, according to the first pass, contained inverse PCR ligation junctions (second pass in the Supplementary Table 2). During second pass, reads were trimmed immediately before barcode sequence, in a same manner as we did for Library 1. Both first and second passes of cutadapt were executed independently for each of two read mates, and obtained results were combined together. Details of cutadapt run parameters are presented in Supplementary Table 2.

### Barcode identification and reads merging

We used custom python script to analyze cutadapt output and count identical read pairs. At this stage, we discarded reads which were too short (read length after trimming < 18 bp) and reads with too low quality. As a quality threshold, we required that within 21 base pairs at the 5’-end of the read, which presumably represent barcode sequence, all base calls had mapQ values above 20.

Resulting dataset of read pairs was further analyzed using python regular expression module to find barcode sequences within each read. Although all barcodes were supposed to share similar pattern (NN CGA NN GCA NN TGC or NN ACT NN GAT NN GGT NN), we expected that some of them might be slightly different due to errors, which may occur during oligonucleotide synthesis. Thus, we composed a set of regular expressions covering both exact and partial matches of barcode pattern (Sup. Table 5). Here, it is pertinent to note, that vast majority of identified barcodes (∼96%) perfectly matched expected pattern. Once barcode sequences were identified in each read, we collapsed those read pairs, which have identical barcode sequence followed by same or different 3’-endings. After the merging, we obtained dataset of pairs hereinafter referred as *barcode pairs*, each characterized by 5’- and 3’-barcode sequences and number of reads, supporting the pair.

### Pairs filtering

PCR amplification may introduce mutations which can be interpreted as “novel” barcodes. One can suppose that original sequences will be supported by higher number of reads than their counterparts containing mutations introduced by PCR. However, it is not clear how to select the read count threshold reflecting this difference between correct and erroneous sequences. To solve this problem, we employed the fact that correct barcode sequences should be shared between embryo data (NGS Libraries 2 and 3) and initial plasmid library (NGS Library 1). For each data, we gradually increased read count threshold and monitored increase of the overlap between embryo’s and plasmid barcode sequences, and selected the point when overlap size jumps. An example of such jump for the embryo #3 data is shown in Sup. Fig. 21. Note that for some embryos these approach was not efficient, as overlap size gradually increases and there was no defined jumping point. Thus, we always performed additional visual inspection of read counts distribution to define optimal threshold. For Library 1 (initial plasmid library) analyses, we used relaxed threshold of 8 reads.

After read counts filter, we removed all barcodes identified in embryo data, which were not found in original barcoded plasmid data. At this filtering step, we made exception for those barcodes that were within top 5% by read count in current dataset.

Next, we applied empirical thresholds for pair counts. As barcode pairs maybe shuffled by the cellular DNA repair machinery, and thus pairs observed in embryo may not correspond to pairs determined in initial plasmid library, we could not apply same strategy to find threshold count as we did for barcodes. Thus, we manually selected thresholds for each sample based on read counts distribution (provided in Sup. Table 6). Moreover, if barcode was observed in several pairs in one embryo, we discarded pairs accounting for less than 1% of all paired reads containing this barcode sequence. When processing initial plasmid library, we raised this threshold to 5%.

### Availability of data and materials

The sequencing results will be available in the NCBI GEO database upon article acceptance. Processed data (transgene “subway maps” in web-browser file format) are available in supplementary (Sup. Files 1-10) and at http://icg.nsc.ru/ontogen/

## Results

### Generation of the barcoded concatemers by pronuclear microinjection

We decided to replicate conditions of a typical microinjection experiment. In order to investigate the mechanisms leading to formation of concatemers we injected zygotes with a library of transgene molecules labeled with unique DNA barcodes. The strategy for introducing barcodes is shown in Figure 1A. A 7 Kb plasmid vector expressing Clover was tagged with two barcodes, 280 bp from the future ends (Sup. Fig. 1). We sought to find compromise between the length of the DNA ends precluding barcodes, as longer fragments would preserve barcodes from exonuclease trimming, while shorter ends are more suitable for generating PCR products for NGS sequencing (about 700 bp in our case). We sequenced the DNA of the barcodes in the plasmids and estimated that our library consists of 12,657 different molecules (Fig. 1A). This barcoded plasmid library was linearized with BsmBI which cuts between two barcodes to generate incompatible 4 nt 5’-overhangs. Linear DNA was subsequently injected into pronuclei following standard protocol (each zygote received around 1000-2000 DNA molecules) and zygotes were transferred into pseudopregnant foster mothers. Embryos were collected at the day E13.5 of development and their DNA analyzed by PCR genotyping (Sup. Fig. 2). Out of 50 embryos, 10 turned out to be positive for the transgene integration (20%), constituting a normal outcome for this method.

### Determination of the transgene copy number

First, we quantified transgene copy number using droplet digital PCR (ddPCR). A pair of probes was designed for multiplex ddPCR: (1) transgene-specific probe for the Clover gene in the middle of the vector, and (2) standard reference probe for the gene Emid1 at chromosome 11 as control (tested in (22)). As seen in Sup. Fig. 4, transgene copy numbers varied greatly between embryos. In some cases, this number was less than one copy due to mosaicism of the embryo tissues (embryos #5 and #6) which resulted in dilution of the transgenic alleles with wild-type alleles. Fortunately, we managed to obtain genomic localization information for some embryos, using TAIL-PCR. Transgene-genomic border is a unique site that could be used as a probe target region for ddPCR to implement mosaicism correction (Sup. Fig. 3). Thus, replacing the standard reference Emid1 gene with a transgene-genome border specific probe allowed us to clarify copy number for four of the embryos (Sup. Fig. 4). For example, embryo #4 had 23 copies corrected to 45 (roughly 50% mosaicism), embryo #5 - 0.4 to 1, embryo #9 - 5 to 22, embryo #10 - 3.5 to 4. In summary, we obtained 10 transgenic embryos with a broad distribution of transgene copy numbers, ranging from 1 to ∼300 copies, as expected in typical pronuclear microinjection experiments (23).

### Next-generation sequencing of DNA barcodes in the concatemers

In order to understand internal structure and origin of concatemers, we sequenced barcodes at the ends of the individual transgene copies. We performed two alternative sequencing experiments. First, we applied the inverse PCR method to determine the head and tail barcodes for each of the molecule in concatemer (Fig. 1B). This was important, considering transgene recombination that may take place prior to integration. Genomic DNA from transgenic embryos was digested with PciI endonuclease that makes a single cut inside transgene-transgene junctions (Fig. 1B). Ligation was performed in a highly diluted solution of digested DNA. Such conditions favor self-ligation of individual transgenes - as a result, terminal barcodes come close at a distance of about 700 bp, and they can be PCR amplified and sequenced using paired-end NGS. DNA sequencing of the inverse PCR library made it possible to establish genuine pairs of barcodes at the ends of each transgene that constitute concatemers. Additionally, we directly PCR amplified and sequenced barcodes at the transgene-transgene head-to-tail junctions to get information about relative positions of molecules in concatemers (barcodes of adjacent copies) (Fig. 1C).

Combining NGS information from all sequencing experiments (initial plasmid library + inverse PCR + junction PCR) allowed us to collect comprehensive data on which transgene molecules were injected into pronuclei, how each molecule changed during end processing, and their relative positions inside concatemers. We visualized the structure of concatemers as a graph, with nodes representing individual barcodes and edges corresponding to connections (based on the NGS data for each of the transgenic embryos) (Fig. 2A) (Sup. Fig. 7–16) (Sup. Files 1-10). Figure 2A illustrates organization of all connections between barcodes of the embryo #9, taken as an example. For clarity, each type of barcode connection was assigned one of three colors. Green color – connection between two barcodes located on different ends of individual molecule within concatemer (based on the inverse PCR data); red color - shorter connections that correspond to the transgene-transgene junctions; blue color – copies that retained combinations of barcodes that were present in the injected plasmid. We named these colored schemes “transgene subway maps”. Counting of unique barcode connections in 10 embryos revealed more than 1000 individual copies of barcoded transgenes (summed up in Table 1 and Sup. Fig. 4).

**Figure 2.**
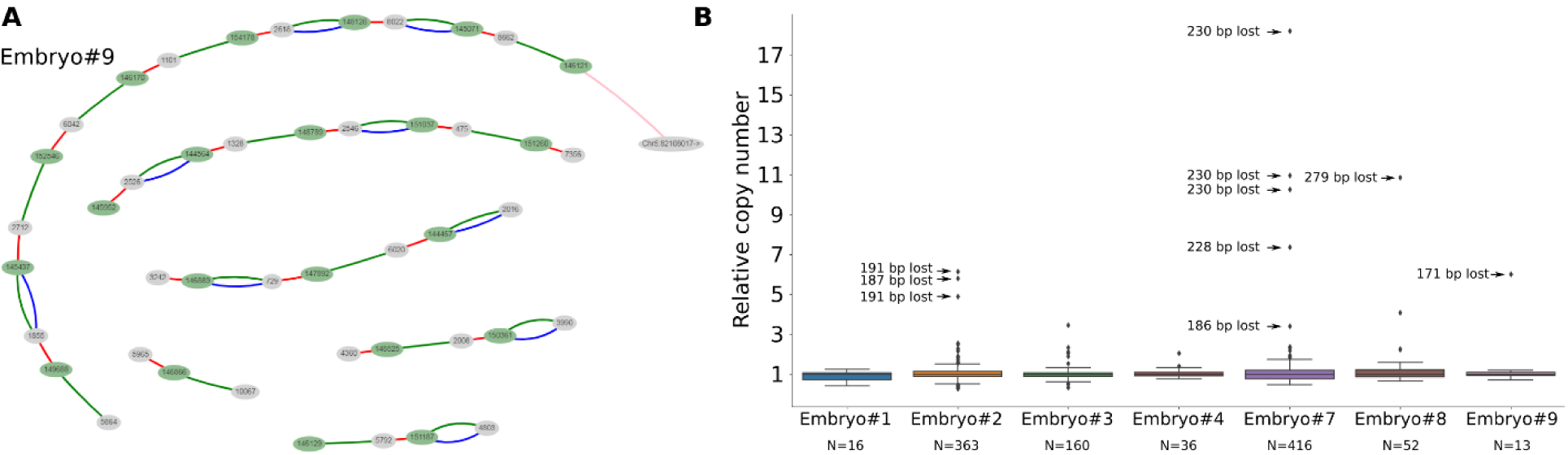
Concatemer structure. **(A)** Transgene “subway map” for embryo #9.Transgenes are oriented in head-to-tail fashion: green and gray colored ellipses designate head and tail barcodes respectively. Gaps between transgene chains are either due to deletions or alternative junction orientations. **(B)** Box plots represent the distributions of relative copy number for each terminal barcode combinations (green connections) in transgenic embryos. Relative copy numbers were calculated as read counts divided by median. Most of the outliers (relative copy number >1) were tied to deletions that create shorter PCR product. N indicates copy number (green connections) analyzed for each embryo.

**Table 1.**
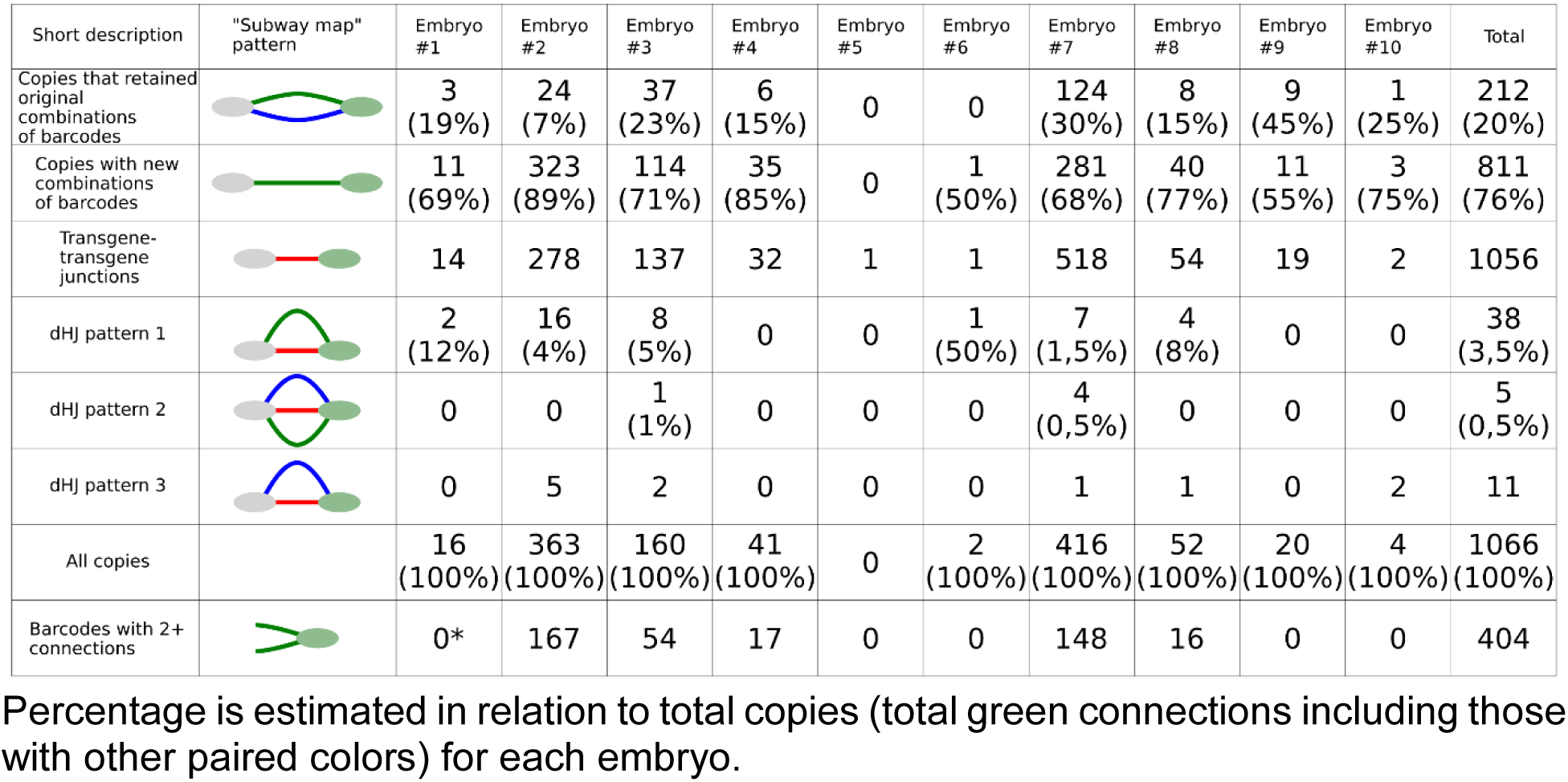
Frequencies of various connection patterns in transgenic embryos.

| Short description | "Subway map" pattern | Embryo #1 | Embryo #2 | Embryo #3 | Embryo #4 | Embryo #5 | Embryo #6 | Embryo #7 | Embryo #8 | Embryo #9 | Embryo #10 | Total |
| --- | --- | --- | --- | --- | --- | --- | --- | --- | --- | --- | --- | --- |
| Copies that retained original combinations of barcodes |  | 3<br>(19%) | 24<br>(7%) | 37<br>(23%) | 6<br>(15%) | 0 | 0 | 124<br>(30%) | 8<br>(15%) | 9<br>(45%) | 1<br>(25%) | 212<br>(20%) |
| Copies with new combinations of barcodes |  | 11<br>(69%) | 323<br>(89%) | 114<br>(71%) | 35<br>(85%) | 0 | 1<br>(50%) | 281<br>(68%) | 40<br>(77%) | 11<br>(55%) | 3<br>(75%) | 811<br>(76%) |
| Transgene-transgene junctions |  | 14 | 278 | 137 | 32 | 1 | 1 | 518 | 54 | 19 | 2 | 1056 |
| dHJ pattern 1 |  | 2<br>(12%) | 16<br>(4%) | 8<br>(5%) | 0 | 0 | 1<br>(50%) | 7<br>(1,5%) | 4<br>(8%) | 0 | 0 | 38<br>(3,5%) |
| dHJ pattern 2 |  | 0 | 0 | 1<br>(1%) | 0 | 0 | 0 | 4<br>(0,5%) | 0 | 0 | 0 | 5<br>(0,5%) |
| dHJ pattern 3 |  | 0 | 5 | 2 | 0 | 0 | 0 | 1 | 1 | 0 | 2 | 11 |
| All copies |  | 16<br>(100%) | 363<br>(100%) | 160<br>(100%) | 41<br>(100%) | 0 | 2<br>(100%) | 416<br>(100%) | 52<br>(100%) | 20<br>(100%) | 4<br>(100%) | 1066<br>(100%) |
| Barcodes with 2+ connections |  | 0* | 167 | 54 | 17 | 0 | 0 | 148 | 16 | 0 | 0 | 404 |

Percentage is estimated in relation to total copies (total green connections including those with other paired colors) for each embryo.

Amongst noteworthy map features are discontinuities in connection chains (embryo #1 and #9 are prominent examples) (Sup. Fig. 7, 15). The gaps obviously indicate lack of PCR product connecting barcodes in our sequence reads. This could happen for two reasons. First, transgene-genome borders are not subjects for PCR with our NGS primers, thus at least two detached barcodes are ensured for any map. In addition, we cannot exclude multiple integration events that will increase number of connection gaps. Partial transgene deletions and inversions are another source of discontinuity: most of the gaps are certainly caused by NGS primer site loss at the concatemer junctions. As expected, we detected many transgene deletions and complex rearrangements with conventional PCR and TAIL-PCR. Number of transgene rearrangements correlated with copy number for each embryo. Embryos #2, 3, 7, 8 have hundreds of copies and plethora of rearrangements (Sup. Fig. 5). On the contrary, remaining embryos had few (#4, 6, 9) or zero (#1, 5, 10) abnormal transgene junctions. The list of some sequenced deletions and rearrangements is available in Supplementary material next to the corresponding concatemer maps. Some interesting cases (embryos #9 and #10) are highlighted in Discussion section (Sup. Fig. 15, 16).

Next, we inspected our NGS sequence data presented in a form of transgene “subway map” to understand molecular mechanisms that lead to concatemer emergence.

### *De novo* amplification does not contribute to concatemer formation

One of the motivations for our work was to test the hypothesis that rolling circle replication or analogs can take part in the formation of a tandem of head-to-tail oriented copies (24). This mechanism is used by some viruses of eukaryotes to amplify their genome (25), and is suspected to participate in telomere maintenance (t-circles) (26) and in yeast mitochondria replication (27). It is established that after microinjection some DNA copies can be circularized by NHEJ (28). Such circular molecule could probably undergo rolling circle replication with the involvement of a BIR-like mechanism and strand displacement, for example. Conceptually, this mechanism can be a good candidate for the role of the constructor of tandemly oriented concatemers. However, in our data we did not find any evidence supporting this hypothesis, since the number of unique barcoded molecules found in concatemers was in good agreement with the estimates obtained by the ddPCR method (Sup. Fig. 4), even in multicopy embryos (70-300 copies). Nevertheless, in order to assess whether individual transgenes were amplified, we analyzed distribution of the sequence read counts for each of the unique transgene copies (green connections). Although we found several transgenes that had increased read counts, in most of these cases the shift was explained by deletions in the transgene-transgene junction regions, resulting in shorter PCR products which altered PCR kinetics. Still, there are several copies for which this technical explanation does not work. Apparently, these are genuine examples of molecules that have doubled their copy number (embryo #3 (Fig. 2B)). Although our data do not allow us to suggest a non-contradictory mechanism for this phenomenon, it is worth noting that in our analysis there were more than 1000 molecules in concatemers, of which only around 10 could be suspected of amplification. Thus, we can conclude that the concatemers are formed by direct linkage of injected molecules, rather than by de novo amplification mechanism.

### HR is essential for concatemer formation

Our transgene “subway maps” illustrate high recombination activity that assembles transgenes into concatemers, including barcode “switching” (green connections without paired blue connections) and “branching” of the barcode nodes in the multicopy lines. We identified several typical connection patterns in the transgene “subway maps” and proposed which of the known DNA repair pathways could led to their formation. Of these, we examined NHEJ and two sub pathways of HR - SDSA and DSBR (Sup. Fig. 17). First, it is important to note that identical linear copies of DNA cannot be combined by HR mechanism without template region bridging two copies. Therefore, the formation of any concatemer undoubtedly begins with non-homologous end joining. However, aside from initial ligation, NHEJ plays only a minor role in assembling concatemers (see estimates in next chapters). For example, as seen in Figure 2A, in embryo #9 only 9 out of 20 copies preserved initial combination of barcodes that were observed in the injected plasmid library (coinciding blue and green connections) while the other 11 contain the head barcode from one molecule, and the tail one from the other (therefore they do not have blue connection). Such an exchange of genetic information between molecules is a characteristic signature of homologous recombination and strikingly differs from the simple combination of intact (with the exception of small indels at the junction) molecules produced by NHEJ mechanism. In our total sample of 1066 copies, only 20% (212) retained the original combination of barcodes (Table 1). Thus, it can be concluded that at least 80% of the molecules in the concatemers were processed by the homologous recombination mechanism. Most likely, this is an underestimation, because in our experimental system barcodes are located almost 300 bp from the ends and resection might not always reach barcode sequence to change it through recombination (Sup. Fig. 17).

### Recombination mechanisms devised from connection patterns

To make comprehensive analysis of the connection patterns, it is important to discuss how the structures could be formed. We planned to use terminal barcodes as mere informative tags for concatenation analysis, but their location at the ends unexpectedly turned them into an indicator of recombination activity. Figure 4 shows possible mechanisms explaining the formation of recombined copies. In DSBR, after 3’-end resection and homologous duplex invasion, the D-loop synthesis reaches the barcode and forms a mismatch on one of the strands. This mismatch is a substrate for the mismatch repair system, which removes the fragment of the strand containing the mismatch and completes the gap on the template of the remaining strand. It is known that during recombination strand discrimination removes information from the invading strand rather than from the repair template, resulting in gene conversion (29, 30). Thus, mismatch repair leads to the copying of the donor barcode into the invading transgene. Simply put, the exchange of barcodes between the copies is a typical example of gene conversion. Interestingly, our assumption about the active participation of the mismatch repair system was confirmed by the analysis of embryo #1. Here we found two chains of transgenes consisting of 3 and 11 copies. As can be seen in Fig. 3A, some barcode connections form an unusual structure with a fork at one end (bottom part of the map). We assumed that the embryo #1 is a mosaic, whose cells contain one of two barcodes in a given position of concatemer in equal proportions. This is possible if, for some reason, the mismatch that was formed during recombination was not repaired before replication and daughter cells received two different barcode variants (Fig. 3B) (Sup. Fig. 18). These separated barcodes have about half the number of reads than the other copies in concatemer (Fig. 3C). We did not observe this pattern in any other embryo, so it must represent a unique event.

**Figure 3.**
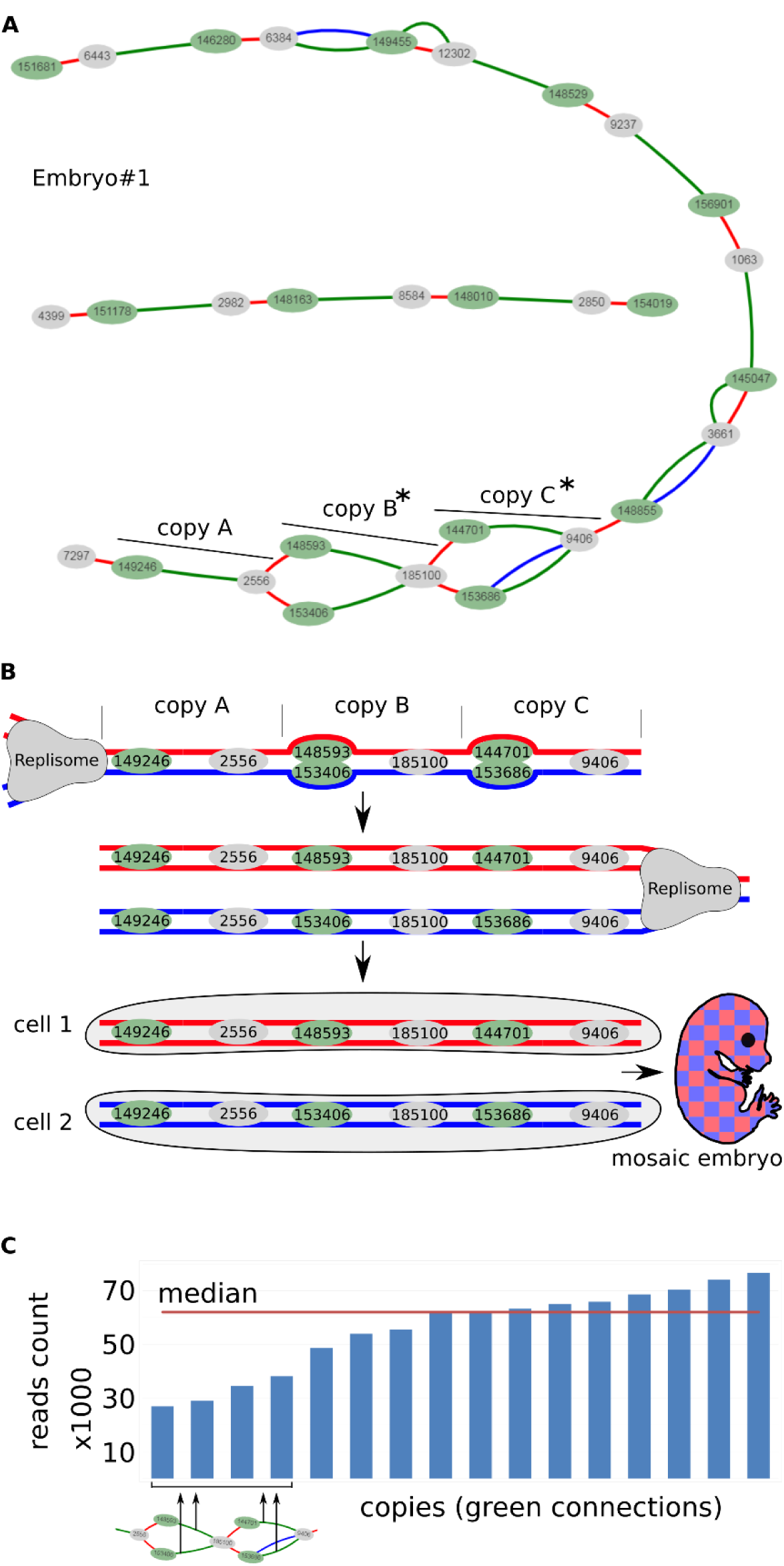
**(A)** Transgene «subway map» for embryo #1. Copies B and C have two alternative head barcodes. **(B)** The scheme explaining the emergence of mosaic embryo consisting of two cell populations in case the barcode mismatches were not repaired in time before DNA replication (more details in Sup. Fig. 18). **(C)** Copies B and C have roughly half as many reads as the other copies in the embryo #1.

Copying barcodes from a donor template explains why in many cases concatemer maps look like a web of branched nodes (embryos #2, #3, #4, #7, #8) (Sup. Fig. 8, 9, 10, 13, 14). Whenever the junctions that were originally linked together by HR or NHEJ serve as a donor template, they can share their barcode (one or both) with invading copy causing it to switch barcodes (Fig. 4). This way, one barcode will be connected with two partners at independent junctions even if these regions lie at distant positions in concatemer. Abundance of these nodes demonstrates intensive HR process that sometimes copies one junction 3-5 times with different invading ends (404 counts overall, Table 1), but also greatly complicates the “subway map” for visual inspection. Note that, as stated earlier, most of the original transgenes (blue connections) fail to be incorporated into the concatemers even if they provide recombination templates for other copies.

**Figure 4.**
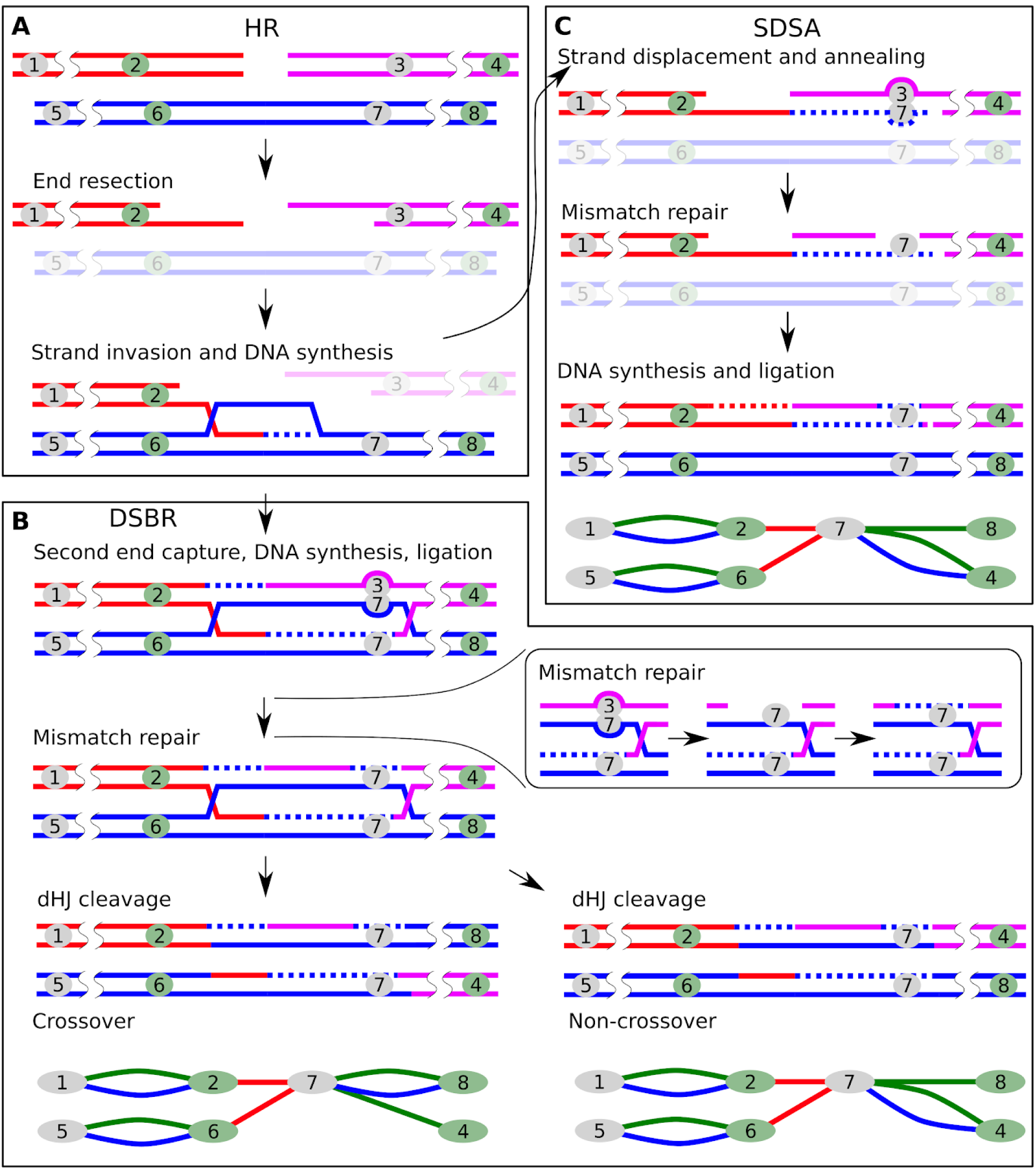
Principle of recombination between transgene copies causing barcode “switching”. Stages common to all pathways of homologous recombination **(A)** and stages characteristic of DSBR **(B)** and SDSA **(C).** The numbers denote barcodes. Outcomes of recombination are shown as elements of the transgene “subway map”. The mismatch repair steps are shown in the box.

### Evidence of dHJ formation

In the schemes described above, SDSA products are indistinguishable from non-crossover DSBR products (Fig. 4 and Sup. Fig. 17). However, we found three connection patterns that strongly support the fact of dHJ resolution with crossing-over. <u>dHJ pattern 1:</u> If both ends of a single transgene molecule invade one junction, dHJ is formed and can be resolved with the formation of crossover products. This leads to the integration of the “attacking” copy between the two original ones, and both of the “attacking” copy’s barcodes are overwritten by those in the junction (Fig. 5A). On our concatemer maps, this is represented by paired green and red connections. For example, there are two such connections in the embryo #1 (Sup. Fig. 7) and 32 in total (Table 1). Interestingly, if dHJ is resolved without formation of crossover products, then the attacking molecule becomes circularized. Such circles are either lost during cell division or serve as templates for other linear transgene copies.

**Figure 5.**
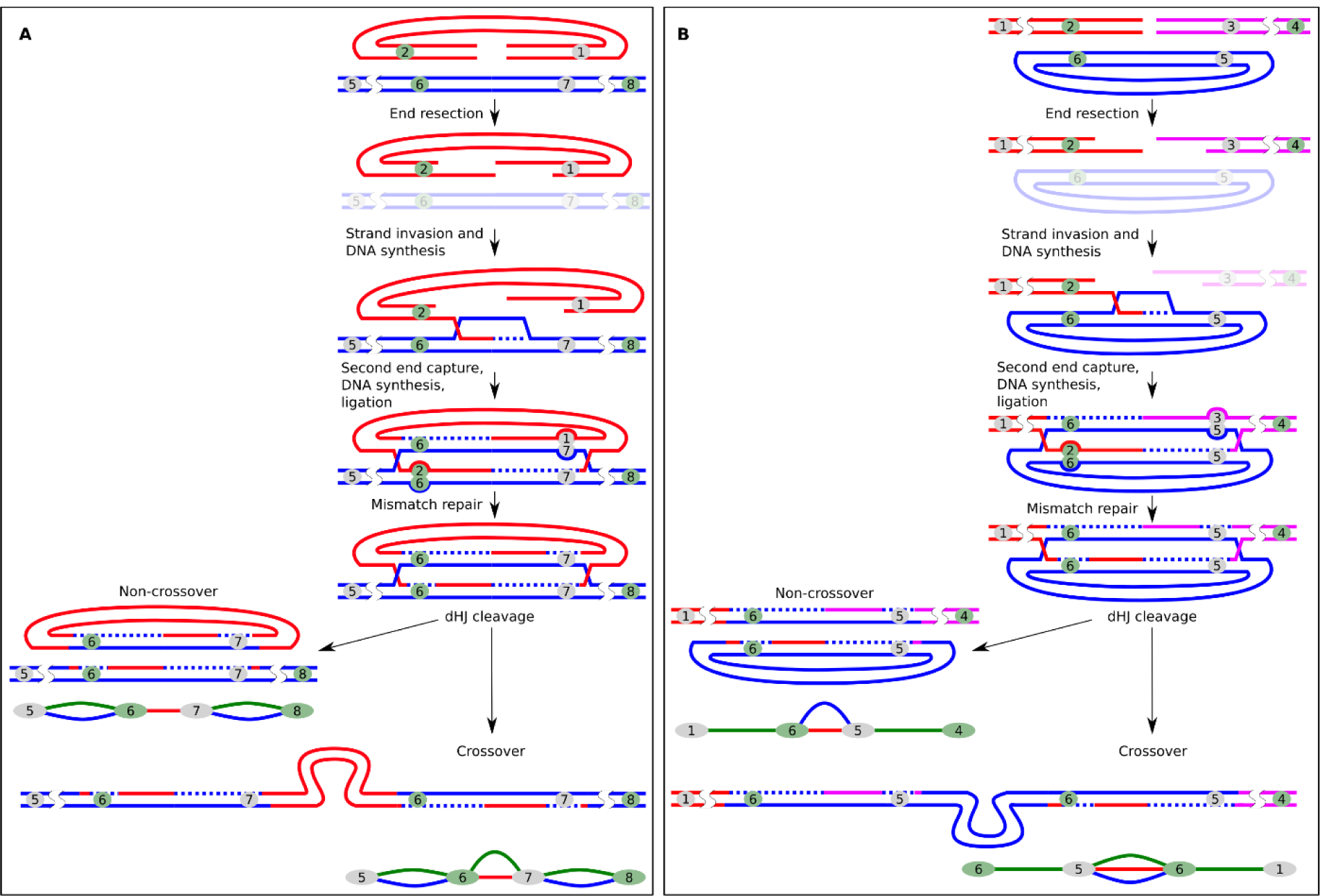
Resolution of dHJs during DSBR leads to characteristic connection patterns. **(A)** Crossing-over between copies could result in assimilation of another copy (“attacker”) into junction with loss of the “attacker”’s barcodes (green + red pattern). **(B)** DSBR between linear ends and a circular copy can have two detectable outcomes: circular copy donates barcodes without crossing-over (red + blue pattern) or gets incorporated into “attacking” molecule while also donating barcodes (green + red + blue pattern). Outcomes of recombination are shown as elements of the transgene “subway map”.

<u>dHJ pattern 2:</u> Another crossover scenario occurs when a single circular copy is attacked by the ends of two other molecules (Fig. 5B). In this case, dHJ resolution with the formation of crossover products leads to the joining of all three copies, while the “attacking” molecules copy barcodes from the circular template. This outcome appears as two barcodes linked by three connections at once. There are five such structures in our maps (Table 1). In addition, even if the “attacking” molecules copy barcodes from the template without crossing-over and physical integration of the circular copy, we would still see such events (Fig. 5B). On our maps, such barcodes are connected by red and blue connections (<u>dHJ pattern 3</u>). This pattern is characteristic of a circularized copy as well. However, these are only a few (11 of 1066 total molecules in our analysis), which says that the closure of a single molecule in a ring is a rare event (Table 1). This is important, because according to the initial theories, circularization and subsequent random breakage were considered one of the key stages of the formation of concatemers (28).

We would like to emphasize that although we found only sparse evidence of crossing-over (5% of the copies) (dHJ patterns in Table 1), all of these were simple, categorizable cases which are just a tip of the iceberg, as many simultaneous recombination events must have created higher order patterns. For instance, crossovers could be formed by multicopy tandems that are incorporated into junctions (this would result in a side loop on transgene “subway map”). As many of the individual transgenes also switch barcodes by junction invasions, this side loop would be connected to multiple nodes in other concatemer regions, thus vanishing in the complex “subway map” (like in embryos #2, 3, 4, 7, 8) (Sup. Fig. 8, 9, 10, 13, 14).

### Role of NHEJ in concatemer formation

Our data suggests that HR plays a leading role in the formation of tandemly oriented copies, but as noted earlier, initially HR requires template junctions created by NHEJ. Since the typical signatures of NHEJ are small indels at the repair sites (4), we decided to explore the repertoire of indels at the transgene-transgene junctions in multicopy embryos (#2, 3, 7). In total, we analyzed the sequences of junctions adjacent to 1803 barcodes. We found almost a hundred possible variants for the structure of ligation sites between copies (Fig. 6A). However, the frequency of sequence variants was distributed very unevenly, so that the three most frequent variants were found in 80% of the junctions (Fig. 6B). These top 3 variants were the same in all examined embryos. These variants were clippings of the protruding 4 nt 5’-ends: -5 nt (Var1), -5 nt (Var2), -7 nt (Var3). Remarkably, other deletions of the same or even smaller size were rare (Fig. 6A). The fact that identical indels appeared in three independently injected zygotes confirms that NHEJ has preferred ligation products. In our case, processing of the 5’-overhangs might have revealed complementary nucleotides (GA in Var1 and AG in Var3), hence favoring ligation of these variants over others. Recent rigorous analysis of NHEJ patterns in mouse ES cells (31) demonstrated that 5’-protruding ends are repaired by either NHEJ or TMEJ (polymerase theta-mediated end-joining). In our case, sequence variants Var1-3 did not have any additional insertions or SNPs and formed uniform clusters, thus TMEJ activity could be excluded (Fig. 6B).

**Figure 6.**
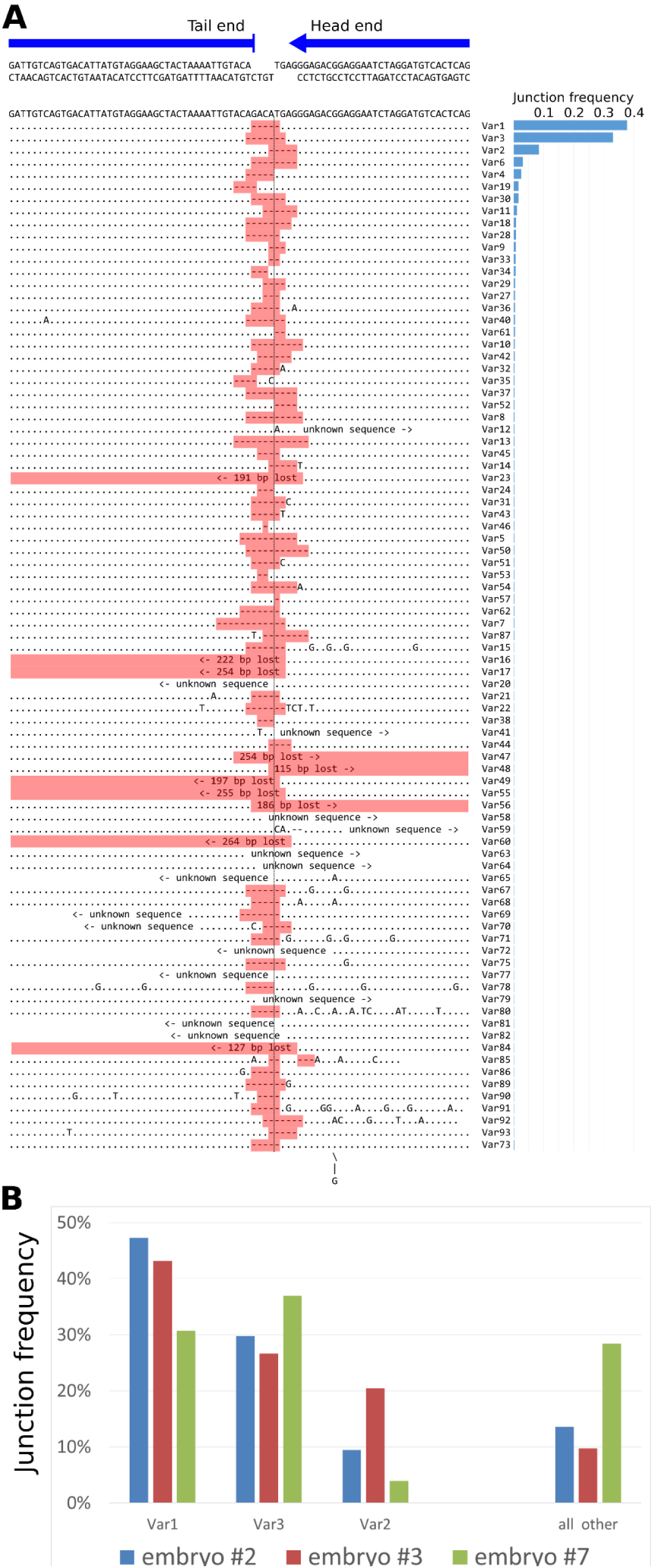
Sequence variants of indels at the junctions. **(A)** 4 nt 5’-protruding ends were generated by BsmBI digestion prior to microinjection. Sequence variants of the transgene-transgene head-to-tail junction region (Var1-Var93) are aligned below the original sequence. In most of the junctions, end processing removed only 5’-overhangs. **(B)** We calculated junction variant frequency for all detected mutations. Diagram shows distribution of the top three variants (Var1, 2, 3) and remaining variants (“all other”) in analyzed embryos.

Unfortunately, the fact that NHEJ favored few junction variants disrupted our initial idea to estimate the total number of transgene molecules, which were independently joined by NHEJ and served as template for HR. Nevertheless, information about sequence of the junctions made it possible to check our prediction that the result of the joining of molecules by HR mechanism would be an exact copying of the template junction. For example, different copies with the same barcodes in dHJ patterns (Fig.5) should also have the same variant of junction. We checked it and this is true for most of the cases. Only in 8% of cases, the barcode had not one, but two different variants of the junction (data not shown).

We can roughly estimate efficiency of NHEJ-mediated ligation: the number of independent NHEJ-ligated molecules should be no less than the number of unique junction variants from the embryos. According to our data, this corresponds to 1 event per 10 injected copies. Obviously, this is the lower estimate and the real value is several times higher. We also did not analyze other copy orientation variants (head-to-head or tail-to-tail), because their sequencing was impossible due to technical reasons (see Discussion section and Sup. Fig. 6). In general, we can conclude that NHEJ plays a prominent role during initial ligation of exogenous DNA in zygote.

## Discussion

Concatemers are a prominent feature of pronuclear microinjection method (Sup. Table 1). To our knowledge, the only paper that addressed the mechanism of concatemer formation was published by Mario Capecchi group more than 35 years ago (14). In this seminal experiment, albeit performed on cultured mammalian cells, nuclei were injected with 2-500 copies of DNA molecules (linear or circular). In addition to clarifying various technical aspects, the group also devoted a part of the report to investigating how concatemers are formed. They injected nuclei with a mixture of two similar transgenes, A- and B-molecules, - plasmid backbones with HSV thymidine kinase gene in two orientations. Southern blot analysis with specific restriction enzymes showed that transgene concatemers consisted of interspersed tandemly oriented A/B copies. This elegant effort challenged transgene amplification hypothesis, but authors were cautious about low copy number integrations and random fluctuations due to the presence of only two transgene versions. Our data unequivocally confirm that concatemers are formed by recombination of individual transgenes without *de novo* amplification.

We also managed to obtain decisive evidence that head-to-tail tandems are mostly formed by HR between linear copies. We base this conclusion on the lack of red+blue double connections (self-circularized copies). SDSA and DSBR are two main ways of repairing double-stranded DNA breaks by homologous recombination (5). These pathways have similar initial stages and usually resolve into indiscernible non-crossover products, evident in the form of barcode “switching” in our assay (green connections without blue connection from initial plasmid library). However, DSBR sometimes manifests itself in the formation of crossover products after dHJ resolution. We found several convincing examples of crossovers leaving traces at the concatemer “subway map” (5% copies) (Table 1). As far as we know, this is the first described case of somatic crossing-over in early mammalian embryos. Apparently, the formation of crossovers is quite dangerous for somatic cells as it can lead to the loss of heterozygosity of a large chromosome fragment (32). The fact that crossing-over is not completely suppressed in early embryos is of interest and expands our scarce knowledge of DNA repair at this stage.

In addition to crossovers and barcode “switching”, we also noticed another indication of HR activity. Analysis of junctions and transgene-genome integration sites revealed that sometimes transgene copies contain junction sequences, corresponding to the D-loop disruption intermediates (33). One can imagine that resected transgene’s 3’-end invades homologous template at the transgene-transgene junction, copies a portion of junction, and then, after D-loop disruption, it gets incorporated into concatemer or genome by NHEJ or MMEJ (Sup. Fig. 19). We also noticed similar pattern in some published transgene integration models (34, 35). We have two reasons to suspect SDSA involvement, instead of traditionally accepted random breakage explanation. First of all, junctions which border these fragments lack blue barcode connection and does not originate from a self-ligated copy. Second, these fragments are frequently terminated at the barcode sequence, which hints that synthesized displaced strand probably could not reinvade homologous duplex, because of barcode heterogeneity. These cases demonstrate that HR intermediates could be processed by NHEJ at par with fragmented copies. It is also worth mentioning, that we couldn’t directly detect activity of alternative HR pathways, such as single-strand annealing (SSA) and break-induced replication (BIR). SSA might potentially link randomly broken transgene circles (formed after initial circularization) and therefore result in red + blue connections, which were in fact very rare. BIR would manifest itself as amplification of contiguous regions of concatemers, and, indeed, we found one region encompassing 3 transgene molecules with double read counts (embryo #3) (Fig. 2B), but this was a single event. We deduced that these pathways do not contribute to the concatemer formation, probably because long-range resection is prevented by competition with SDSA (36).

The real proportion of NHEJ-processed copies in concatemers has always remained enigmatic. Southern blot estimations from many transgenic mouse lines (Sup. Table 1) confirm that head-to-tail orientation is dominant (>90% of copies vs 50% in case of random ligation). Likewise, our transgene “subway maps” display contiguous tandem head-to-tail chains (>10 copies) with no gaps (e.g. in embryos #1 (Sup. Fig. 7), #4 (Sup. Fig. 10A) or #9 (Sup. Fig. 15A)). Unfortunately, studying complex rearrangements in concatemers is nearly impossible at present, because PCR is not suitable for detection of palindromic junctions (our experience; (37)), and repetitive nature of concatemers complicates NGS-based methods (38, 39). However, it is well established that NHEJ often contributes to concatemer emergence with fragmented and truncated copies arranged in random orientation (38, 40). We ourselves detected truncated copies in most of transgenic embryos and their abundance correlated with transgene copy number (Sup. Fig. 5A,B).

Why is there no perfect palindromic head-to-head or tail-to-tail junctions then? Palindromic sequences are quite stable in mammalian cells (41), thus palindromes are likely wiped out at the initial step, during extrachromosomal recombination. We documented high activity of HR recombination between ends (>80% of copies) and it made us believe that frequent strand invasion and D-loop formation could provoke secondary structures such as hairpins and cruciforms in template palindromic junctions (Sup. Fig. 20). Another possible HR-related mechanism is folding back of the single-stranded resected end of a linear transgene after copying fragment of the palindromic junction (42). These hairpin structures are next recognized and removed by cell repair systems.

Ultimately, we aimed to utilize barcode connections to figure out unequivocal transgene order in concatemers. The closest we achieved this goal was in embryo #1 (only 1 gap unresolved) (Sup. Fig. 7), embryo #9 (Sup. Fig. 15A) and embryo #10 (Sup. Fig. 16). In embryo #9, tandemly oriented chains of variable lengths are presumably separated by inverted copies (full-sized or truncated). We sequenced some of them (Sup. Fig. 15B). Lastly, embryo #10 puzzled us with complex barcode distribution pattern. As seen in Sup. Fig. 16, transgene “subway map” in this embryo has quite peculiar plan with four transgene copies but only 2 unique barcode pairs (8 barcodes total). We validated duplication of each barcode with ddPCR (Sup. Table 2). We employed long-distance PCR with barcode-specific primers to position all copies and their respective barcodes in the four-copy concatemer (Sup. Fig. 16). It appears that two initially ligated copies with unique barcode pairs underwent recombination and assimilated two other copies. It definitely was not a simple duplication event, because barcodes were shuffled between copies.

### Concluding remarks

Using DNA barcodes helped us to explain some of the long-standing questions in the field of transgenesis. First of all, we showed that hundreds of copies are joined together independently without contribution of long range de novo synthesis. Terminal barcodes were also useful to track self-ligated copies and we found no concatemer contribution from such rings, although it had been frequently proposed that concatemers are formed by recombination of overlapping fragments of broken circular copies. In theory, injection of circular copies that go through random breakage and recombine with backbone regions must lead to complete disappearance of barcode “switching” in our assay. Clearly, understanding HR regulation in zygote will be of great importance for implementation of revolutionary transgenesis methods such as newly designed gene drivers (43) or contiguous DNA assembly by overlaps (44), and might help finetune HR to prevent unwanted recombination that complicates CRISPR/Cas9 knock-in experiments (45).

## Authors Contribution

AS and NB initiated the study. AY constructed barcoded plasmid library. AK and IS carried out the mouse experiments. AS performed the experiments. AS, VF and NB analyzed data. AS and NB wrote the manuscript. All authors read and approved the final version of the manuscript.

---

**Supplementary Figure 1.**
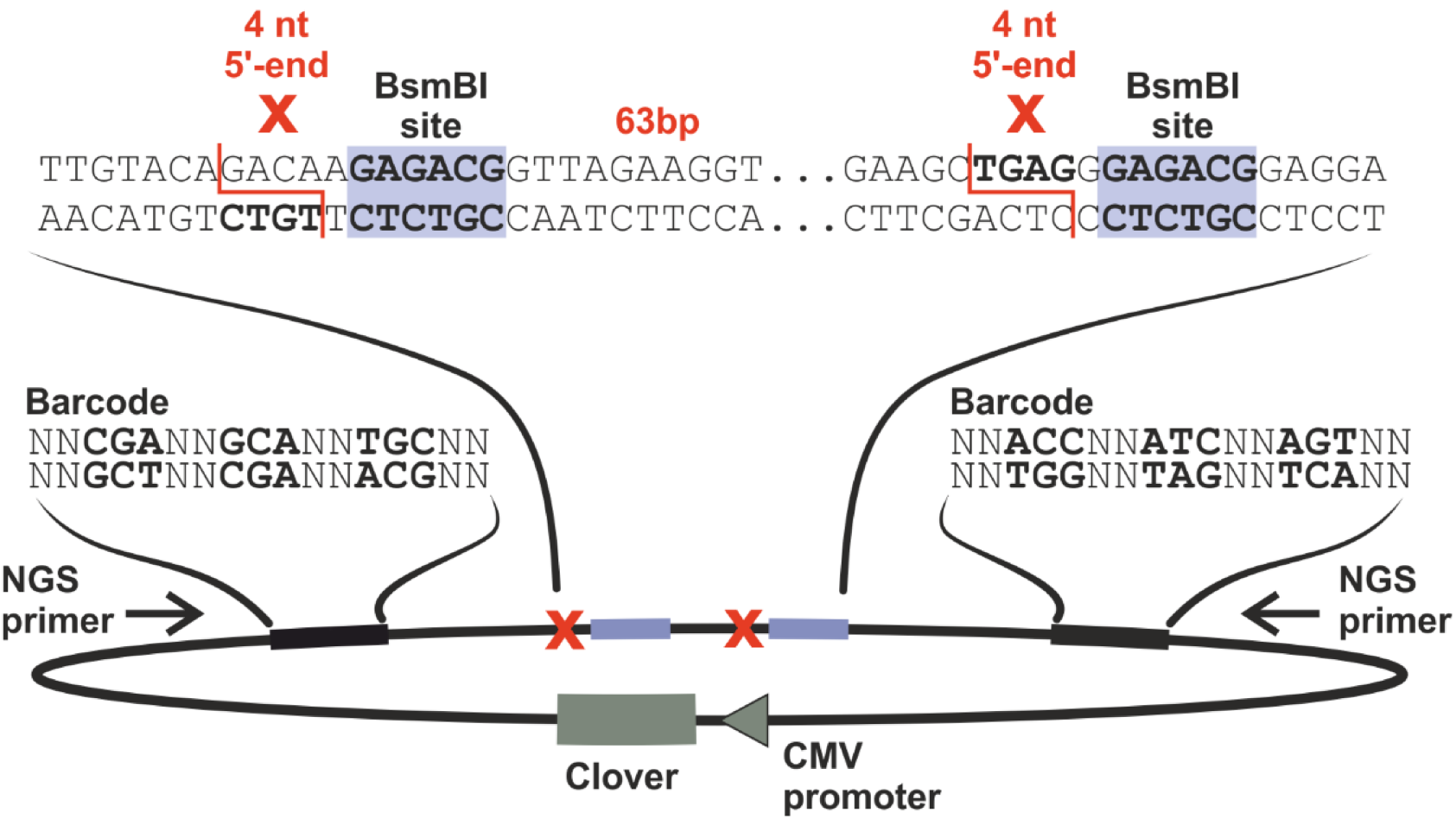
Schematic representation of the barcoded plasmid. Plasmid contains two barcodes separated by a pair of BsmBI restriction sites (blue) producing incompatible ends. NGS primers were used to amplify transgene-transgene junctions and self-ligated copies (inverse PCR). Not to scale.

**Supplementary Figure 2.**
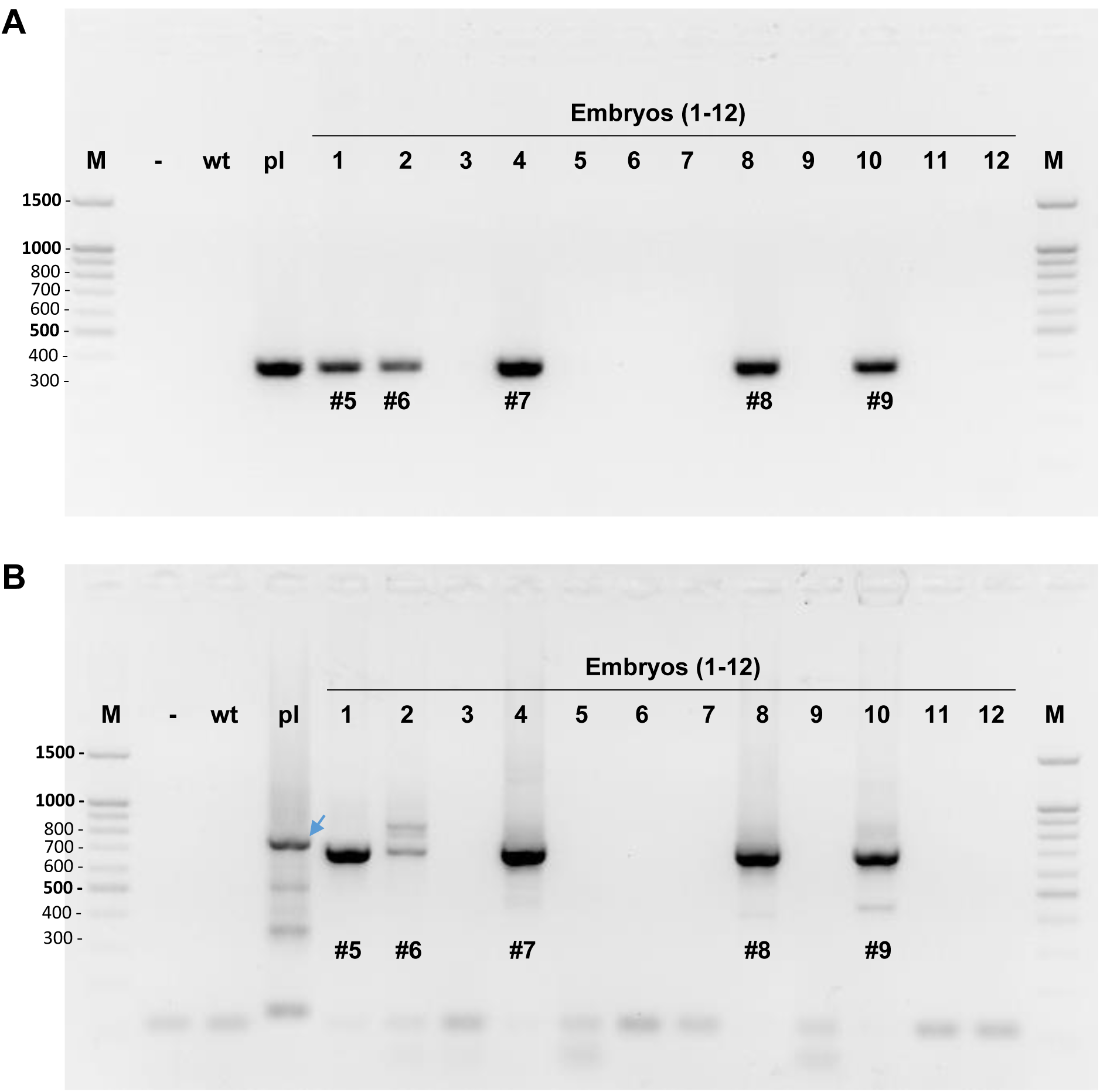
PCR genotyping of the 12 embryos from one of the foster mothers. **(A)** PCR with primers for the Clover gene (product size - 350bp). **(B)** PCR with NGS primers (Sup. Fig. 1) to detect head-to-tail transgene-transgene junctions (product size - ∼700bp). Legend: M – DNA ladder 100bp; wt – wild-type DNA; pl – barcoded plasmid library (arrow indicates corresponding PCR product with BsmBI linker still present); #5-9 – transgenic embryos selected for concatemer sequencing.

**Supplementary Figure 3.**
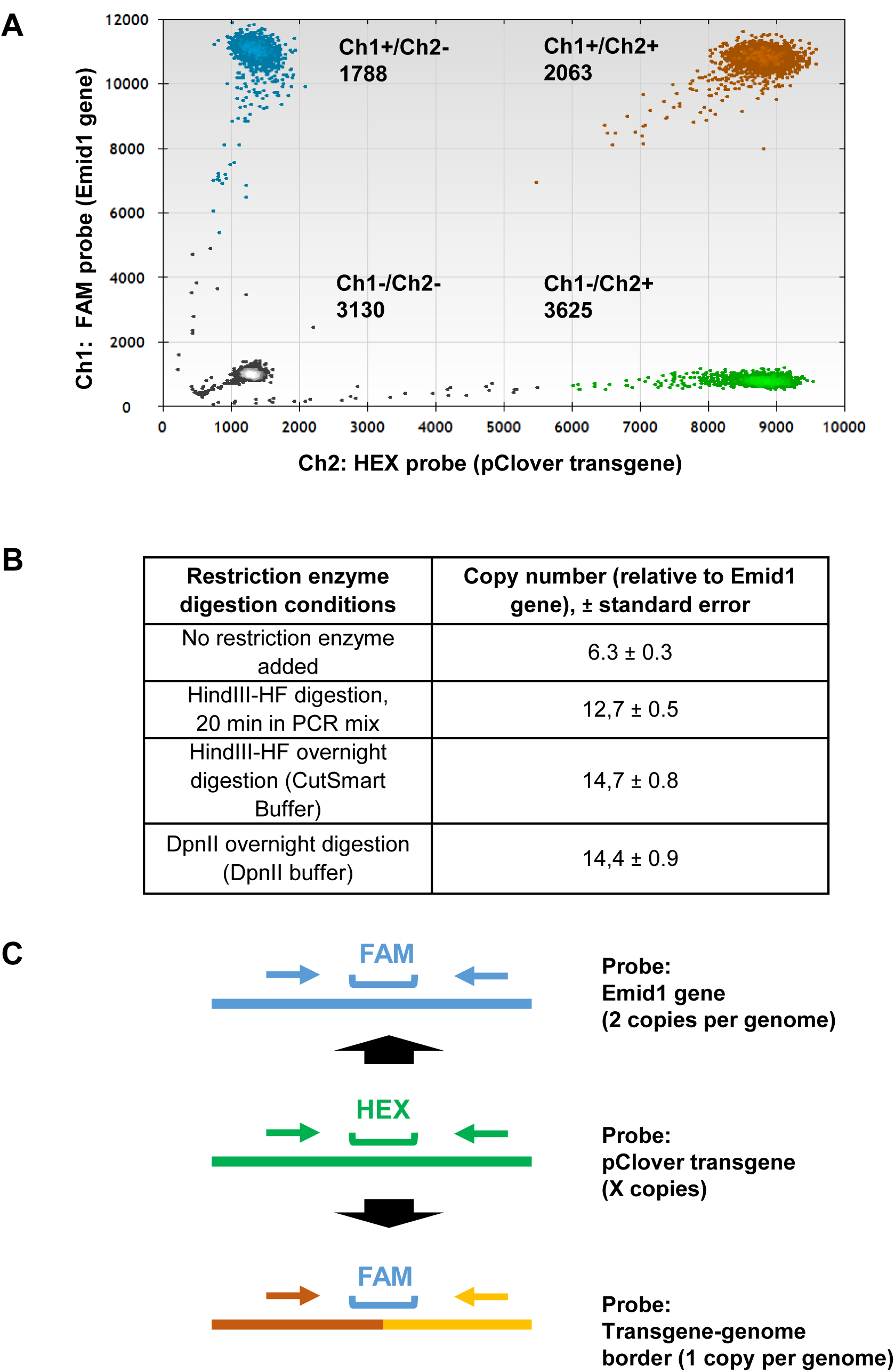
Droplet digital PCR (ddPCR) analysis of transgene copy number. **(A)** Two-dimensional plot of the transgene (pClover) vs control (Emid1 gene) droplet measurement. **(B)** Evalution of the restriction enzyme digestion conditions for optimal separation of the copies in concatemers (DNA from embryo #1). **(C)** Scheme of the ddPCR analysis with probes against transgene-genome border (to exclude mosaicism).

**Supplementary Figure 4.**
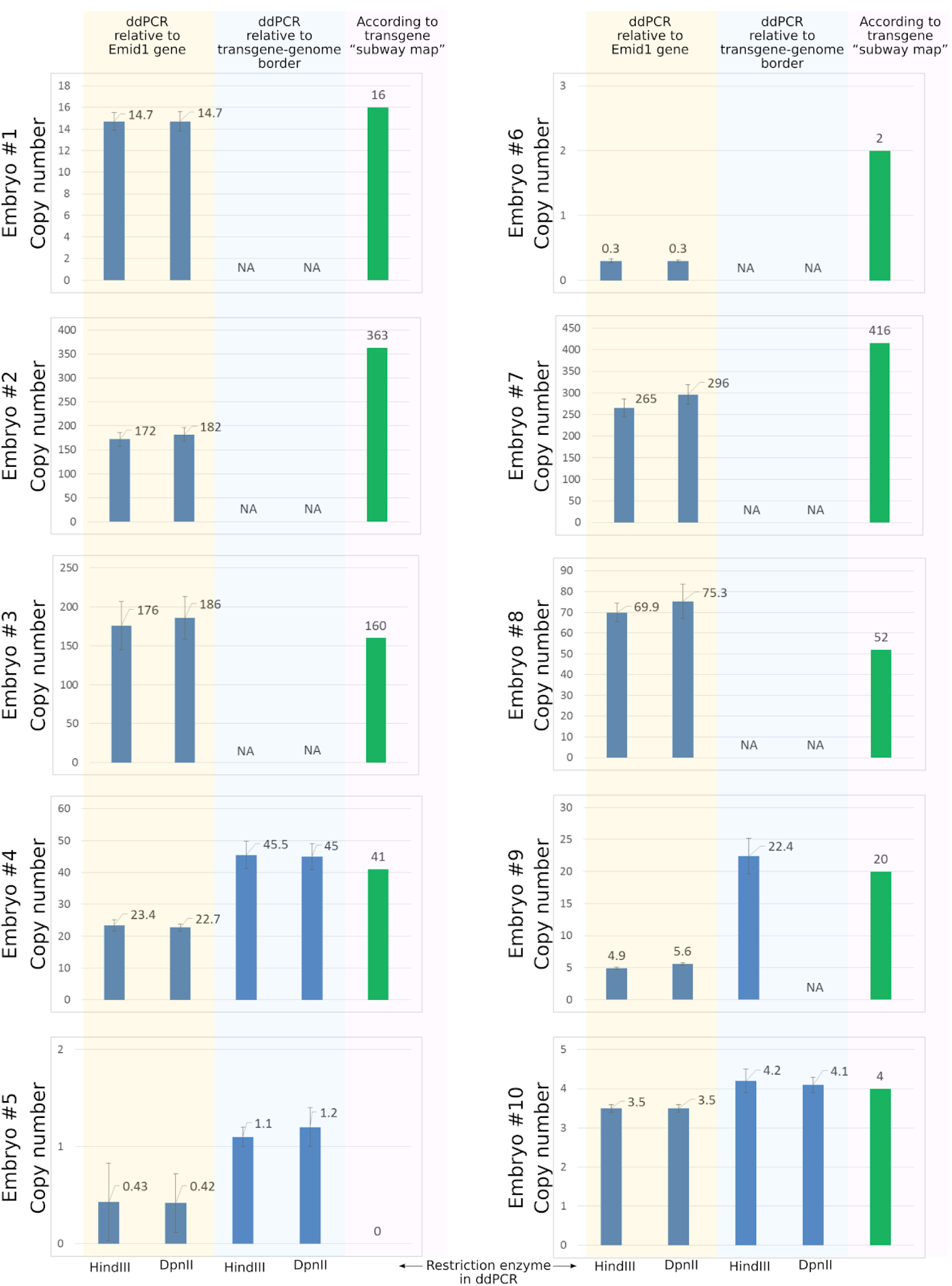
Comparison of transgene copy numbers derived from ddPCR (ivory area), mosaicism-corrected ddPCR (light blue area) and NGS data (green column).

**Supplementary Figure 5A.**
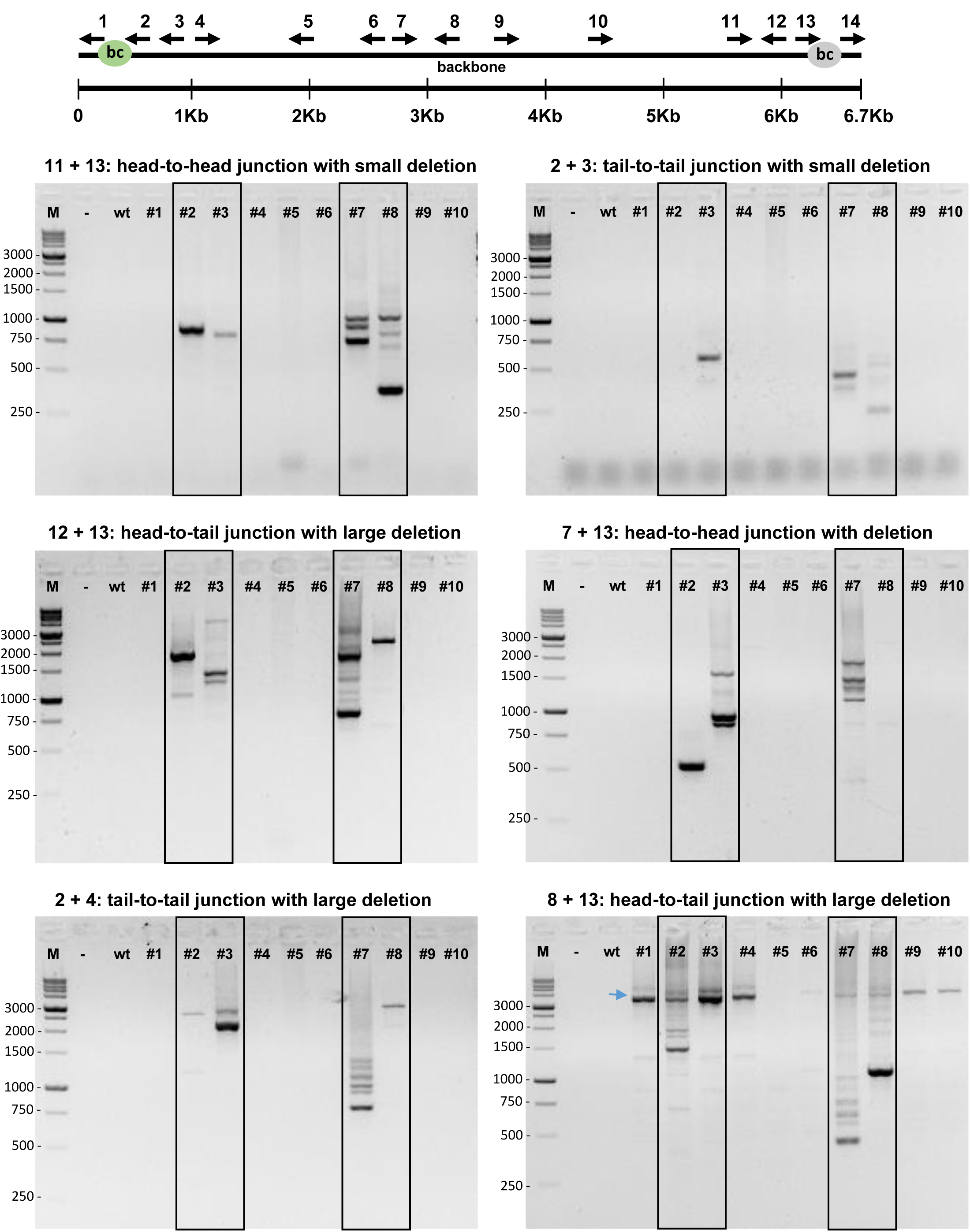
PCR detection of various rearrangements at transgene-transgene junctions. Different primer pairs (positions shown on the transgene map at the top) were used to amplify transgene-transgene junctions. Arrows indicate expected PCR product lengths for intact junctions. Legend: M – Marker (1Kb); - – H2O; wt – wild-type DNA; #1-10 – transgenic DNA. Black frames highlight multicopy embryos #2, 3, 7, 8.

**Supplementary Figure 5B.**
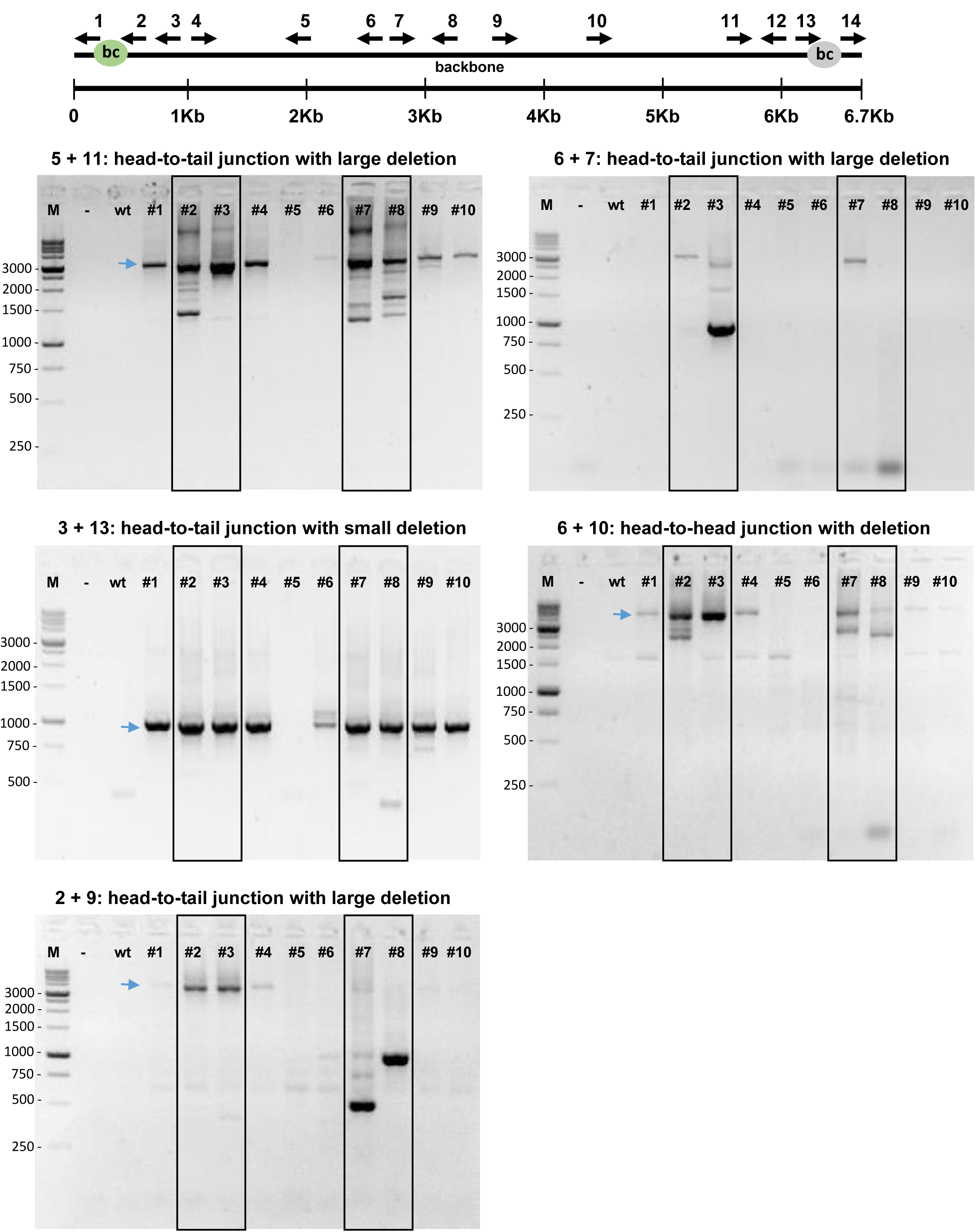
PCR detection of various rearrangements at transgene-transgene junctions. Different primer pairs (positions shown on the transgene map at the top) were used to amplify transgene-transgene junctions. Arrows indicate expected PCR product lengths for intact junctions. Legend: M – Marker (1Kb); - – H2O; wt – wild-type DNA; #1-10 – transgenic DNA. Black frames highlight multicopy embryos #2, 3, 7, 8.

**Supplementary Figure 6A.**
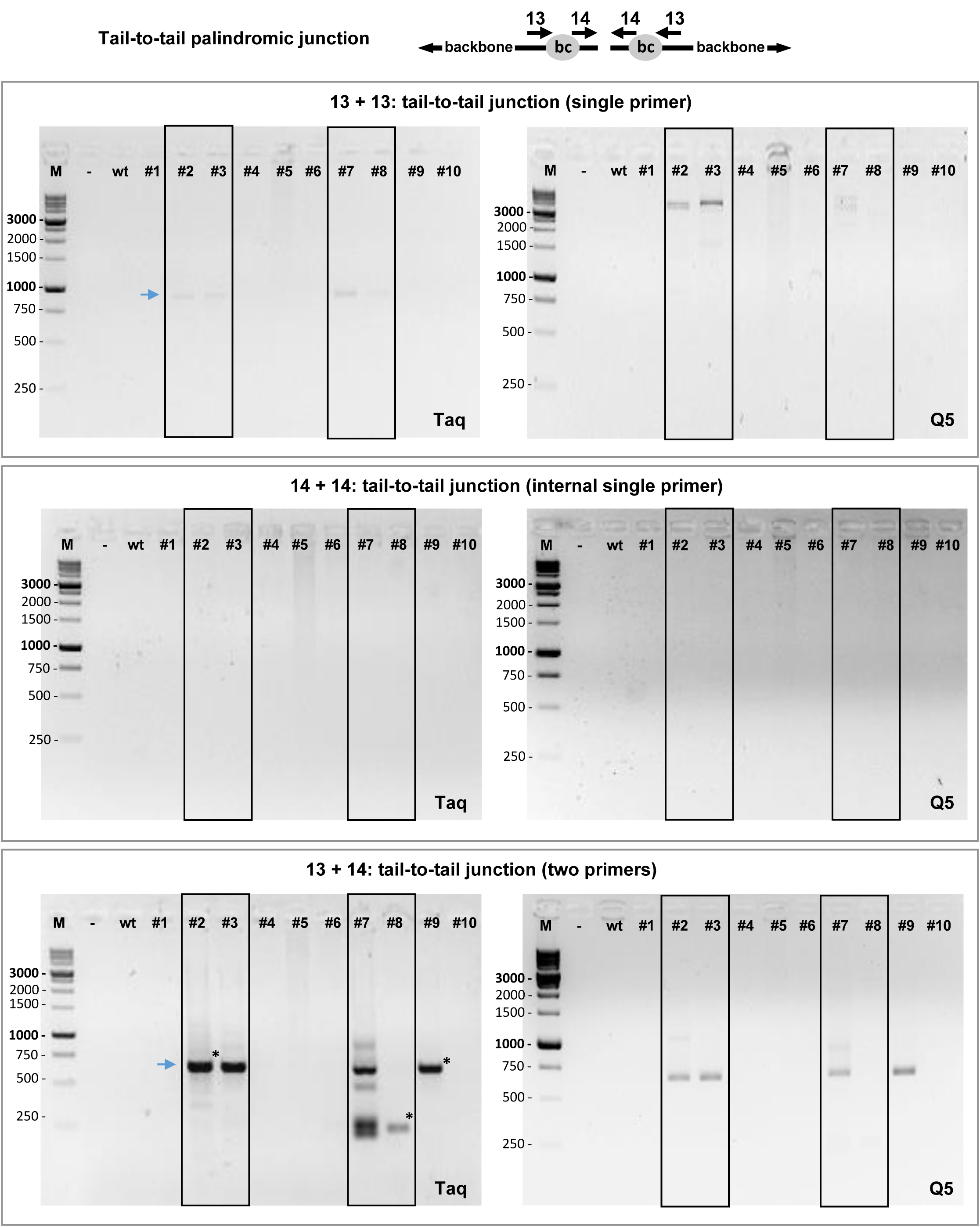
PCR detection of the *tail-to-tail* palindromic junctions with Taq and Q5 polymerases. Either single primers or primer pairs were used to amplify transgene-transgene tail-to-tail junctions. Expected PCR product size for the perfect palindromic junctions – 734 bp (13+13), 282 bp (14+14). Legend: M –Marker (1Kb); - – H2O; wt – wild-type DNA; #1-10 – transgenic DNA. Black frames highlight multicopy embryos #2, 3, 7, 8.

**Supplementary Figure 6B.**
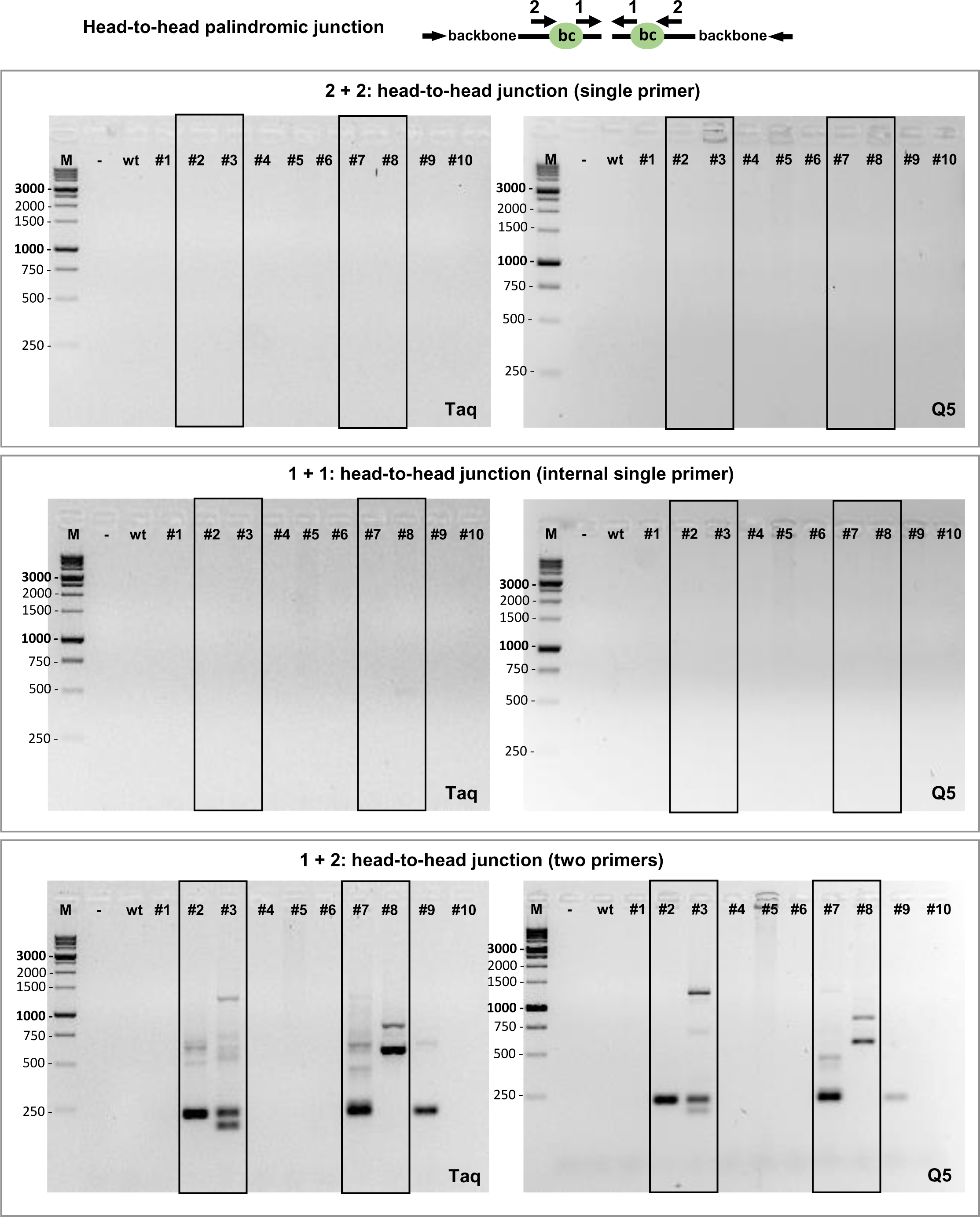
PCR detection of the *head-to-head* palindromic junctions with Taq and Q5 polymerases. Either single primers or primer pairs were used to amplify transgene-transgene head-to-head junctions. Expected PCR product size for the perfect palindromic junctions – 708 bp (2+2), 214 bp (1+1). Legend: M –Marker (1Kb); - – H2O; wt – wild-type DNA; #1-10 – transgenic DNA. Black frames highlight multicopy embryos #2, 3, 7, 8.

**Supplementary Figure 7.**
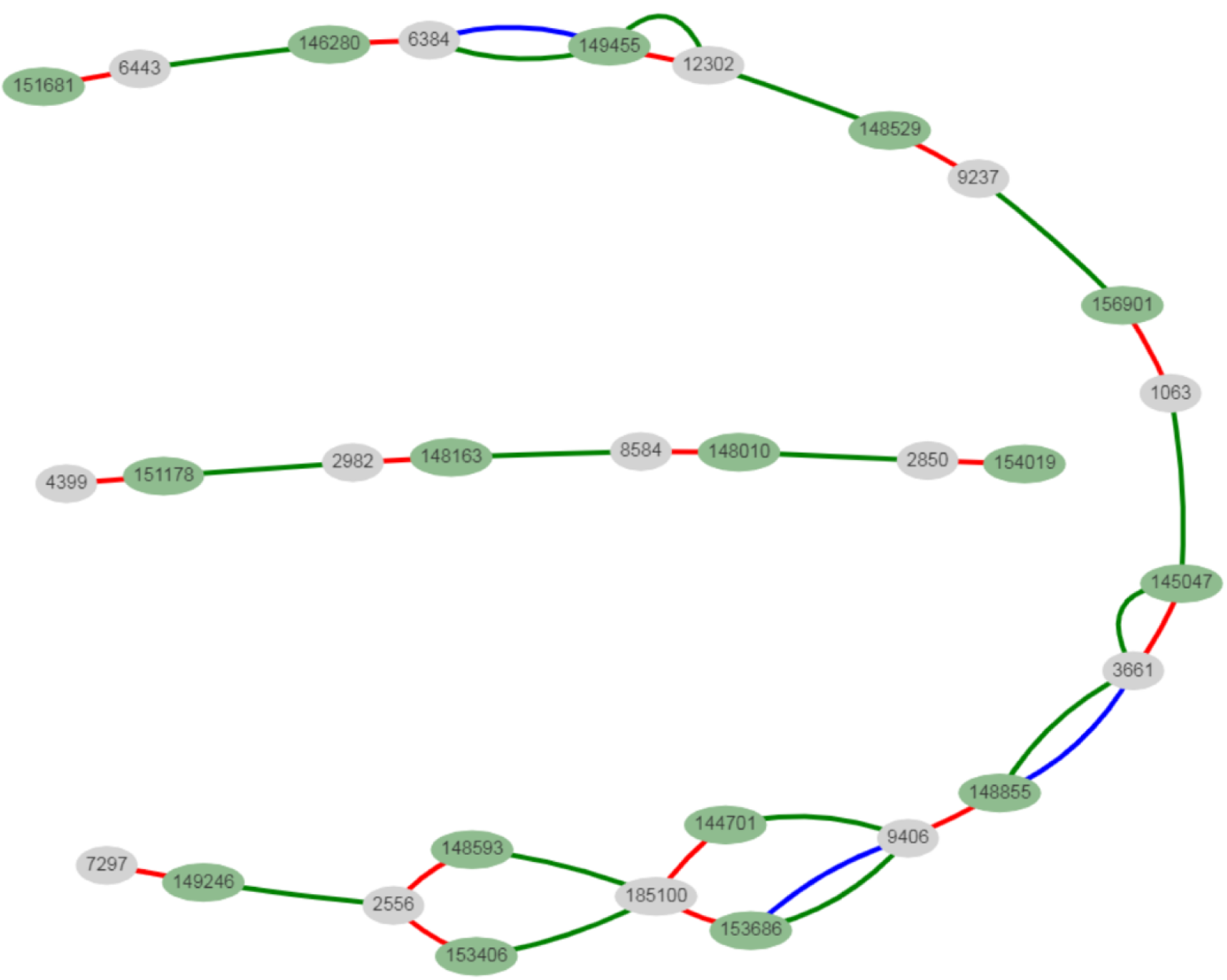
Transgene “subway map” for embryo #1 (File 1). This embryo has 15 transgene copies according to ddPCR. Types of connections: red (head-to-tail junction); blue + green (transgene from original library); only green (recombined transgene with “switched” barcodes); red + green (dHJ pattern 1 – incorporation of the transgene copy). Note branched structure at the bottom region – product of unsynced mismatch repair (see Fig. 3 in main text for explanation).

**Supplementary Figure 8.**
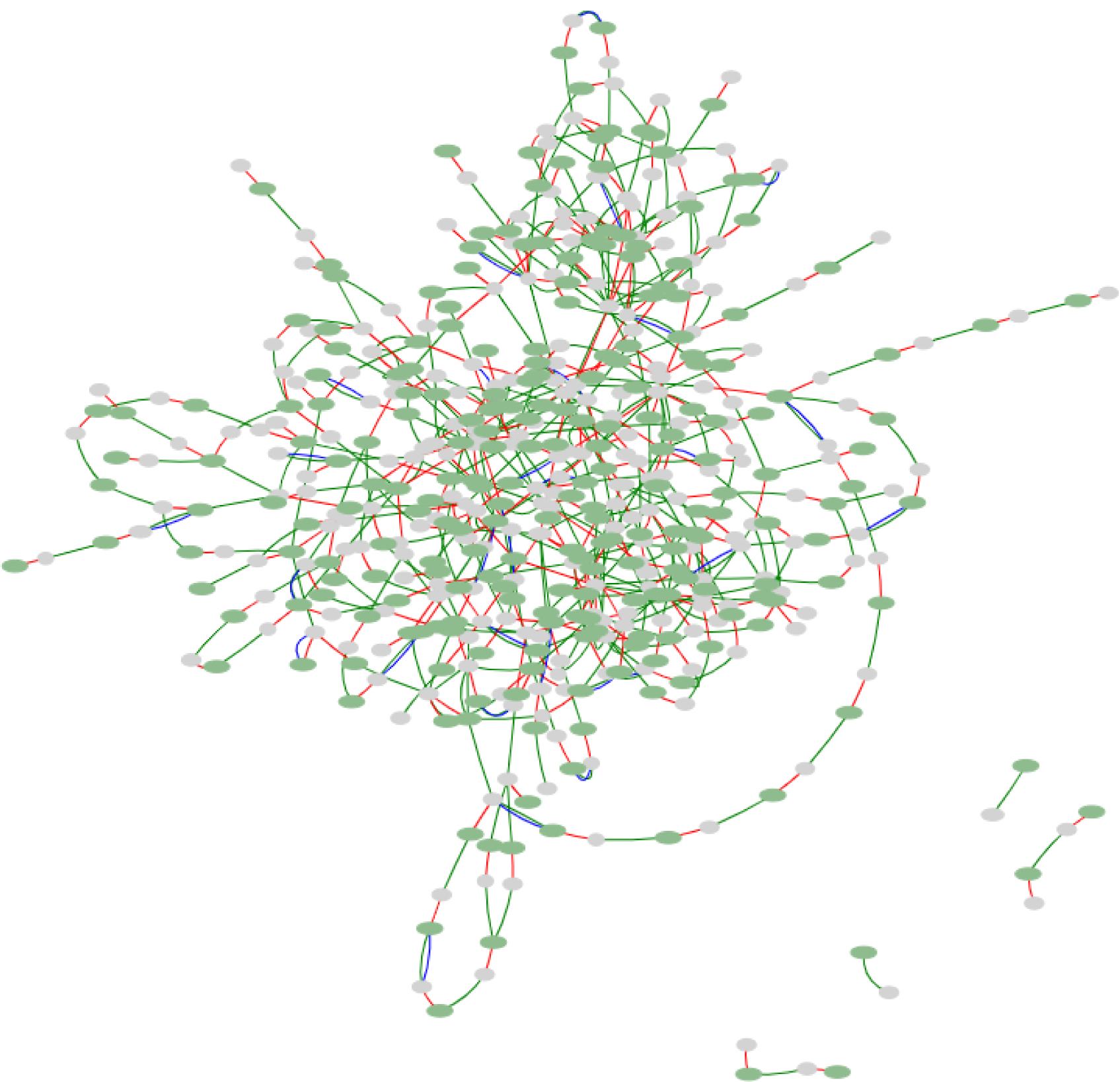
Transgene “subway map” for embryo #2 (File 2). This embryo has around 175 transgene copies according to ddPCR. Types of connections: red (head-to-tail junction); blue + green (transgene from original library); only green (recombined transgene with “switched” barcodes); red + green (dHJ pattern 1 – incorporation of the transgene copy); red + blue (dHJ pattern 3 – transfer of junction barcodes into another junction, but could indicated self-ligated junction as well).

**Supplementary Figure 9.**
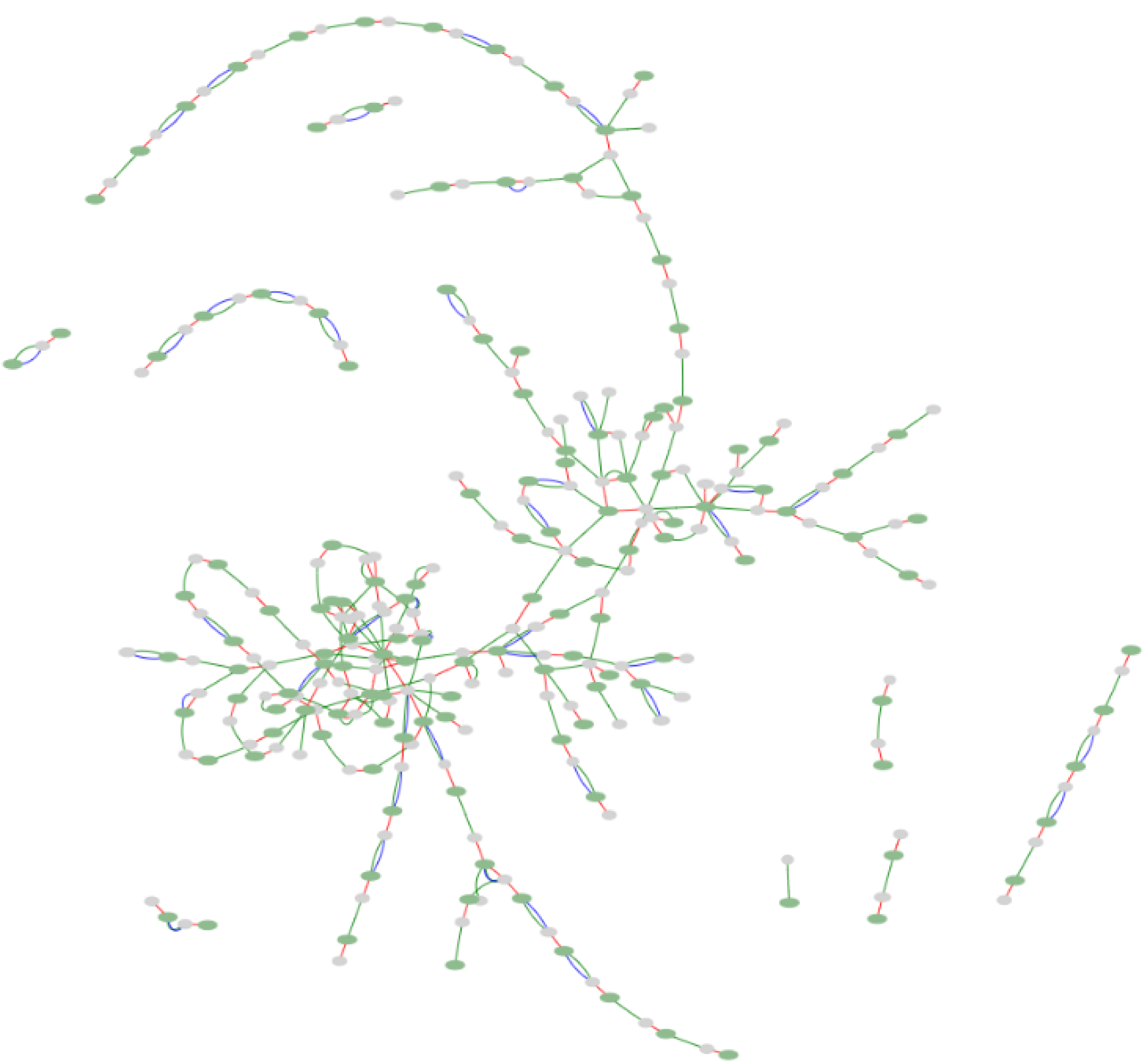
Transgene “subway map” for embryo #3 (File 3). This embryo has around 180 transgene copies according to ddPCR. Types of connections: red (head-to-tail junction); blue + green (transgene from original library); only green (recombined transgene with “switched” barcodes); red + green (dHJ pattern 1 – incorporation of the transgene copy); red + blue (dHJ pattern 3 – transfer of junction barcodes into another junction, but could indicated self-ligated junction as well), red + blue + green (dHJ pattern 2 – incorporation of a circular copy into junction with preservation of invading barcodes).

**Supplementary Figure 10A.**
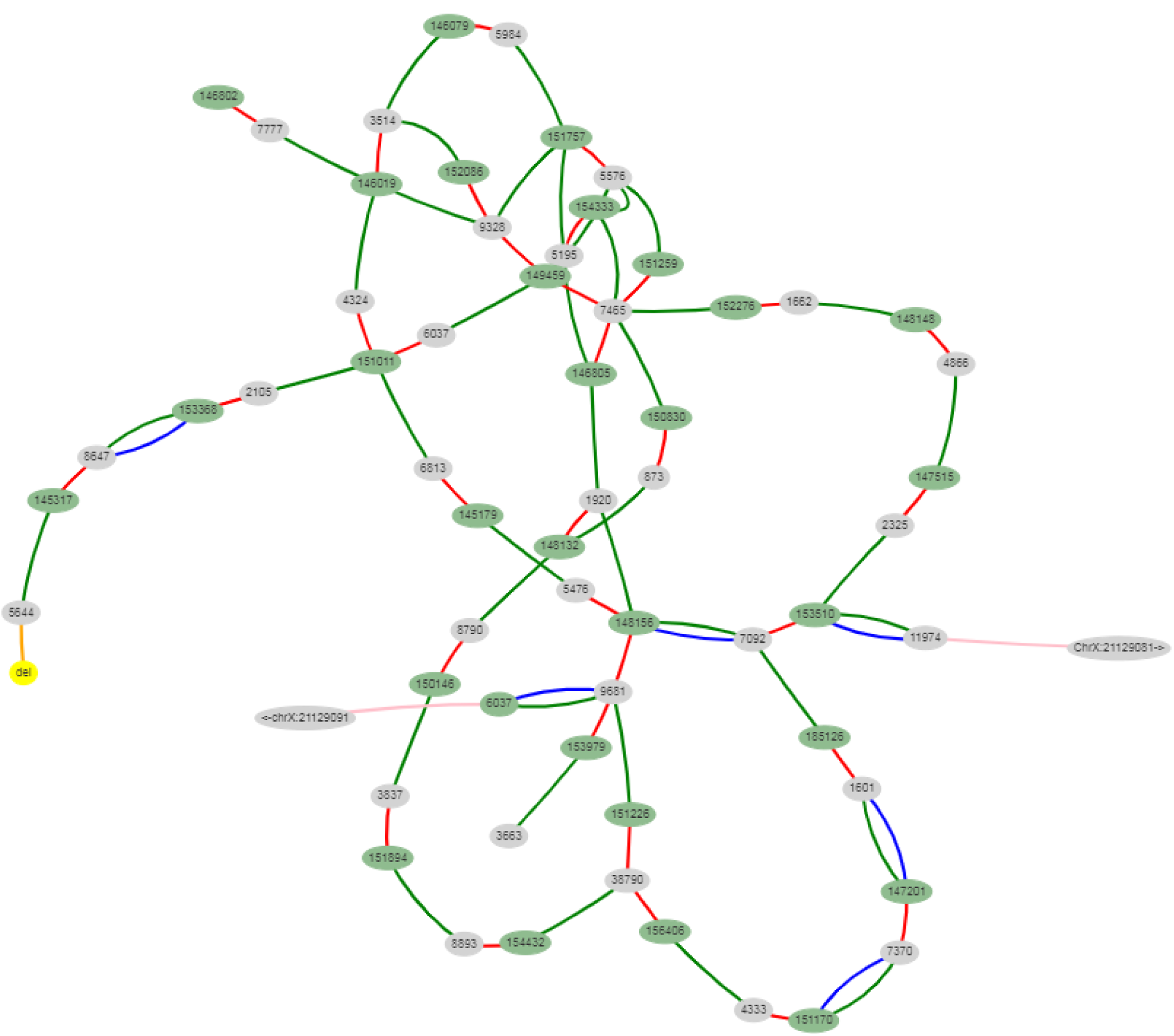
Transgene «subway map» for embryo #4 (File 4). This embryo has around 45 copies and one sequenced rearrangement (deletion + inversion). Types of connections: red (head-to-tail junction); blue + green (transgene from original library); only green (recombined transgene with «switched» barcodes); orange (head-to-head connection with inverted truncated copy) (added manually); pink (transgene-genome borders) (added manually).

**Supplementary Figure 10B.**
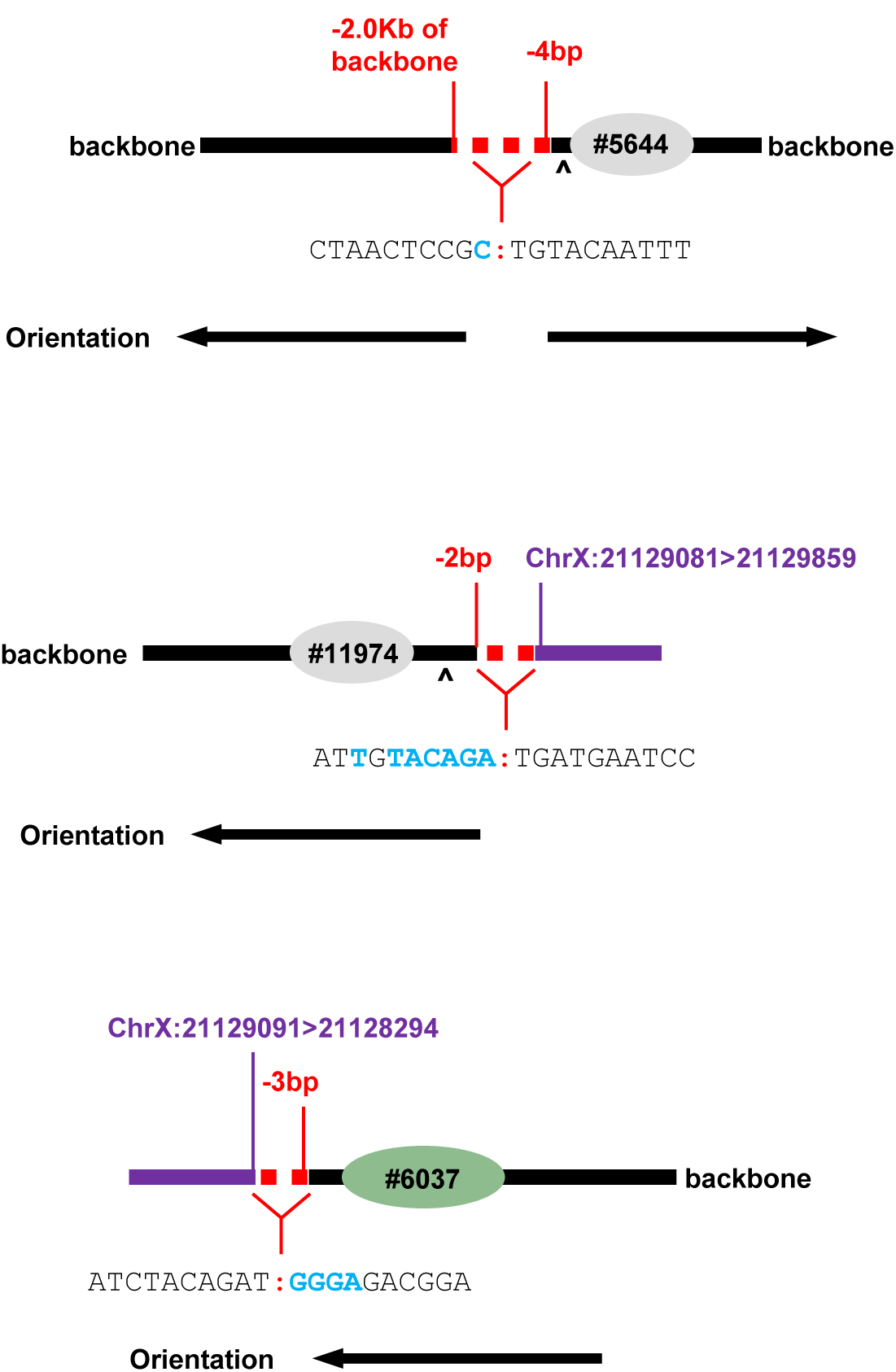
Scheme of the rearranged junction (top) and transgene-genome borders in embryo #4. Triangle indicates PciI site. Blue letters – microhomologies between ends. Chromosome locus did not lose any nucleotides, since broken ends were directly ligated to concatemer by microhomology mediated end-joining.

**Supplementary Figure 11.**
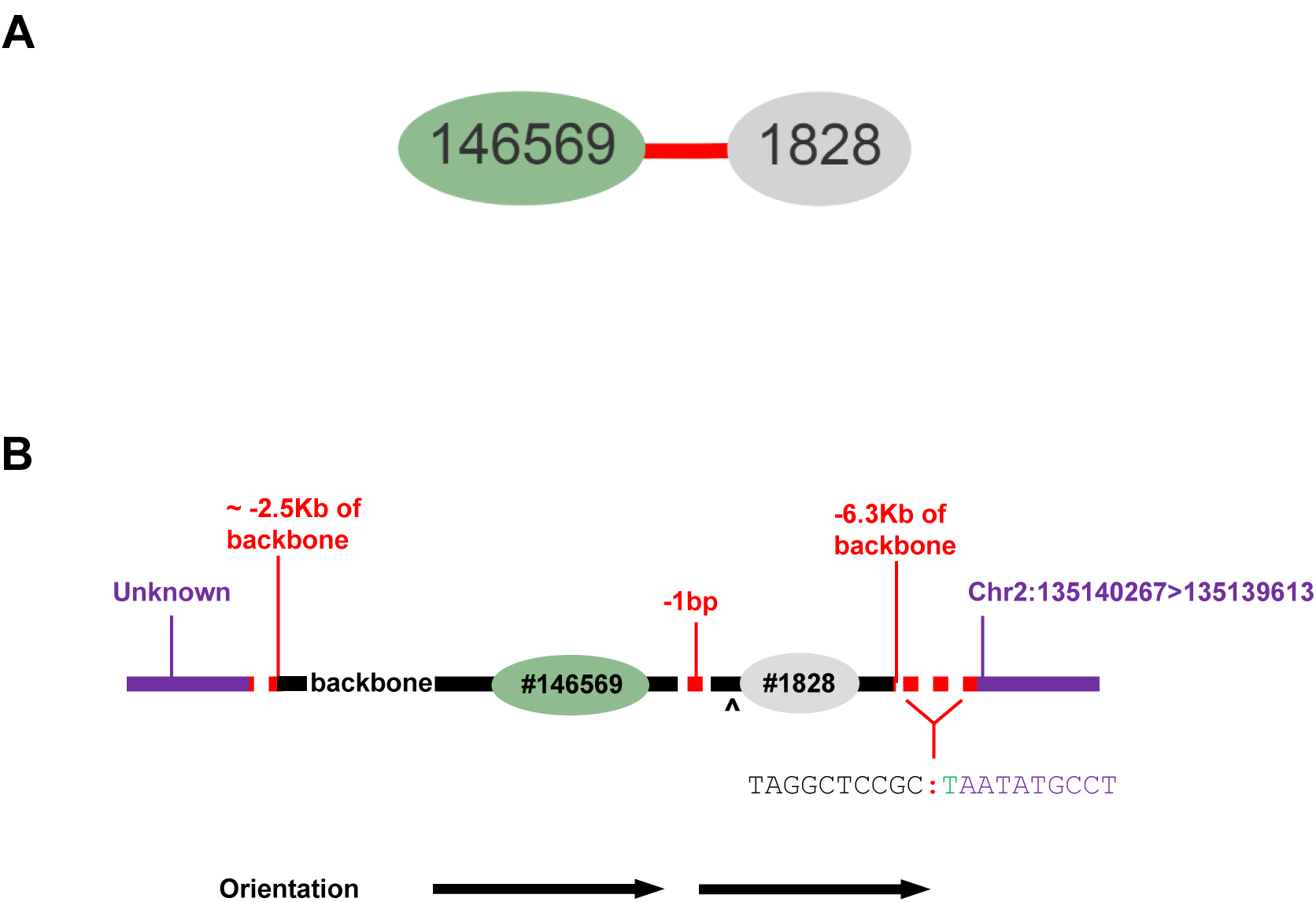
Transgene “subway map” for embryo #5 (File 5). This embryo has 1 truncated copy, which, at first sight, could have probably originated from randomly broken circular copy, but lack of blue connection signifies recombined junction. **(A)** Type of connection: red (head-to-tail junction). **(B)** Scheme of the transgene-transgene junction and transgene-genome border. Triangle indicates PciI site. Green letter – insertion of 1 bp at the transgene-genome junction.

**Supplementary Figure 12.**
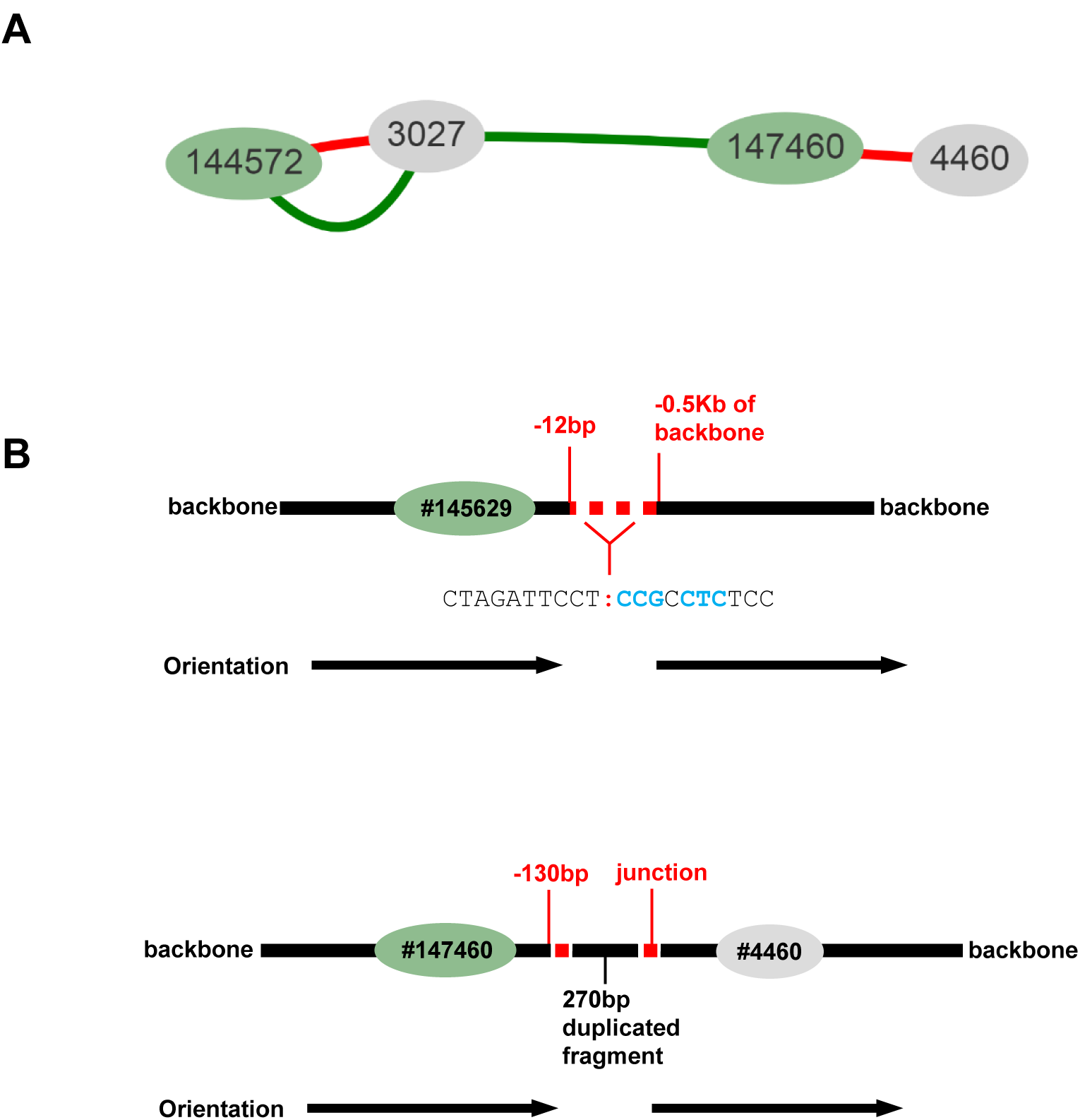
Transgene «subway map» for embryo #6 (File 6). This embryo has 2 copies and two rearrangements. **(A)** Type of connection: red (head-to-tail junction); only green (recombined transgene with “switched” barcodes); red + green (dHJ pattern 1 – incorporation of the transgene copy). **(B)** Two sequenced rearrangements in embryo #6: deletion (top) and duplication of the junction fragment (below). Triangle indicates PciI site. Blue letters – microhomologies between ends.

**Supplementary Figure 13.**
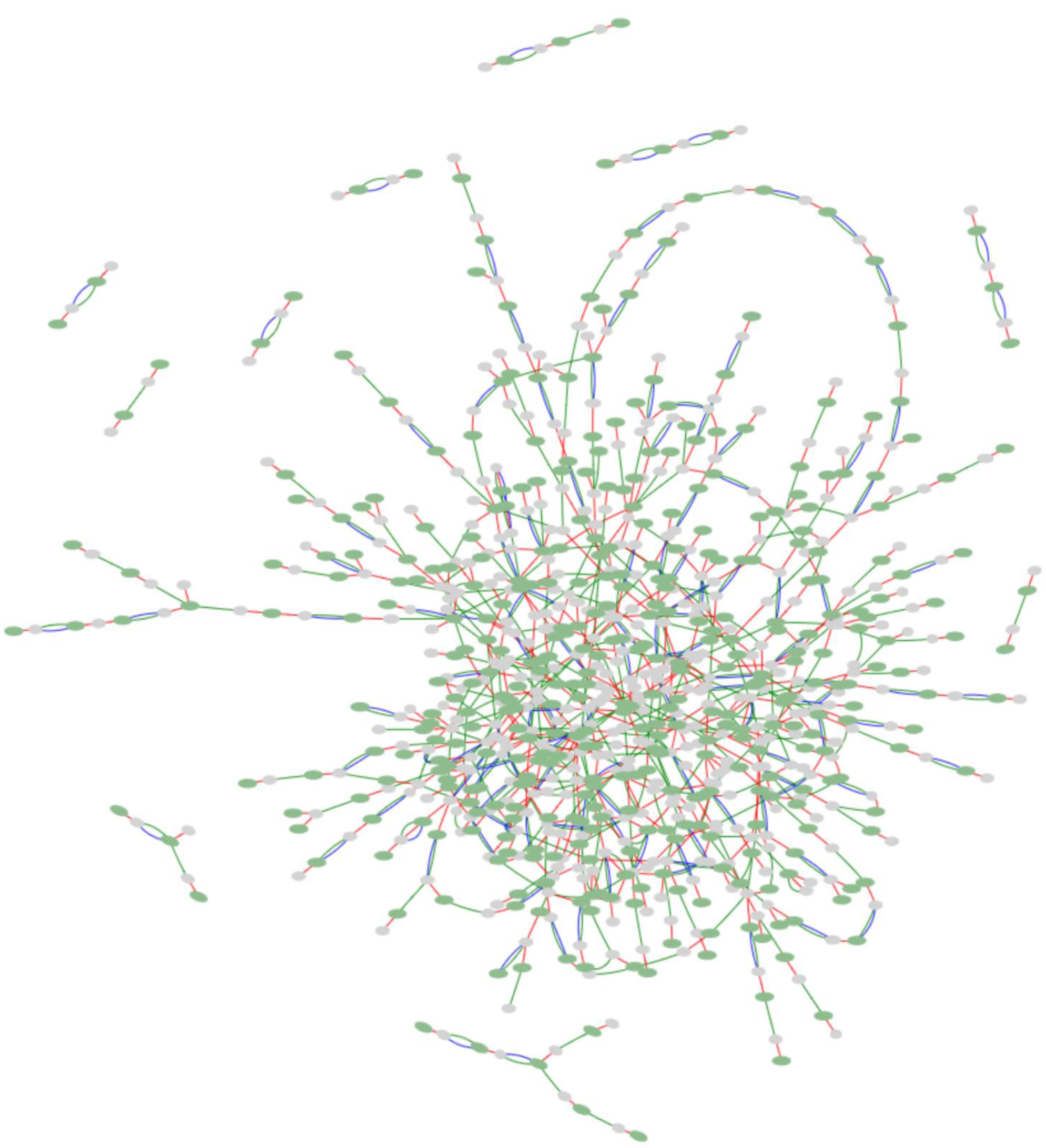
Transgene “subway map” for embryo #7 (File 7). This embryo has around 280 transgene copies according to ddPCR. Types of connections: red (head-to-tail junction); blue + green (transgene from original library); only green (recombined transgene with “switched” barcodes); red + green (dHJ pattern 1 – incorporation of the transgene copy); red + blue (dHJ pattern 3 – transfer of junction barcodes into another junction, but could indicated self-ligated junction as well), red + blue + green (dHJ pattern 2 – incorporation of a circular copy into junction with preservation of invading barcodes).

**Supplementary Figure 14.**
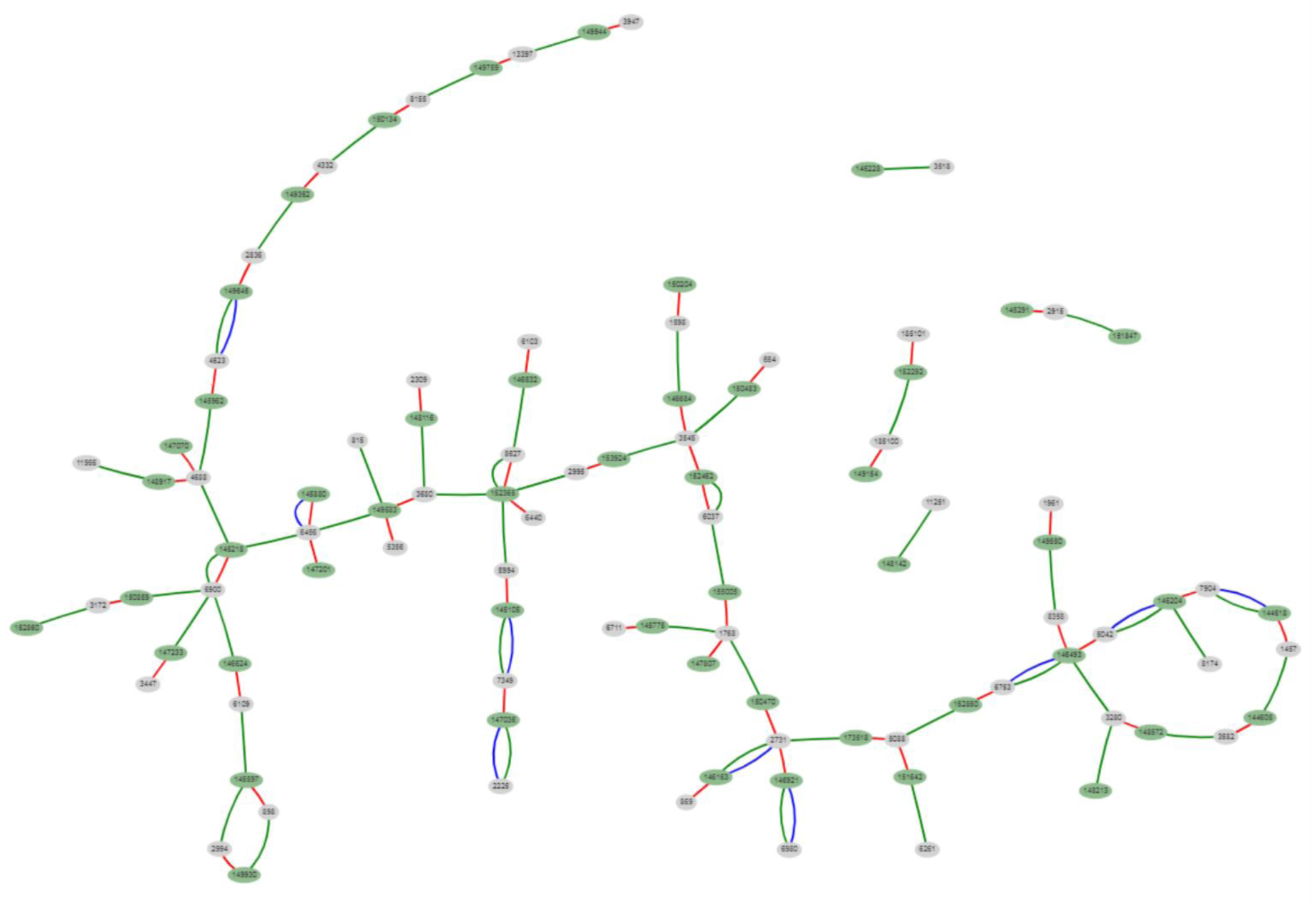
Transgene “subway map” for embryo #8 (File 8). This embryo has around 72 transgene copies according to ddPCR. Types of connections: red (head-to-tail junction); blue + green (transgene from original library); only green (recombined transgene with “switched” barcodes); red + green (dHJ pattern 1 – incorporation of the transgene copy); red + blue (dHJ pattern 3 – transfer of junction barcodes into another junction, but could indicated self-ligated junction as well).

**Supplementary Figure 15A.**
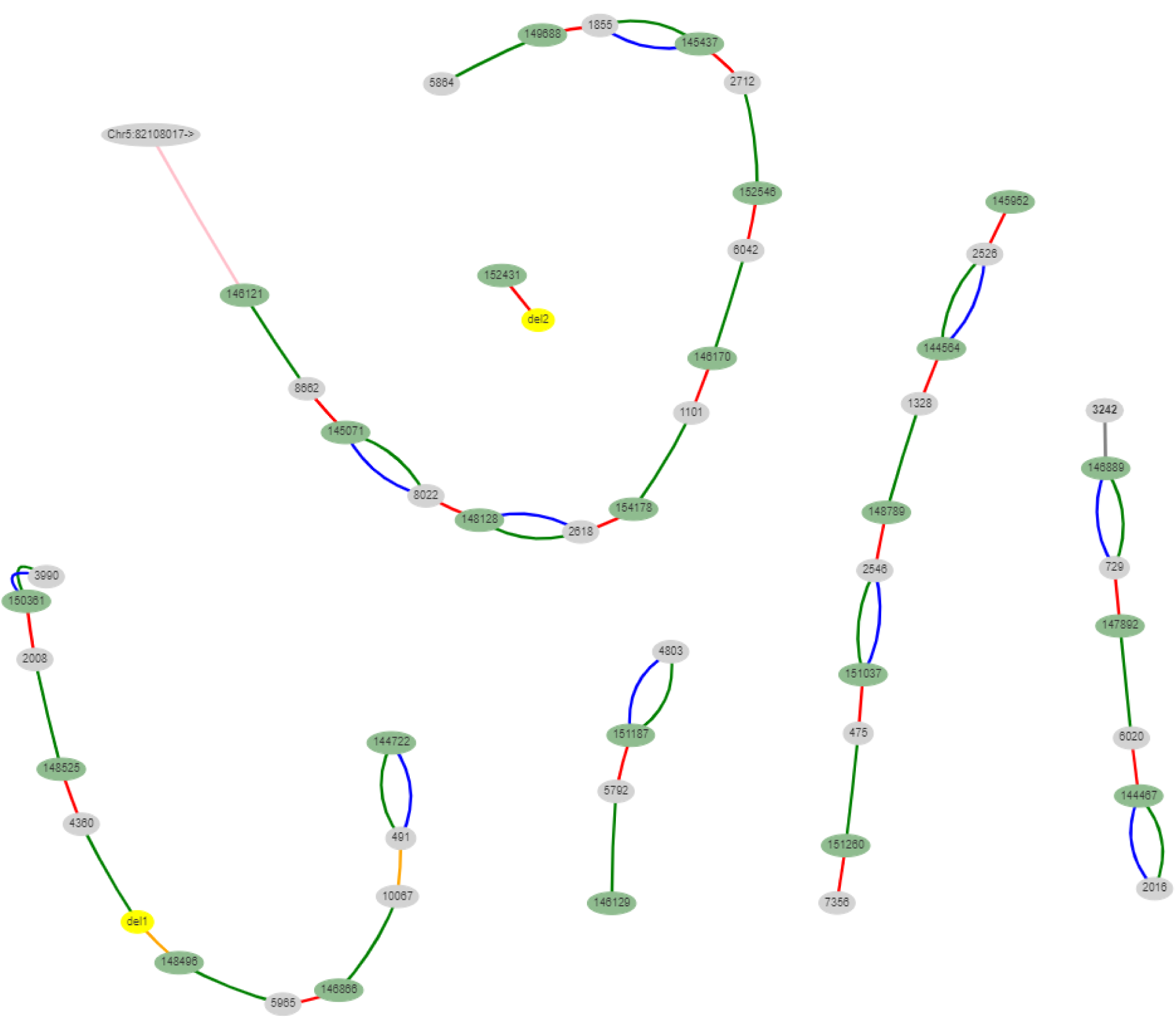
Transgene “subway map” for embryo #9 (File 9). This embryo has around 22 transgene copies according to ddPCR. Types of connections: red (head-to-tail junction); blue + green (transgene from original library); only green (recombined transgene with “switched” barcodes); orange (head-to-head or tail-to-tail connection with inverted copy) (added manually); pink (transgene-genome borders) (added manually).

**Supplementary Figure 15B.**
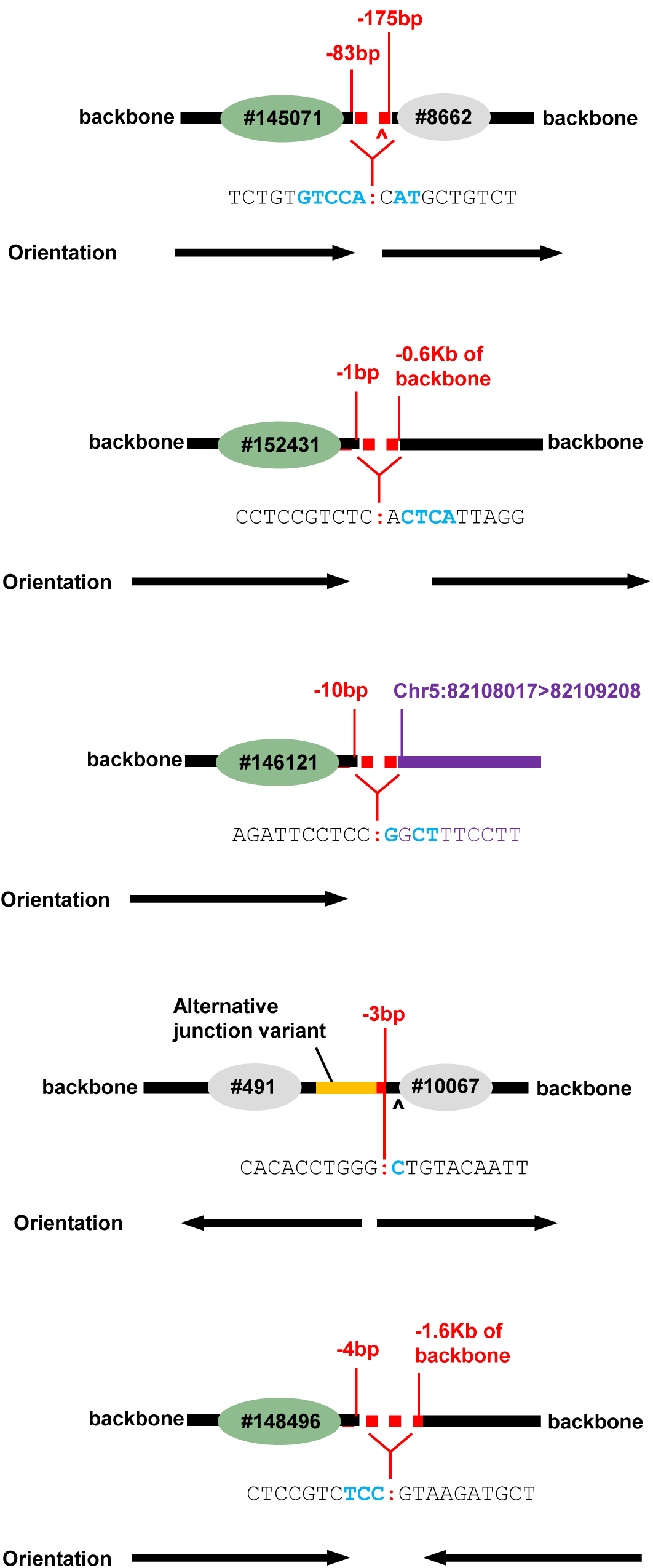
Transgene “subway map” for embryo #9 (File 9). This embryo has several inverted and truncated junctions. Triangle indicates PciI site. Blue letters – microhomologies between ends. Note that in one case, alternative junction variant was introduced in plasmid backbone during cloning (orange fragment). This “palindromic” junction is readily amplified by PCR, because junction fragments are different and introduce asymmetry. Other embryos also have this “quasi-palindromic” junctions (Sup. Fig. 6)

**Supplementary Figure 16.**
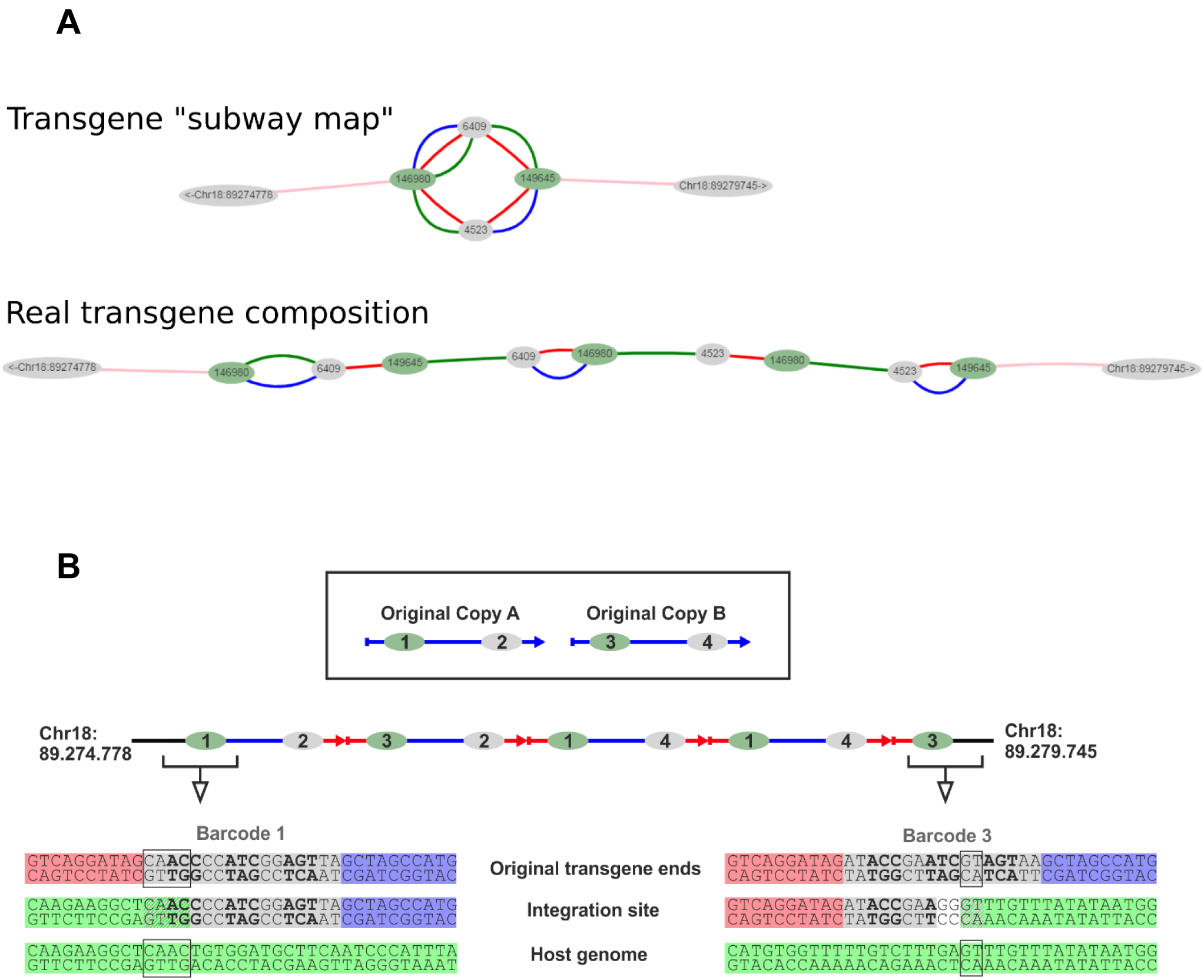
Transgene “subway map” for embryo #10 before and after PCR validation (File 10). This embryo has 4 transgene copies according to ddPCR. **(A)** Types of connections: red (head-to-tail junction); blue + green (transgene from original library); only green (recombined transgene with “switched” barcodes); pink (transgene-genome borders) (added manually). **(B)** Simplified representation of concatemer in embryo #10, showing transgene-genome borders that have microhomologies directly at barcode sequences.

**Supplementary Figure 17.**
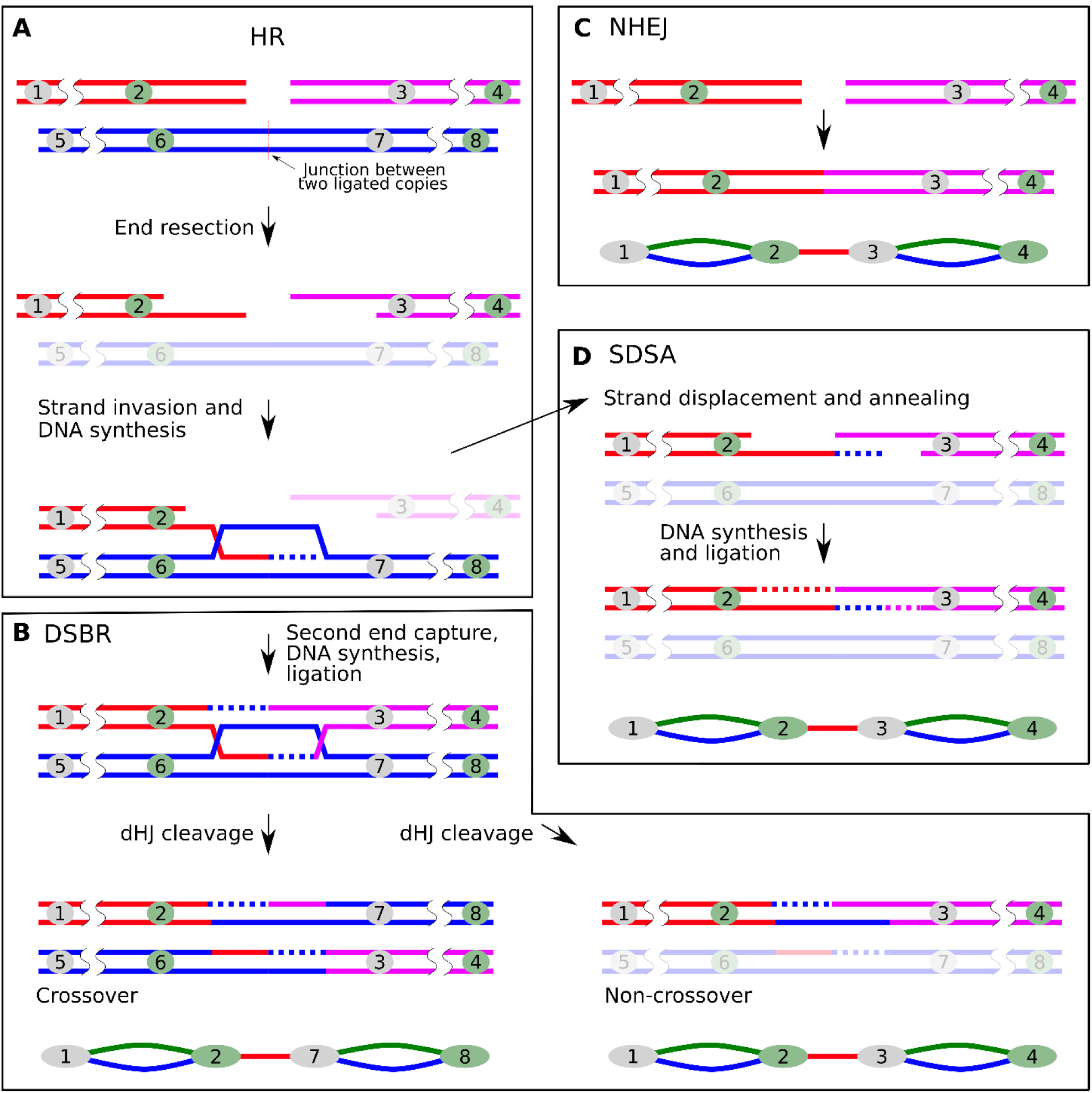
Joining of transgene copies in concatemer by DSB repair pathways. **(A)** Homologous recombination is initiated by end resection of two linear transgenes (1/2 and 3/4 barcodes). At the second step, filaments invade into homologous template region, a head-to-tail junction between transgenes (5/6 and 7/8). **(B, D)** Depending on the pathway choice, both ends could be either captured into dHJ (DSBR) or independently anneal complementary ends after strand displacement (SDSA). **(C)** NHEJ directly ligates two copies after end processing. Note that in the cases shown here, DSB repair does not lead to barcode “switching”, because resection does not reach barcodes (in reality most of barcodes are copied into the invading copy).

**Supplementary Figure 18.**
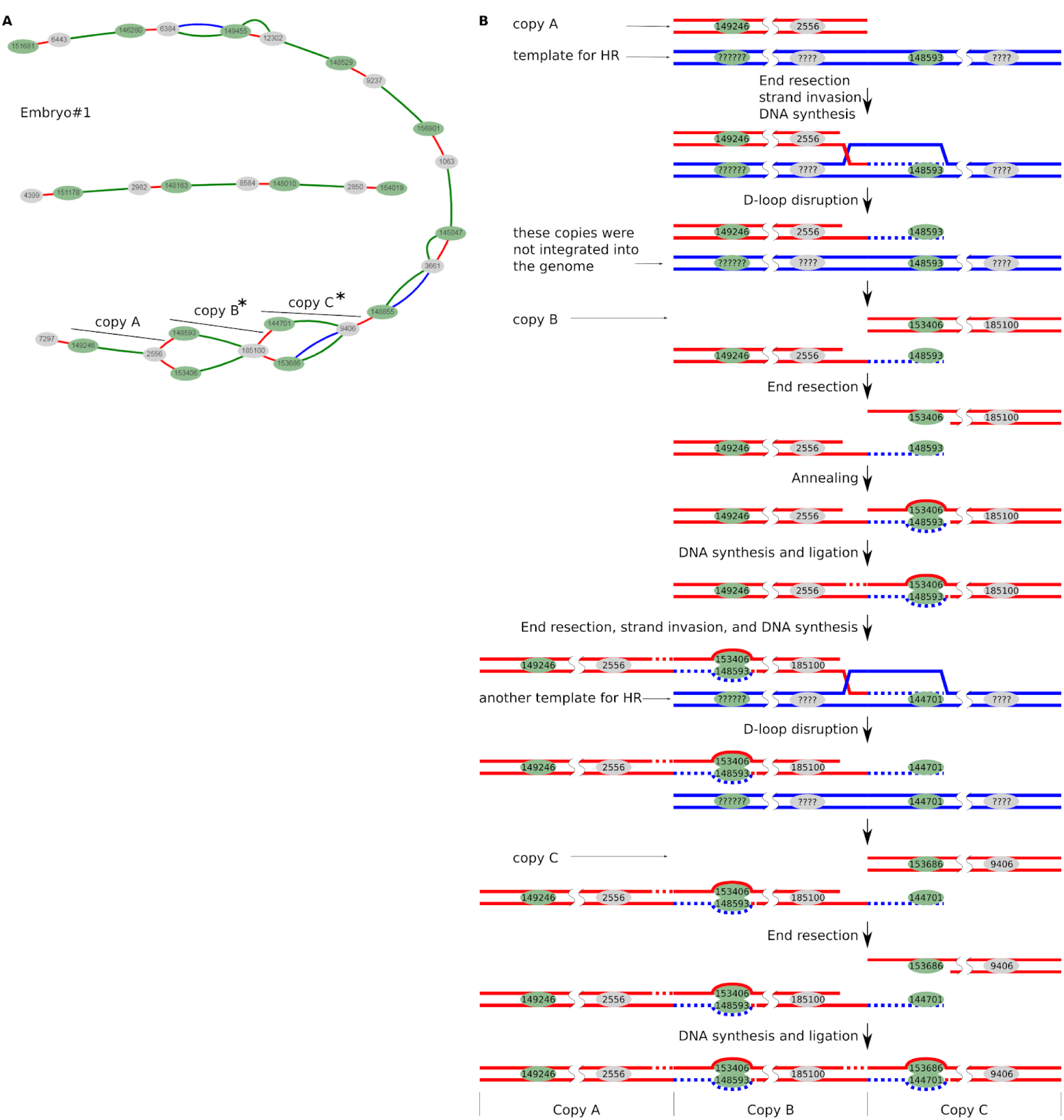
Detailed scheme of the unsynced mismatch repair in embryo #1 (see Fig.3 in main text). **(A)** Transgene “subway map” for embryo #1. Alternative barcodes are present in the copies B and C. **(B)** Several rounds of strand annealing could led to accumulation of unrepaired mismatches at two regions of newly formed concatemer (copies B and C).

**Supplementary Figure 19.**
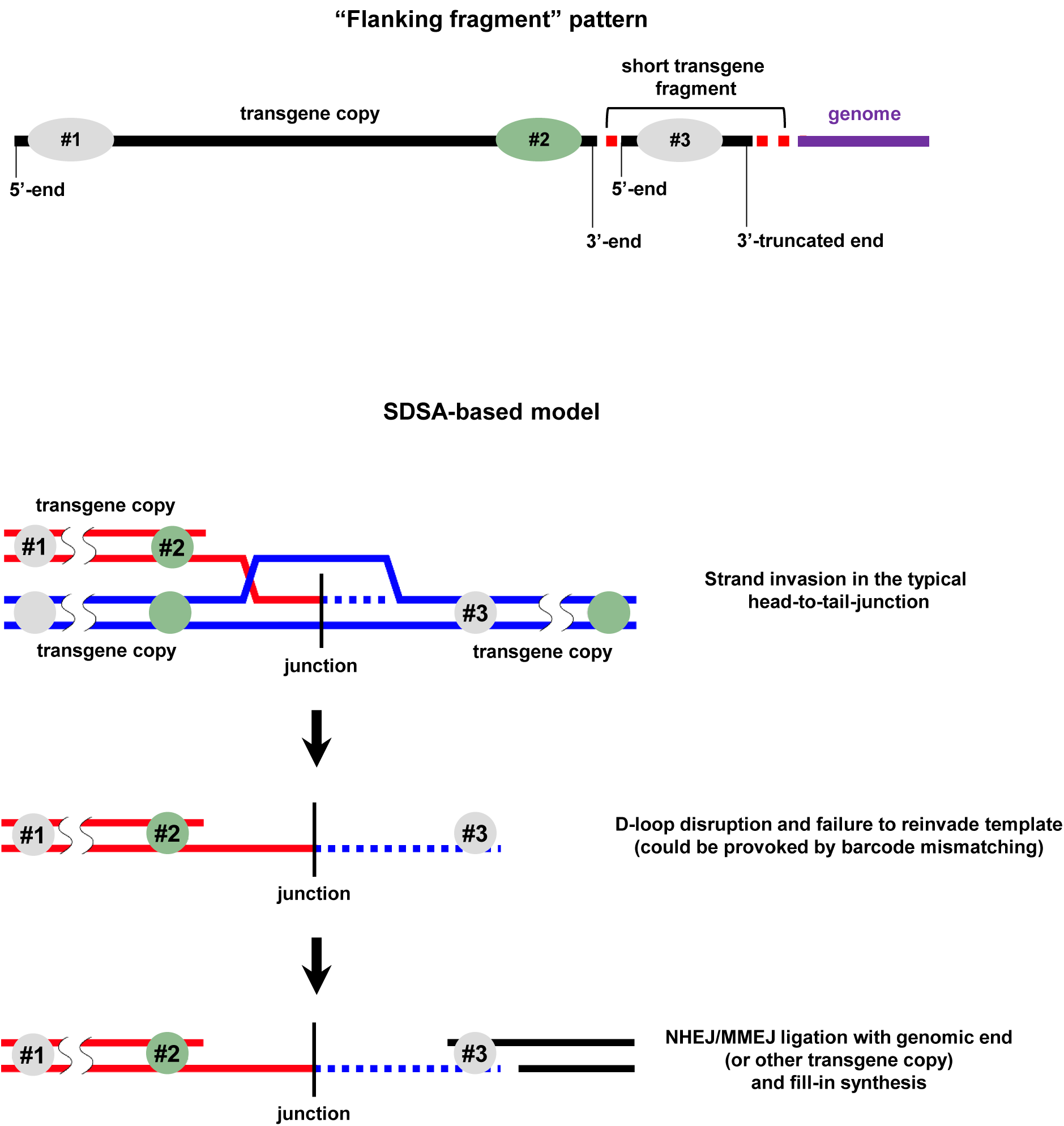
Model for “flanking fragment” pattern emergence. This pattern is frequently observed in concatemers (our data; Hamada et al., 1993; Yan et al., 2010; other reports). For example, short 5’-terminal fragment of transgene could be found near 3’-end of another full-size copy. This is especially prominent at the transgene-genome borders (internal transgene-transgene junctions are more difficult to interpret). We have detected 2 independent cases of this pattern at genome borders (embryos #5, #10) and several cases of small duplication of internal concatemer junction fragments (embryos #2, #6).

**Supplementary Figure 20.**
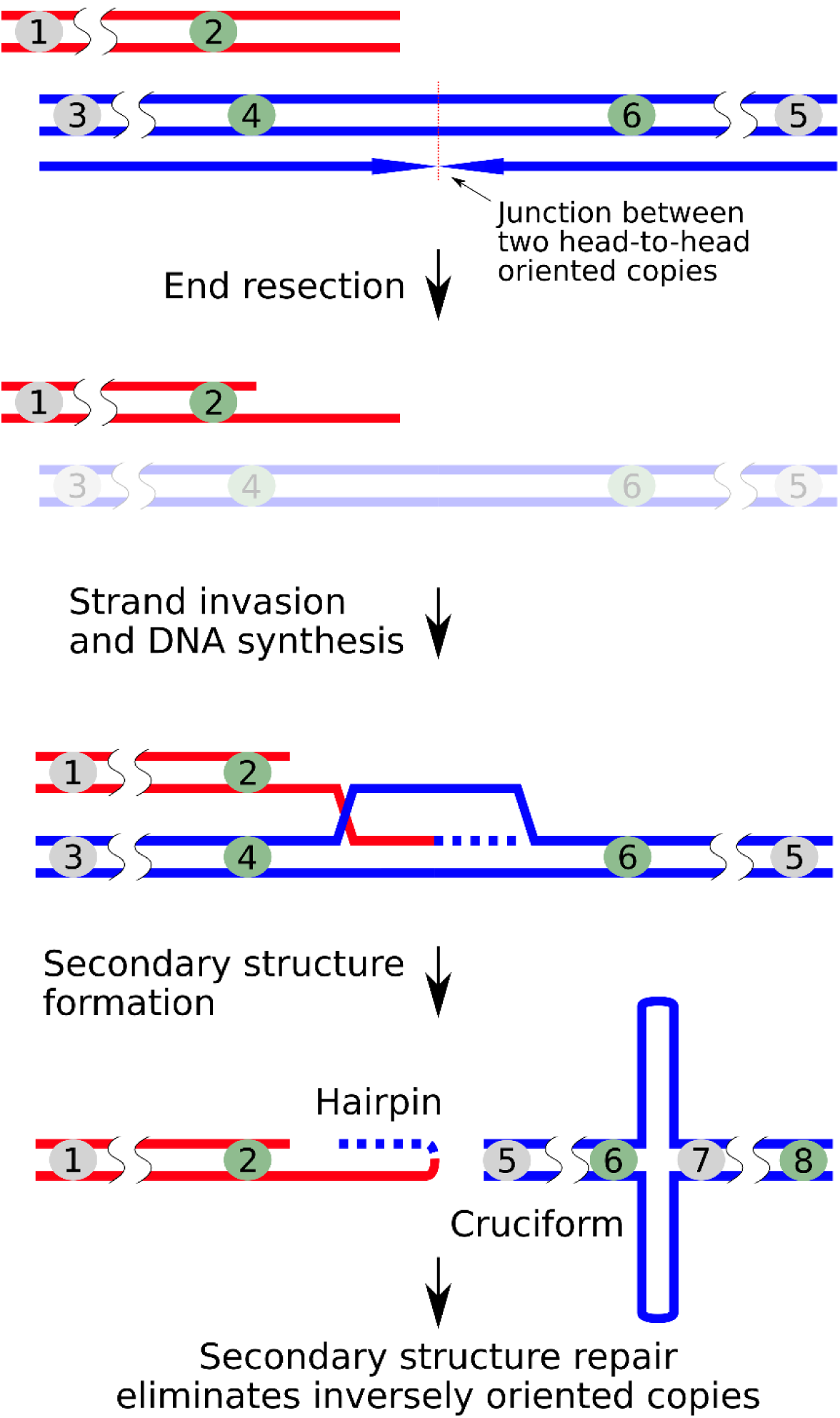
HR-based model for selective palindrome wiping out. Destabilization of a palindromic junction by filament invasion leads to exposure of ssDNA and hairpin formation.

**Supplementary Figure 21.**
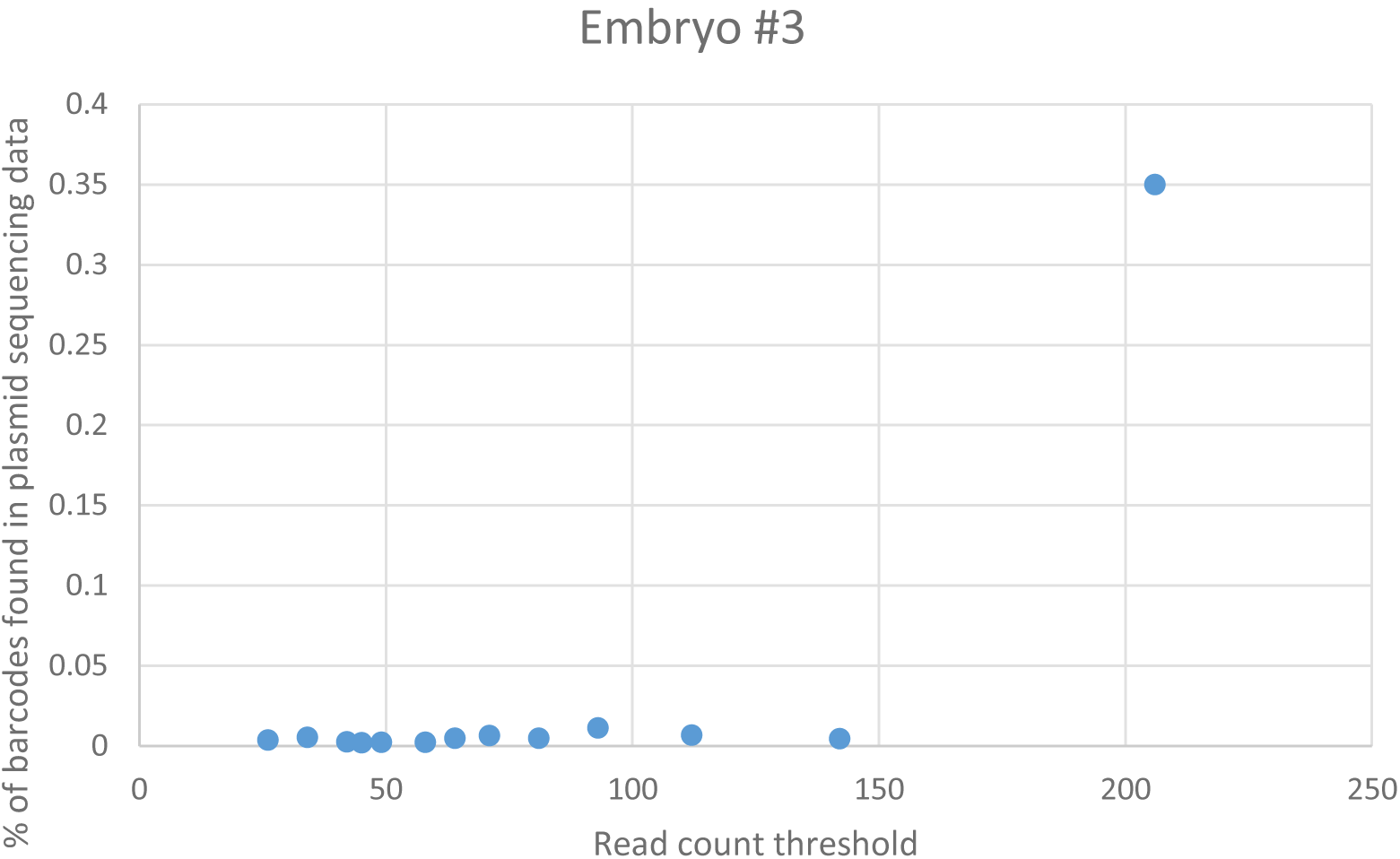
Setting threshold for barcode filtering based on the plasmid library/embryo data overlap (embryo #3) (see Methods for explanation).

**Supplementary Table 1.** List of publications documenting pronuclear microinjection outcomes, with focus on transgene copy number and copy orientations.

| Publication | Transgene | Methods | Copy number and orientation (H/T – head or tail junction) | Integration site structure (transgene-genome borders) |
| --- | --- | --- | --- | --- |
| Westendorf et al., 2016 | B6·Cg-Tg(Cd4-cre)1Cwi (Cre recombinase under Cd4-promoter)<br>Transgene size – 15,6Kbp | WGS | 15-16 copies, 10 copies lack 3'-end region<br>No data about orientation | 1. Genomic deletion (5,6Kb) (chr3:60335693-60341285)<br>2. Small flanking inversion at the 5'-end (chr3:60335695- 60335808) |
| Harkins, Whitton, 2016 | MerCreMer (tamoxifen-inducible Cre recombinase under Myh6-promoter)<br>Transgene size – 6,5Kbp | SKYFISH<br>Genome Walking | Single copy, but with multiple flanking transgene fragments at both 5'- and 3'-end (attributed to the “transgene chromotripsis”) | Genomic deletion (19,5Kb) (chr19:31852892-31872391) |
| Masumura et al., 2015 | gpt delta (Lambda phage shuttle vector EG10), two transgenic lines: mouse and rat<br>Transgene size – 48Kb | WGS<br>Southern Blot<br>FISH | Mouse line: ~41 H-T, 1 H-H, 1 T-T, ~14 fragmented copies<br>Rat line: ~15 H-T, 1 T-T, ~4 fragmented copies | Mouse line:<br>1. Integration at Chr17:40878810-40878879<br>2. Duplication of the flanking genomic region (69bp)<br>3. 5'-border: 2bp microhomology; 3'-border: 5 bp microhomology<br>4. Inverted flanking transgene copies<br>Rat line:<br>1. Integration at Chr4:79828427-79900397<br>2. Genomic deletion (72Kb)<br>3. 5'-border: no homology; 3'-border: 14 bp insertion (10bp microhomology)<br>4. Inverted flanking transgene copies |
| Yong et al., 2015 | WAP-Her2<br>Linearized with HindIII<br>Transgene size – 7Kb | WGS | ~162 H-T; H-H or T-T not observed | 1. Integration at Chr5:150719804–150719794<br>2. Duplication of the flanking genomic region (10bp) |
| Yan et al., 2013 | 7 different transgenes<br>38 integration sites<br>Transgene size – 2-6Kb | TAIL-PCR | Not examined | 1. Frequent integration in the coding regions (17 of 38 sites)<br>2. Frequent microhomologies (1-19bp) at the borders (76% in 30 mouse lines) |
| Chiang et al., 2012 | HTT: 2 transgenes in mouse (1,9Kb) and sheep (11,6Kb) | Biotin Target Enrichment + NGS | 2 mouse lines<br>5 sheep lines | Extensive information about 7 genomic loci (deletions, translocations) |
|  |  |  | Multiple rearrangements and mostly random orientation of copies |  |
| Shwed et al., 2010 | MutaMouse lgt10-lacZ<br>(Lambda phage shuttle vector)<br>Transgene size – 48Kb | qPCR Southern Blot | ~30 H-T, several inverted copies | Not examined |
| Yan et al., 2010 | CMV-CBA-fad2<br>Linearized with SalI and BamHI<br>Transgene size – 3,6Kb | TAIL-PCR | H-T (not examined in detail) | 1. Integration at ChrX: 87462177-87507732<br>2. Genomic deletion (45,6Kb)<br>3. 5'-border: 3bp microhomology + 330 bp insertion (transgene fragment); 3'-border: 7bp microhomology + 29 bp insertion (transgene fragment) |
| Suzuki et al., 2006 | Ganglioside GM1 synthase cDNA (pCAGGS backbone)<br>Linearized with SalI and HindIII<br>Transgene size – 2,5Kb | Genome Walking | At least 3 copies + fragments in random orientation: 2 H-T, 1 T-T | 1. Integration at Chr1<br>2. 5'-border: 11 bp lost; 3'-border: 7 bp lost<br>3. Inverted flanking transgene copies |
| Licence et al., 2003 | VpreB1-CD122<br>Linearized with HindIII<br>Transgene size – 3,7Kb | qPCR Southern Blot | 11 mouse lines: 6 lines - 2-7 copies; 3 lines - 10-12 copies, 1 line - 33 copies<br>All copies have H-T orientation | Not examined |
| Suemizu et al., 2002 | Tg-rasH2<br>Linearized with BamHI<br>Transgene size – 7Kb | Southern Blot | 3 H-T, no deletions between copies (religated at BamHI site) | 1. Genomic deletion (1,8Kb)<br>2. 5'-border: 4bp microhomology + 148 bp transgene deletion; 3'-border: 10bp microhomology + 24 bp transgene deletion |
| Tacken et al., 2001 | VLDLR gene<br>Linearized with NotI<br>Transgene size – 35Kb + 40Kb (overlapping cosmid inserts) | Southern Blot<br>Fiber-FISH | Line 1: >44 copies, (>90% H-T)<br>Line 2: >64 copies (>90% H-T) | Not examined |
| Koetsier et al., 1996 | pAd2E2AL-CAT<br>Linearized with NdeI + XmnI<br>Transgene size – 3,3Kb | Southern Blot | Line 7-1: 4 H-T and 6 H-T at separate sites<br>Line 5-8: 4 H-T<br>Line 8-1: 20 copies | Not examined |
| McFarlane, Wilson, 1996 | PyLMP<br>Linearized with BamHI + EcoRI<br>Transgene size – 3,8Kb | Southern Blot | 9 mouse lines: 2-10 copies H-T | One of the lines (2 H-T copies) was examined:<br>1. No deletion in genome<br>2. 5'-border: 7 bp at transgene end lost; 3'-border: no loss<br>3. Transgene-transgene junction with 4 bp loss (EcoRI site) |
| Pawlik et al., 1995 | HS2/beta-globin<br>Linearized with KpnI + SalI<br>Transgene size – 5,2Kb | Southern Blot | 8 mouse lines: 10-100 mostly H-T,<br>but many lines also have T-T<br>junctions | 1. Many transgene-transgene junctions sequenced:<br>minor deletion or fill-in synthesis of restricted ends |
| Hamada et al., 1993 | Metallothionein-I mouse<br>promoter + human TTR gene<br>(pUC backbone)<br>Linearized with blunt ends<br>Transgene size – 10,8Kb | Southern Blot<br>Inverse PCR | Line MPA434: single copy, with<br>short 3'-end fragment at the 5'-end of<br>the full-sized copy<br>MPA551 2: 3 H-T | Line MPA434:<br>1. Genomic deletion (32bp)<br>2. 5'- and 3'-borders: 3bp microhomology<br>Line MPA551:<br>1. Genomic deletion (10Kb)<br>2. 5'- and 3'-borders: 1-2bp microhomology<br>3. 2-3bp deletions at transgene-transgene junctions |
| Mark et al., 1992 | LambdaRbetaG2<br>Transgene size – 50Kb | Southern Blot<br>Phage Library | 6-8 H-T | 1. Genomic deletion (22Kb)<br>2. 5'-border: 21bp unknown sequence cointegration;<br>3'-border: 57bp fragment of lambda vector |
| Burdon et al., 1991 | WAP gene<br>Linearized with EcoRI<br>Transgene size – 7,2Kb | qPCR | 6 lines: 3-10 H-T | Not examined |
| Radice et al., 1991 | Human B- globin gene<br>Linearized with HindIII<br>Transgene size – 7,7Kb | Southern Blot<br>Phage Library | 10-20 H-T | 1. Genomic deletion (2-3Kb)<br>2. Cointegration of a small genomic fragment |
| Rohan et al., 1990 | RSVcat gene<br>Linearized with PstI + NdeI<br>Transgene size – 2,7Kb | Southern Blot | 8 lines:<br>6 lines - only fragments (0 copies)<br>1 line - 2-5 H-T<br>1 line - 25 H-T (but only two variants<br>of junctions) | 1. Nibbled ends at the transgene-transgene junctions<br>(1-62bp) |
| Mahon et al., 1988<br>Overbeek et al., 1986 | pRSV-CAT | Southern Blot<br>FISH | 9 lines:<br>2 lines - only fragments (0 copies)<br>4 lines - 2-13 H-T<br>3 lines - 31-180 H-T | 1. В одной из линий (50 копий) интеграция<br>привела к транслокации большого фрагмента<br>плеча 6 хромосомы на 17. |
| Wilkie,<br>Palmiter, 1987 | pMK fusion gene in pBR322<br>Linearized with BamHI<br>Transgene size – ~7Kb | Southern Blot<br>Phage Library | 4 copies: 2 T-T + 2 fragmented<br>copies | 1. Duplication 5Kb at the borders<br>2. Cointegration of 525bp genomic fragment |
| Covarrubias et al., 1986<br>Wagner et al., 1983 | phGH plasmid<br>Transgene size – 4,3Kb | Southern Blot | 2-5 H-T, some copies fragmented | 1. Genomic deletion (10Kb) |
| Khillan et al., 1985 | pPi25.1 (*Drosophila* P-element)<br>Linearized with XbaI<br>Transgene size – ~9Kb | Southern Blot | Line 1: single copy<br>Lines 2-3: 25-30 H-T | 1. In Line 1 self-ligated transgene was linearized<br>again at random point before integration |
| van der Putten et al., 1984 | pHIGfbRI<br>Circular DNA<br>Transgene size – ~17Kb | Southern Blot | 3 H-T, 1 H-H, 1T-T | Not examined |
| Brinster et al., 1983 | Immunoglobulin gene VkM.21<br>Linearized with BglI + XhoI<br>Transgene size – ~15Kb | Southern Blot | 3 lines: 20, 50, 95+40 (two sites) H-T | Not examined |
| McKnight et al., 1983 | Chicken transferrin gene<br>Linearized with BglI + XhoI or circular<br>Transgene size – ~17Kb | Southern Blot<br>Dot Blot | Circular transgene - 3 lines (1, 11, 12 copies)<br>Linearized transgene - 4 lines (2, 11, 11, 14 copies)+ 4 lines with fragments only<br>Of 6 lines analyzed all are H-T | Not examined |
| Wagner et al., 1983 | phGH plasmid<br>Transgene size – 4,3Kb | Southern Blot | 6 lines: 0, 1, 4, 4, 8, 20 H-T with flanking fragments | Not examined |
| Palmiter et al., 1982 | MT-1 promoter + HSV TK<br>Linearized with BamHI<br>Transgene size – ~7Kb | Southern Blot<br>Dot Blot | 7 lines: 1, 2, 2, 8, 32, 100, 150 H-T | Not examined |
| Palmiter et al., 1982 | MT-1 promoter + rat GH<br>Linearized with BamHI<br>Transgene size – ~5Kb | Southern Blot | 7 lines: 1, 2, 2, 8, 10, 20, 35 H-T | Not examined |

**Supplementary Table 2.**
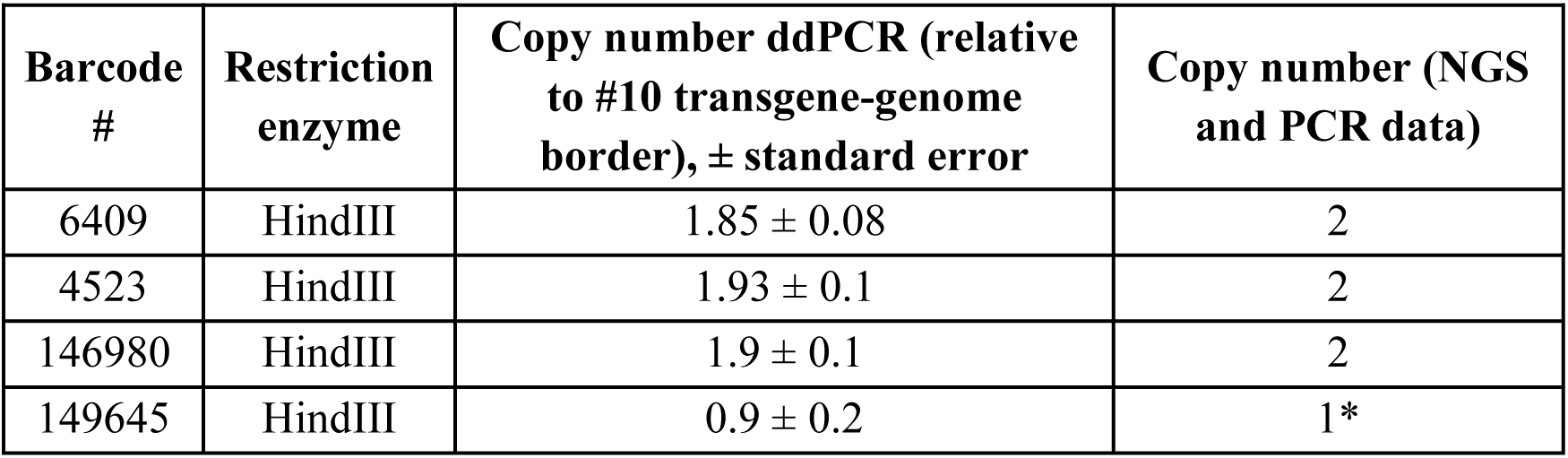
Calculating barcodes in embryo #10. Barcode flagged with asterisk actually has two copies, but one of them is unaccounted by PCR due to deletion of the primer site.

| Barcode # | Restriction enzyme | Copy number ddPCR (relative to #10 transgene-genome border), ± standard error | Copy number (NGS and PCR data) |
| --- | --- | --- | --- |
| 6409 | HindIII | $1.85 \pm 0.08$ | 2 |
| 4523 | HindIII | $1.93 \pm 0.1$ | 2 |
| 146980 | HindIII | $1.9 \pm 0.1$ | 2 |
| 149645 | HindIII | $0.9 \pm 0.2$ | 1* |

**Supplementary Table 3.**
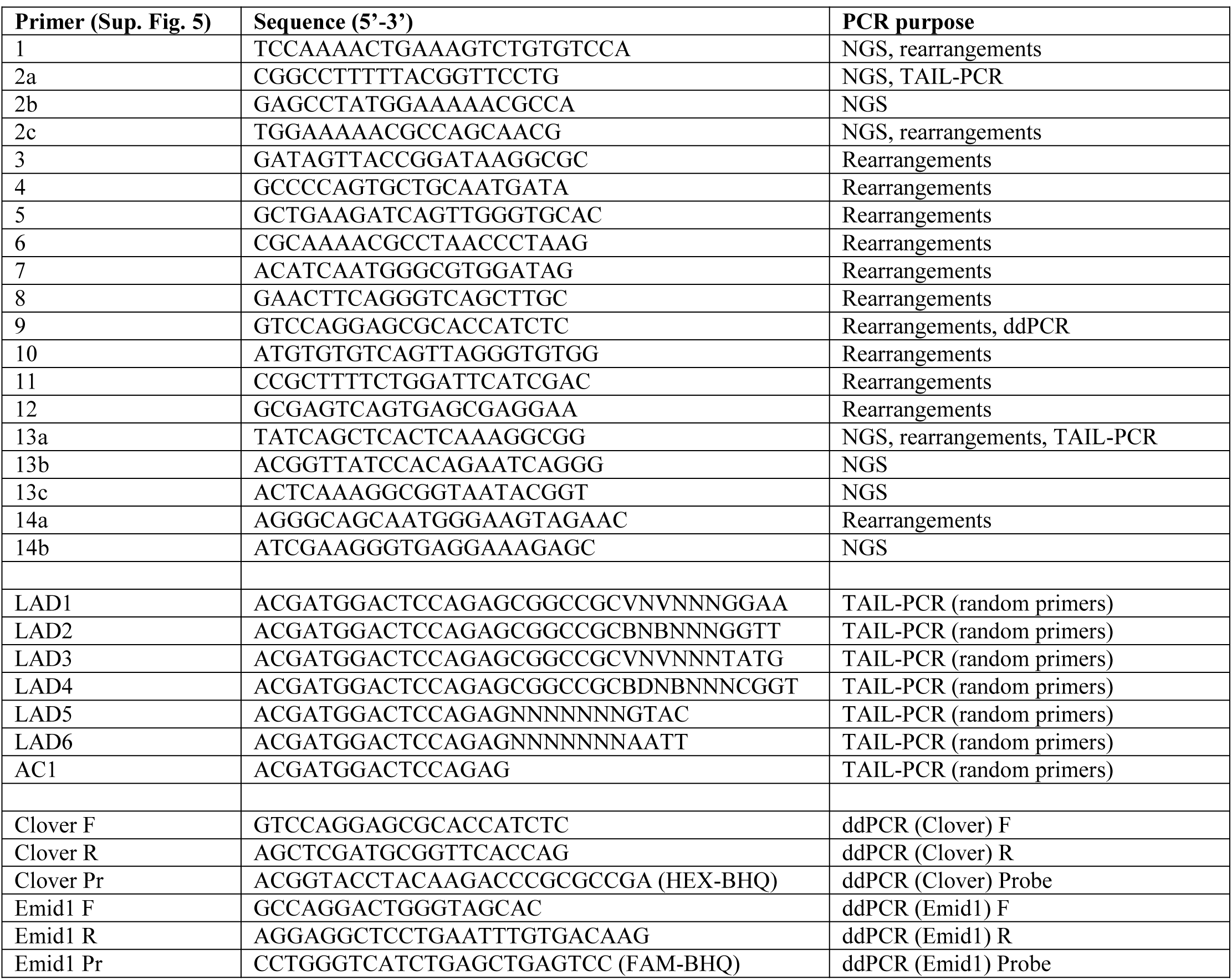
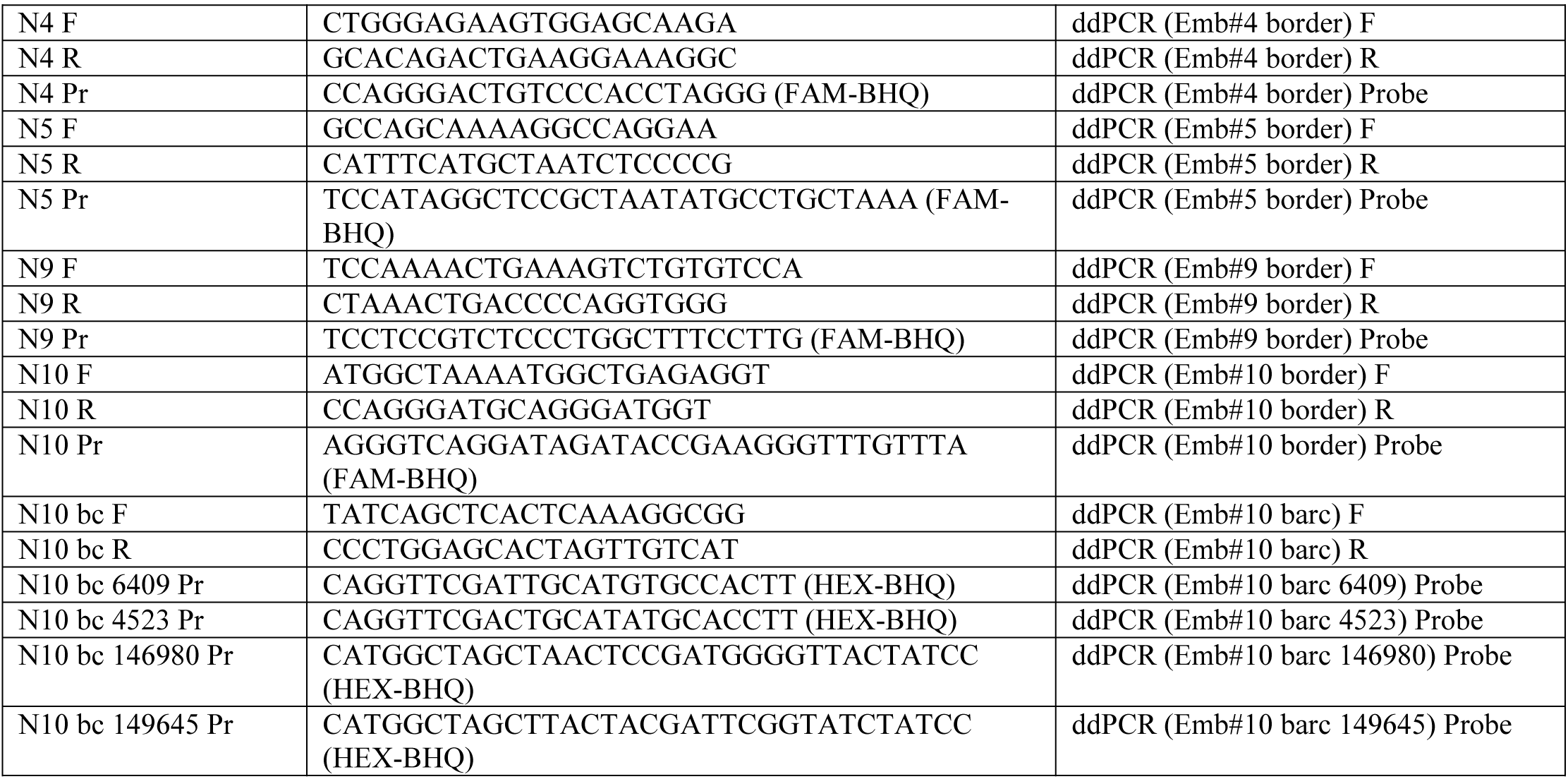
PCR primers and ddPCR probes used in the study.

**Supplementary Table 4.**
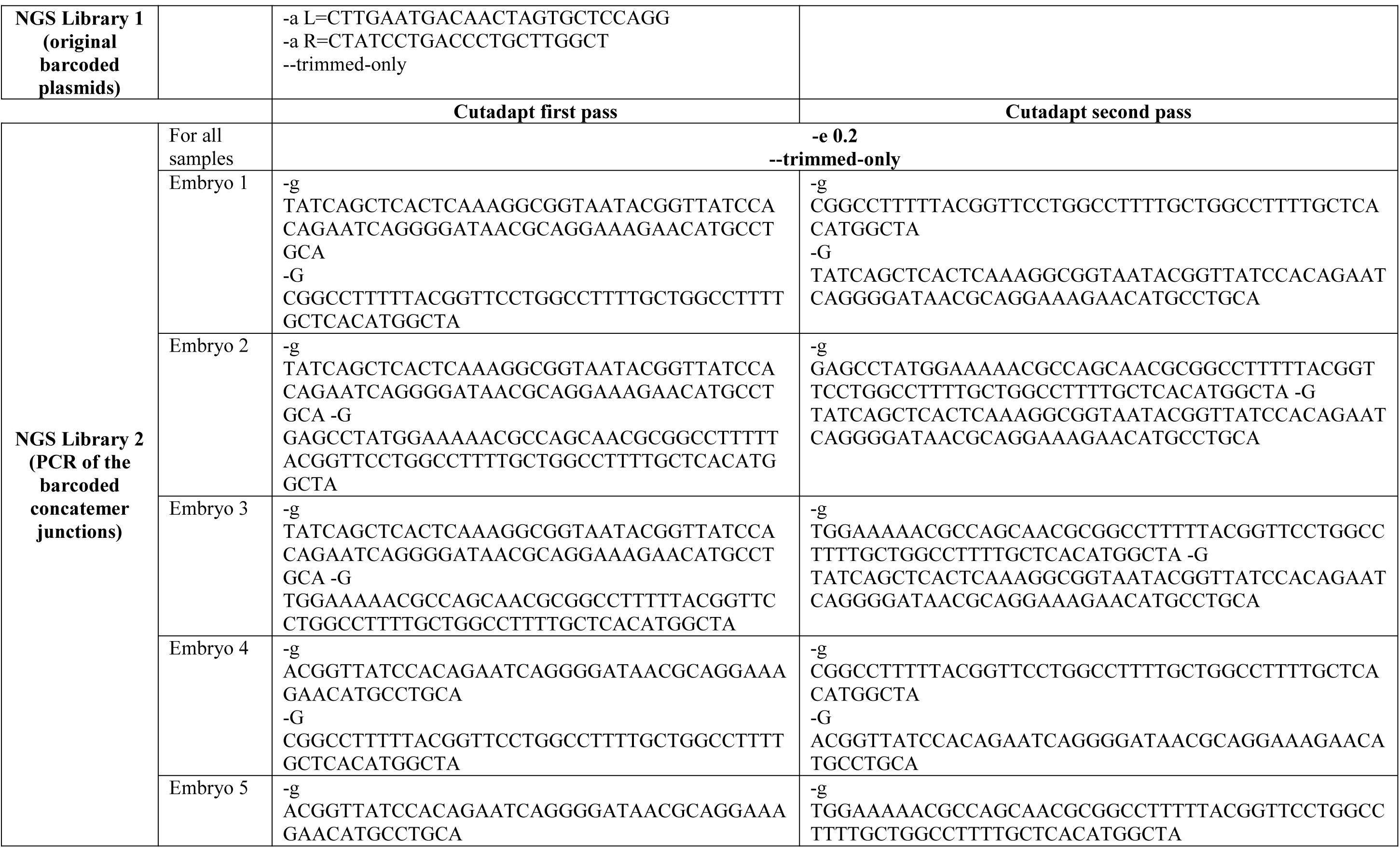
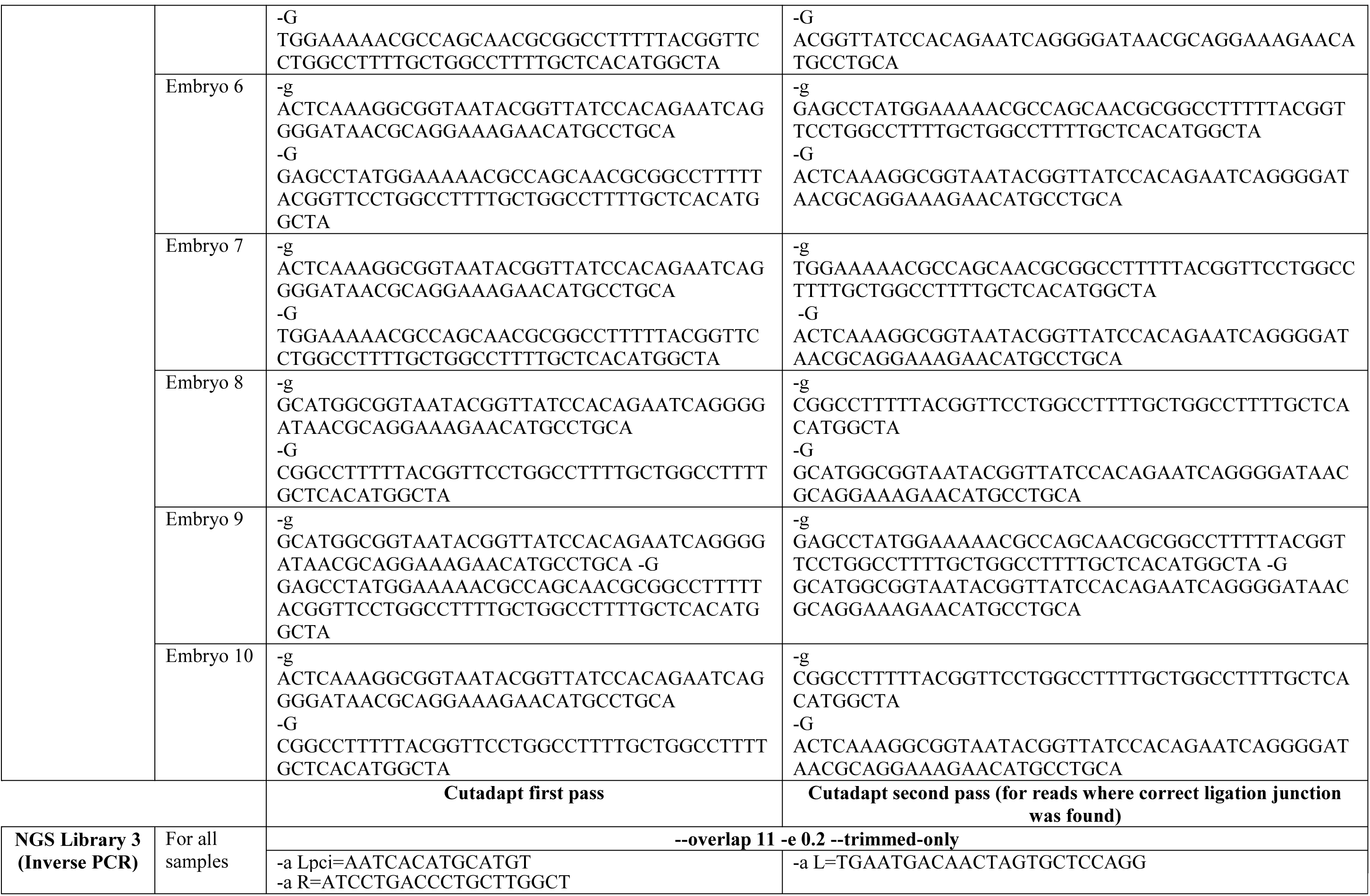
Cutadapt options used for reads trimming.

**Supplementary Table 5.** Barcode patterns.

| Pattern description | Python RE-pattern (5'-barcode sequence) | Python RE-pattern (3'-barcode sequence) |
| --- | --- | --- |
| Pattern 1<br>Exact match | [ATGC]{2}CGA[ATGC]{2}GCA[ATGC]{2}TGC[ATGC]{2}<br>Example of matching pattern:<br>AT **CGA** AG **GCA** GG **TGC** CC | [ATGC]{2}ACT[ATGC]{2}GAT[ATGC]{2}GGT[ATGC]{2} |
| Pattern 2<br>+/- 1 redundant base | [ATGC]{1,3}CGA[ATGC]{1,3}GCA[ATGC]{1,3}TGC[ATGC]{1,3}<br>Example of matching patten:<br>AT **CGA** A**C**G **GCA** GG **TGC** CC | [ATGC]{1,3}ACT[ATGC]{1,3}GAT[ATGC]{1,3}GGT[ATGC]{1,3} |
| Pattern 3<br>SNP (1 bp insertion,<br>deletion or substitution)<br>in constant bases | 75 different patterns<br>Example of matching pattern:<br>AT **CGGA** AG **GCA** GG **TGC** CC | 74 different patterns |
| Both pattern 2 and<br>pattern 3 | 75 different patterns<br>Example of matching pattern:<br>AT **CGGA** AG **GCA** **G** **TGC** CC | 74 different patterns |

**Supplementary Table 6.** Barcode pair thresholds for filtering.

| Library | Sample | Count of reads supporting pair threshold |
| --- | --- | --- |
| NGS Library 1 (original barcoded plasmids) | Original barcoded plasmids | 1 |
| NGS Library 2 (PCR of the barcoded concatemer junctions) | Embryo 1 | 1 |
|  | Embryo 2 | 50 |
|  | Embryo 3 | 250 |
|  | Embryo 4 | 2 000 |
|  | Embryo 5 | 500 |
|  | Embryo 6 | 3 000 |
|  | Embryo 7 | 2 |
|  | Embryo 8 | 5 000 |
|  | Embryo 9 | 200 |
|  | Embryo 10 | 1 |
| NGS Library 3 (Inverse PCR) | Embryo 1 | 10 000 |
|  | Embryo 2 | 300 |
|  | Embryo 3 | 500 |
|  | Embryo 4 | 10 000 |
|  | Embryo 5 | 1 |
|  | Embryo 6 | 30 000 |
|  | Embryo 7 | 300 |
|  | Embryo 8 | 3000 |
|  | Embryo 9 | 10 000 |
|  | Embryo 10 | 5000 |

## References

1. Savić, N. and Schwank, G. (2016) Advances in therapeutic CRISPR/Cas9 genome editing. Transl. Res., 10.1016/j.trsl.2015.09.008.

2. Bertolini, L.R., Meade, H., Lazzarotto, C.R., Martins, L.T., Tavares, K.C., Bertolini, M. and Murray, J.D. (2016) The transgenic animal platform for biopharmaceutical production. Transgenic Res., 10.1007/s11248-016-9933-9.

3. Kamthan, A., Chaudhuri, A., Kamthan, M. and Datta, A. (2016) Genetically modified (GM) crops: milestones and new advances in crop improvement. Theor. Appl. Genet., 10.1007/s00122-016-2747-6.

4. Chang, H.H.Y., Pannunzio, N.R., Adachi, N. and Lieber, M.R. (2017) Non-homologous DNA end joining and alternative pathways to double-strand break repair. Nat. Rev. Mol. Cell Biol., 10.1038/nrm.2017.48.

5. Ranjha, L., Howard, S.M. and Cejka, P. (2018) Main steps in DNA double-strand break repair: an introduction to homologous recombination and related processes. Chromosoma, 10.1007/s00412-017-0658-1.

6. Piazza, A. and Heyer, W.D. (2019) Homologous Recombination and the Formation of Complex Genomic Rearrangements. Trends Cell Biol., 10.1016/j.tcb.2018.10.006.

7. Low, B.E., Kutny, P.M. and Wiles, M. V. (2016) Simple, efficient CRISPR-cas9-mediated gene editing in mice: Strategies and methods. Methods Mol. Biol., 10.1007/978-1-4939-3661-8_2.

8. Jiang, F. and Doudna, J.A. (2017) CRISP[1] F. Jiang und J. A. Doudna, „CRISPR–Cas9 Structures and Mechanisms“, Annu. Rev. Biophys., Bd. 46, Nr. 1, S. 505–529, Mai 2017.R–Cas9 Structures and Mechanisms. Annu. Rev. Biophys., 10.1146/annurev-biophys-062215-010822.

9. Ben Jehuda, R., Shemer, Y. and Binah, O. (2018) Genome Editing in Induced Pluripotent Stem Cells using CRISPR/Cas9. Stem Cell Rev. Reports, 10.1007/s12015-018-9811-3.

10. Nishiyama, J. (2018) Genome editing in the mammalian brain using the CRISPR–Cas system. Neurosci. Res., 10.1016/j.neures.2018.07.003.

11. Pawelczak, K.S., Gavande, N.S., VanderVere-Carozza, P.S. and Turchi, J.J. (2018) Modulating DNA Repair Pathways to Improve Precision Genome Engineering. ACS Chem. Biol., 10.1021/acschembio.7b00777.

12. Charpentier, M., Khedher, A.H.Y., Menoret, S., Brion, A., Lamribet, K., Dardillac, E., Boix, C., Perrouault, L., Tesson, L., Geny, S., et al. (2018) CtIP fusion to Cas9 enhances transgene integration by homology-dependent repair. Nat. Commun., 10.1038/s41467-018-03475-7.

13. Auerbach, A.B. (2004) Production of functional transgenic mice by DNA pronuclear microinjection. Acta Biochim. Pol.

14. Folger, K.R., Wong, E.A., Wahl, G. and Capecchi, M.R. (1982) Patterns of integration of DNA microinjected into cultured mammalian cells: evidence for homologous recombination between injected plasmid DNA molecules. Mol. Cell. Biol., 2, 1372–1387.

15. Brinster, R.L., Chen, H.Y., Trumbauer, M.E., Yagle, M.K. and Palmiter, R.D. (1985) Factors Affecting the Efficiency of Introducing Foreign DNA into Mice by Microinjecting Eggs. Proc. Natl. Acad. Sci. U. S. A., 10.1073/pnas.82.13.4438.

16. Lu, R., Neff, N.F., Quake, S.R. and Weissman, I.L. (2011) Tracking single hematopoietic stem cells in vivo using high-throughput sequencing in conjunction with viral genetic barcoding. Nat. Biotechnol., 29, 928–33.

17. Yunusova, A.M., Fishman, V.S., Vasiliev, G. V. and Battulin, N.R. (2017) Deterministic versus stochastic model of reprogramming: New evidence from cellular barcoding technique. Open Biol., 10.1098/rsob.160311.

18. Clarke, A.R., Brown, G.A.J. and Corbin, T.J. (2003) Oocyte Injection in the Mouse. In Transgenesis Techniques.

19. Serova, I.A., Dvoryanchikov, G.A., Andreeva, L.E., Burkov, I.A., Dias, L.P.B., Battulin, N.R., Smirnov, A. V. and Serov, O.L. (2012) A 3,387 bp 5′-flanking sequence of the goat alpha- S1-casein gene provides correct tissue-specific expression of human granulocyte colony-stimulating factor (hG-CSF) in the mammary gland of transgenic mice. Transgenic Res., 10.1007/s11248-011-9547-1.

20. Potapov, V. and Ong, J.L. (2017) Examining sources of error in PCR by single-molecule sequencing. PLoS One, 10.1371/journal.pone.0169774.

21. Martin, M. and N,T. (2011) Cutadapt removes adapter sequences from high-throughput sequencing reads. EMBnet.journal.

22. Codner, G.F., Lindner, L., Caulder, A., Wattenhofer-Donzé, M., Radage, A., Mertz, A., Eisenmann, B., Mianné, J., Evans, E.P., Beechey, C. V., et al. (2016) Aneuploidy screening of embryonic stem cell clones by metaphase karyotyping and droplet digital polymerase chain reaction. BMC Cell Biol., 10.1186/s12860-016-0108-6.

23. Burkov, I.A., Serova, I.A., Battulin, N.R., Smirnov, A. V., Babkin, I. V., Andreeva, L.E., Dvoryanchikov, G.A. and Serov, O.L. (2013) Expression of the human granulocyte-macrophage colony stimulating factor (hGM-CSF) gene under control of the 5′-regulatory sequence of the goat alpha-S1-casein gene with and without a MAR element in transgenic mice. Transgenic Res., 10.1007/s11248-013-9697-4.

24. Rohan, R.M., King, D. and Frels, W.I. (1990) Direct sequencing of PCR-amplified junction fragments from tandemly repeated transgenes. Nucleic Acids Res., 10.1093/nar/18.20.6089.

25. Krupovic, M. and Forterre, P. (2015) Single-stranded DNA viruses employ a variety of mechanisms for integration into host genomes. Ann. N. Y. Acad. Sci., 10.1111/nyas.12675.

26. Tomaska, L., Nosek, J., Kramara, J. and Griffith, J.D. (2009) Telomeric circles: Universal players in telomere maintenance. Nat. Struct. Mol. Biol., 10.1038/nsmb.1660.

27. Chen, X.J. and Clark-Walker, G.D. (2018) Unveiling the mystery of mitochondrial DNA replication in yeasts. Mitochondrion, 10.1016/j.mito.2017.07.009.

28. Bishop, J.O. (1996) Chromosomal insertion of foreign DNA. Reprod Nutr Dev, S0926528797818850 [pii].

29. Stone, J.E., Ozbirn, R.G., Petes, T.D. and Jinks-Robertson, S. (2008) Role of proliferating cell nuclear antigen interactions in the mismatch repair-dependent processing of mitotic and meiotic recombination intermediates in yeast. Genetics, 10.1534/genetics.107.085415.

30. Chakraborty, U., George, C.M., Lyndaker, A.M. and Alani, E. (2016) A delicate balance between repair and replication factors regulates recombination between divergent DNA sequences in Saccharomyces cerevisiae. Genetics, 10.1534/genetics.115.184093.

31. Schimmel, J., Kool, H., van Schendel, R. and Tijsterman, M. (2017) Mutational signatures of non-homologous and polymerase theta-mediated end-joining in embryonic stem cells. EMBO J., 10.15252/embj.201796948.

32. Daley, J.M., Gaines, W.A., Kwon, Y. and Sung, P. (2014) Regulation of DNA Pairing in Homologous Recombination. Cold Spring Harb. Perspect. Biol., 10.1101/cshperspect.a017954.

33. Wright, W.D., Shah, S.S. and Heyer, W.D. (2018) Homologous recombination and the repair of DNA double-strand breaks. J. Biol. Chem., 10.1074/jbc.TM118.000372.

34. Hamada, T., Sasaki, H., Seki, R. and Sakaki, Y. (1993) Mechanism of chromosomal integration of transgenes in microinjected mouse eggs: sequence analysis of genome-transgene and transgene-transgene junctions at two loci. Gene, 10.1016/0378-1119(93)90563-I.

35. Yan, B., Li, D. and Gou, K. (2010) Homologous illegitimate random integration of foreign DNA into the X chromosome of a transgenic mouse line. BMC Mol. Biol., 10.1186/1471-2199-11-58.

36. Verma, P. and Greenberg, R.A. (2016) Noncanonical views of homology-directed DNA repair. Genes Dev., 10.1101/gad.280545.116.

37. Masumura, K., Sakamoto, Y., Kumita, W., Honma, M., Nishikawa, A. and Nohmi, T. (2015) Genomic integration of lambda EG10 transgene in gpt delta transgenic rodents. Genes Environ., 10.1186/s41021-015-0024-6.

38. Chiang, C., Jacobsen, J.C., Ernst, C., Hanscom, C., Heilbut, A., Blumenthal, I., Mills, R.E., Kirby, A., Lindgren, A.M., Rudiger, S.R., et al. (2012) Complex reorganization and predominant non-homologous repair following chromosomal breakage in karyotypically balanced germline rearrangements and transgenic integration. Nat. Genet., 10.1038/ng.2202.

39. Lutz, C., Davis, T.L., Urban, R., Chesler, E.J., He, H., Murray, S.A., Ndukum, J., Braun, R.E., van Min, M., Goodwin, L.O., et al. (2019) Large-scale discovery of mouse transgenic integration sites reveals frequent structural variation and insertional mutagenesis. Genome Res., 10.1101/gr.233866.117.

40. Matsuda, J., Yamamoto, Y., Noguchi, Y., Takekawa, N., Koura, M., Hata, T., Suzuki, O., Takano, K. and Uchio-Yamada, K. (2006) Transgene Insertion Pattern Analysis Using Genomic Walking in a Transgenic Mouse Line. Exp. Anim., 10.1538/expanim.55.65.

41. Akgun, E., Zahn, J., Baumes, S., Brown, G., Liang, F., Romanienko, P.J., Lewis, S. and Jasin, M. (1997) Palindrome resolution and recombination in the mammalian germ line. Mol. Cell. Biol., 10.1103/PhysRevSTAB.14.060703.

42. Chen, H., Lisby, M. and Symington, L. (2013) RPA Coordinates DNA End Resection and Prevents Formation of DNA Hairpins. Mol. Cell, 10.1016/j.molcel.2013.04.032.

43. Grunwald, H.A., Gantz, V.M., Poplawski, G., Xu, X.R.S., Bier, E. and Cooper, K.L. (2019) Super-Mendelian inheritance mediated by CRISPR–Cas9 in the female mouse germline. Nature, 10.1038/s41586-019-0875-2.

44. Tacken, P.J., Zee, A. V.D., Beumer, T.L., Florijn, R.J., Gijpels, M.J.J., Havekes, L.M., Frants, R.R., Dijk, K.W.V. and Hofker, M.H. (2001) Effective generation of very low density lipoprotein receptor transgenic mice by overlapping genomic DNA fragments: High testis expression and disturbed spermatogenesis. Transgenic Res., 10.1023/A:1016682520887.

45. Skryabin, B. V., Gubar, L., Seeger, B., Kaiser, H., Stegemann, A., Roth, J., Meuth, S.G., Pavenstadt, H., Sherwood, J., Pap, T., et al. (2019) Pervasive head-to-tail insertions of DNA templates mask desired CRISPR/Cas9-mediated genome editing events. bioRxiv, 10.1101/570739.

## REFERENCES

Westendorf K, Durek P, Ayew S, Mashreghi MF, Radbruch A. Chromosomal localisation of the CD4cre transgene in B6·Cg-Tg(Cd4-cre)1Cwi mice. J Immunol Methods. 2016 Sep;436:54–7. doi: 10.1016/j.jim.2016.06.005. Epub 2016 Jun 23.

Harkins S, Whitton JL. Chromosomal mapping of the αMHC-MerCreMer transgene in mice reveals a large genomic deletion. Transgenic Res. 2016 Oct;25(5):639–48. doi: 10.1007/s11248-016-9960-6. Epub 2016 May 10.

Masumura K, Sakamoto Y, Kumita W, Honma M, Nishikawa A, Nohmi T. Genomic integration of lambda EG10 transgene in gpt delta transgenic rodents. Genes Environ. 2015 Dec 1;37:24. doi: 10.1186/s41021-015-0024-6.

Yong CS, Sharkey J, Duscio B, Venville B, Wei WZ, Jones RF, Slaney CY, Mir Arnau G, Papenfuss AT, Schröder J, Darcy PK, Kershaw MH. Embryonic Lethality in Homozygous Human Her-2 Transgenic Mice Due to Disruption of the Pds5b Gene. PLoS One. 2015 Sep 3;10(9):e0136817. doi: 10.1371/journal.pone.0136817.

Yan BW, Zhao YF, Cao WG, Li N, Gou KM. Mechanism of random integration of foreign DNA in transgenic mice. Transgenic Res. 2013 Oct;22(5):983–92. doi: 10.1007/s11248-013-9701-z.

Chiang C, Jacobsen JC, Ernst C, Hanscom C, Heilbut A, Blumenthal I, Mills RE, Kirby A, Lindgren AM, Rudiger SR, McLaughlan CJ, Bawden CS, Reid SJ, Faull RL, Snell RG, Hall IM, Shen Y, Ohsumi TK, Borowsky ML, Daly MJ, Lee C, Morton CC, MacDonald ME, Gusella JF, Talkowski ME. Complex reorganization and predominant non-homologous repair following chromosomal breakage in karyotypically balanced germline rearrangements and transgenic integration. Nat Genet. 2012 Mar 4;44(4):390–7, S1. doi: 10.1038/ng.2202.

Shwed PS, Crosthwait J, Douglas GR, Seligy VL. Characterisation of Muta™Mouse λgt10-lacZ transgene: evidence for in vivo rearrangements. Mutagenesis. 2010 Nov;25(6):609–16. doi: 10.1093/mutage/geq048. Epub 2010 Aug 19.

Suzuki O, Hata T, Takekawa N, Koura M, Takano K, Yamamoto Y, Noguchi Y, Uchio-Yamada K, Matsuda J. Transgene insertion pattern analysis using genomic walking in a transgenic mouse line. Exp Anim. 2006 Jan;55(1):65–9.

Licence S, Persson C, Mundt C, Mårtensson IL. The VpreB1 enhancer drives developmental stage-specific gene expression in vivo. Eur J Immunol. 2003 Apr;33(4):1117–26.

Suemizu H, Muguruma K, Maruyama C, Tomisawa M, Kimura M, Hioki K, Shimozawa N, Ohnishi Y, Tamaoki N, Nomura T. Transgene stability and features of rasH2 mice as an Embryo model for short-term carcinogenicity testing. Mol Carcinog. 2002 May;34(1):1–9.

Tacken PJ, van der Zee A, Beumer TL, Florijn RJ, Gijpels MJ, Havekes LM, Frants RR, van Dijk KW, Hofker MH. Effective generation of very low density lipoprotein receptor transgenic mice by overlapping genomic DNA fragments: high testis expression and disturbed spermatogenesis. Transgenic Res. 2001 Jun;10(3):211–21.

Koetsier PA, Mangel L, Schmitz B, Doerfler W. Stability of transgene methylation patterns in mice: position effects, strain specificity and cellular mosaicism. Transgenic Res. 1996 Jul;5(4):235–44.

McFarlane M, Wilson JB. A model for the mechanism of precise integration of a microinjected transgene. Transgenic Res. 1996 May;5(3):171–7.

Pawlik KM, Sun CW, Higgins NP, Townes TM. End joining of genomic DNA and transgene DNA in fertilized mouse eggs. Gene. 1995 Nov 20;165(2):173–81.

Hamada T, Sasaki H, Seki R, Sakaki Y. Mechanism of chromosomal integration of transgenes in microinjected mouse eggs: sequence analysis of genome-transgene and transgene-transgene junctions at two loci. Gene. 1993 Jun 30;128(2):197–202.

Mark WH, Signorelli K, Blum M, Kwee L, Lacy E. Genomic structure of the locus associated with an insertional mutation in line 4 transgenic mice. Genomics. 1992 May;13(1):159–66.

Burdon T, Sankaran L, Wall RJ, Spencer M, Hennighausen L. Expression of a whey acidic protein transgene during mammary development. Evidence for different mechanisms of regulation during pregnancy and lactation. J Biol Chem. 1991 Apr 15;266(11):6909–14.

Radice G, Lee JJ, Costantini F. H beta 58, an insertional mutation affecting early postimplantation development of the mouse embryo. Development. 1991 Mar;111(3):801–11.

Rohan RM, King D, Frels WI. Direct sequencing of PCR-amplified junction fragments from tandemly repeated transgenes. Nucleic Acids Res. 1990 Oct 25;18(20):6089–95.

Wilkie TM, Palmiter RD. Analysis of the integrant in MyK-103 transgenic mice in which males fail to transmit the integrant. Mol Cell Biol. 1987 May;7(5):1646–55.

Covarrubias L, Nishida Y, Mintz B. Early postimplantation embryo lethality due to DNA rearrangements in a transgenic mouse strain. Proc Natl Acad Sci U S A. 1986 Aug;83(16):6020–4.

Wagner EF, Covarrubias L, Stewart TA, Mintz B. Prenatal lethalities in mice homozygous for human growth hormone gene sequences integrated in the germ line. Cell. 1983 Dec;35(3 Pt 2):647–55.

Khillan JS, Overbeek PA, Westphal H. Drosophila P element integration in the mouse. Dev Biol. 1985 May;109(1):247–50.

van der Putten H, Botteri F, Illmensee K. Developmental fate of a human insulin gene in a transgenic mouse. Mol Gen Genet. 1984;198(2):128–38.

Brinster RL, Ritchie KA, Hammer RE, O’Brien RL, Arp B, Storb U. Expression of a microinjected immunoglobulin gene in the spleen of transgenic mice. Nature. 1983 Nov 24-30;306(5941):332–6.

McKnight GS, Hammer RE, Kuenzel EA, Brinster RL. Expression of the chicken transferrin gene in transgenic mice. Cell. 1983 Sep;34(2):335–41.

Palmiter RD, Chen HY, Brinster RL. Differential regulation of metallothionein-thymidine kinase fusion genes in transgenic mice and their offspring. Cell. 1982 Jun;29(2):701–10.

Palmiter RD, Brinster RL, Hammer RE, Trumbauer ME, Rosenfeld MG, Birnberg NC, Evans RM. Dramatic growth of mice that develop from eggs microinjected with metallothionein-growth hormone fusion genes. Nature. 1982 Dec 16;300(5893):611–5.

